# Maternal high fat diet alters lactation-specific miRNA expression and programs the DNA methylome in the amygdala of female offspring

**DOI:** 10.1101/2020.08.13.249300

**Authors:** Sameera Abuaish, Sanoji Wijenayake, Wilfred C. de Vega, Christine M.W. Lum, Aya Sasaki, Patrick O. McGowan

## Abstract

Adverse maternal diets high in saturated fats are associated with impaired neurodevelopment and epigenetic modifications in offspring. Maternal milk, the primary source of early life nutrition in mammals, contains lactation-specific microRNAs (miRNAs). Lactation-specific miRNAs have been found in various offspring tissues in early life, including the brain. We examined the effects of maternal high saturated fat diet (mHFD) on lactation-specific miRNAs that inhibit DNA methyltransferases (DNMTs), enzymes that catalyze DNA methylation modifications, in the amygdala of female offspring during early life and adulthood. Offspring exposed to mHFD showed reduced miR-148/152 and miR-21 transcripts in stomach milk and amygdala in the first week of life. This was associated with increased DNMT1 expression, DNMT activity, and global DNA methylation in the amygdala. In addition, persistent DNA methylation modifications from early life to adulthood were observed in pathways involved in neurodevelopment as well as genes regulating the DNMT machinery and protein function in mHFD offspring. The findings indicate a novel link between exogenous, lactation-specific miRNAs and developmental programming of the neural DNA methylome in offspring.

## Introduction

Adverse maternal diets high in saturated fats (mHFD) are associated with impaired neurodevelopment and epigenetic modifications in offspring. Methylation modifications have been reported both globally (Vucetic et al., 2010) and at select candidate genes in the brain of offspring exposed to maternal nutrition stress during perinatal life (Grissom et al., 2014; Marco et al., 2014; Schellong et al., 2019). However, studies to date have largely focused on adult offspring, long after the period of early developmental exposure. Consequently, the gene regulatory mechanisms involved in DNA methylation modifications during the period of exposure to a maternal obesogenic diet and the extent to which these modifications are maintained into adulthood are unknown.

There is recent evidence that the expression of maternal, lactation-specific microRNAs (miRNAs), which mediate gene silencing via post-transcriptional regulation of target mRNAs, is significantly altered by a maternal obesogenic diet (Chen et al., 2017). Lactation-specific miRNAs, encapsulated in stable milk-derived exosomes, appear to cross the offspring’s developing intestinal endothelium post-ingestion (Modepalli et al., 2014; Zempleni et al., 2019; Zhang et al., 2012), and have been found in various tissues in offspring during early life, including the brain (Baier et al., 2014; Chen et al., 2016; Izumi et al., 2015; Lässer et al., 2011; Manca et al., 2018). miRNAs belonging to miR-148/152 family are of particular interest, because they are highly expressed in maternal milk (Benmoussa and Provost, 2019; Van Herwijnen et al., 2018) and are known regulators of DNA methyltransferases (DNMTs), enzymes that catalyze DNA methylation modifications. *In vitro* studies have shown that miRNA-148a and miRNA-152 directly target and inhibit DNMT1 translation (Long et al., 2014; Pan et al., 2010; Wang et al., 2014; Xu et al., 2013). miRNA-21 is also abundant in maternal milk and indirectly inhibits DNMT1 translation by targeting Ras guanyl nucleotide-releasing protein-1 (RASGRP1) (Pan et al., 2010).

Here, we examined the relationship between lactation-specific miRNAs, the enzymatic machinery responsible for DNA methylation modifications, and programming of the DNA methylome in female Long-Evans rat offspring exposed to mHFD. We found that exposure to mHFD altered levels of lactation-specific miRNA in stomach milk and the brain (amygdala) during the first week of life. These changes in miRNA levels were inversely associated with DNMT transcriptional and enzymatic activity. Correspondingly, mHFD offspring showed lower levels of global DNA methylation and locus-specific DNA methylation modifications, some of which appear to persist to adulthood.

## Results

### Dam and Offspring Body Weight

mHFD dams consuming HFD for 4 weeks were significantly heavier than mCHD dams at conception (t(1,14) = 2.91, p=0.01). During gestation, the caloric intake was significantly higher among mHFD dams compared to mCHD dams (t(1,14) = 3.29, p=0.005). There were no differences in offspring body weight across the two diet groups at birth (p>0.05) and all pups increased in weight during the pre-weaning period (F(1,13)= 953.7, p<0.01). At postnatal day 7 (P7), offspring exposed to mHFD (18 ± 0.72 g) were heavier relative to mCHD offspring (16 ± 0.38 g; Bonferroni post-hoc p=0.006). In data reported previously, body weight and caloric intake for dams and their offspring sacrificed in adulthood for this study, showed similar effects of mHFD during the pre-weaning period (Sasaki et al., 2013). At P90, offspring showed comparable body weights among the two diet groups (p>0.05).

### Expression of maternal milk-derived miRNAs in stomach milk and amygdala

At P7, miR-152-5P (t(1,6)=6.393, p=0.001) and miR-21-5P (t(1,6)=4.15, p=0.006) showed lower transcript abundance in the stomach milk of female neonates exposed to mHFD, whereas expression levels of miR-148-5P (t(1,6)=−0.056, p=0.957), miR-148-3P (t(1,6)=−1.751, p=0.131), and miR-152-3P (t(1,6)=−0.042, p=0.968) remained unchanged (Fig.1A). For the same five miRNAs measured in the female amygdala, miR-148-3P (t(1,12)=2.293, p=0.041) and miR-152-3P (t(1,12)=2.254, p=0.048) decreased in transcript abundance in response to mHFD when compared to mCHD (Fig. 1B). miR-148-5P (t(1,12)=0.911, p=0.380) and miR-152-5P (t(1,12)=0.049, p=0.962) remained unchanged and miR-21-5P was not detected in the amygdala of female offspring at P7.

**Fig 1.**
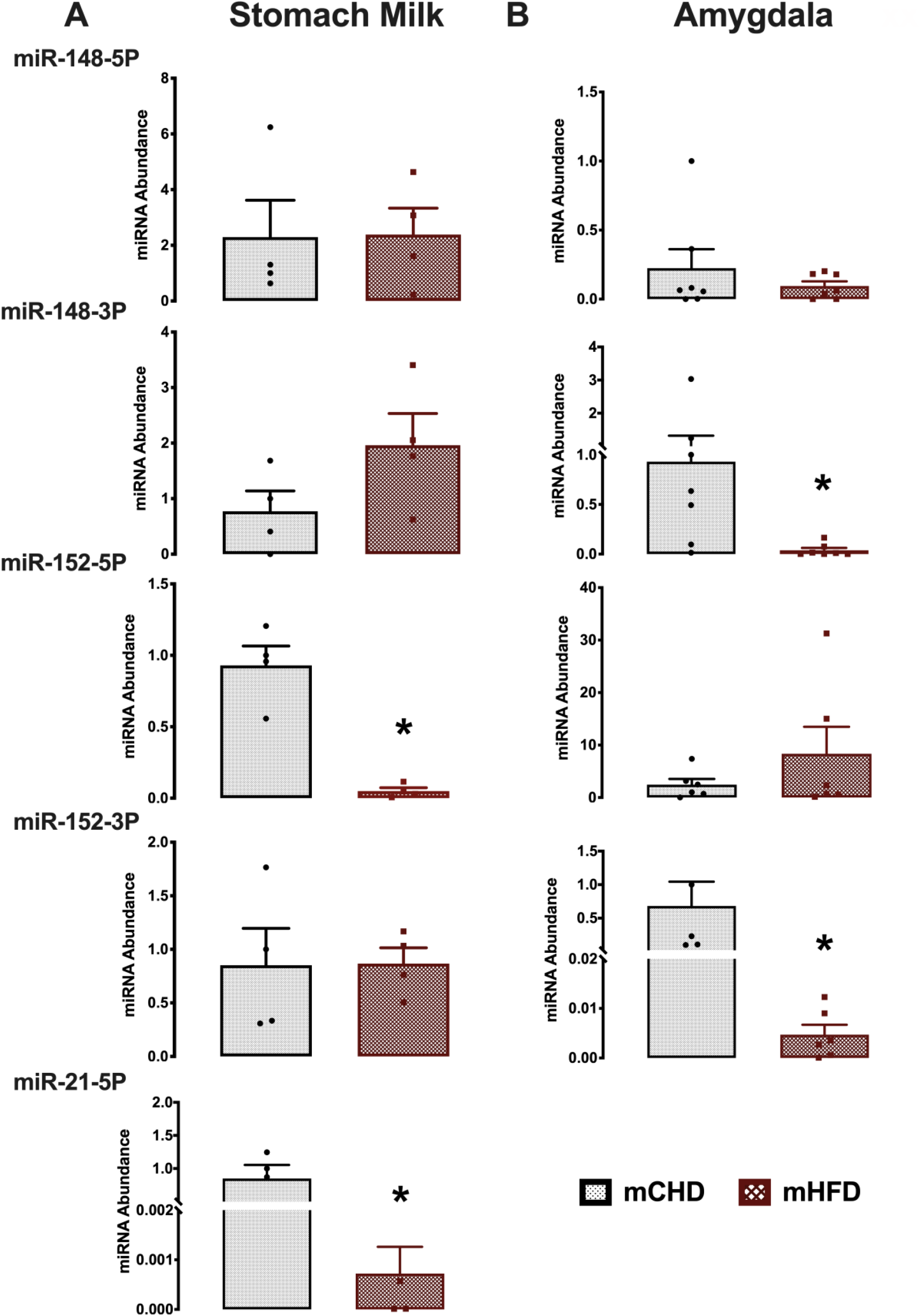
Relative abundance of five lactation-specific miRNAs with maternal HFD exposure (mHFD) compared to chow diet control (mCHD). (A) Stomach milk and (B) amygdala of female offspring at P7. Data are mean ± SEM with n = 4 and n=6 independent biological replicates per experimental group in stomach milk and amygdala, respectively. * Significantly different from mCHD (p < 0.05; two-tailed student t-test).

### mRNA expression of epigenetic regulators

We examined the mRNA transcript abundance of epigenetic regulators involved in DNA methylation modifications, including DNMT1, DNMT3a, DNMT3b, direct post-transcriptional targets of lactation-specific miR-148/152 and miR-21, as well as MeCP2 and GADD45α in the amygdala of female offspring exposed to mCHD and mHFD at P7 and P90. DNMT1 (t(1,10)=−2.346, p=0.041) and MeCP2 (t(1,10)=−2.237, p=0.049) transcript abundance significantly increased in response to mHFD at P7 (Fig. 2A). Transcript abundance of DNMT3a (t(1,12)=−0.942, p=0.365), DNMT3b (t(1,12)=−0.712, p=0.490), and GADD45α (t(1,12)=−0.871, p=0.401) remained unchanged. At P90, transcript abundance of DNMT1 (t(1,8)=−1.20, p=0.265), DNMT3a (t(1,8)=1.482, p=0.177), DNMT3b (t(1,8)=0.623, p=0.551), MeCP2 (t(1,8)=0.724, p=0.490), and GADD45α (t(1,8)=1.186, p=0.270), remained unchanged (Fig. 2B).

**Fig 2.**
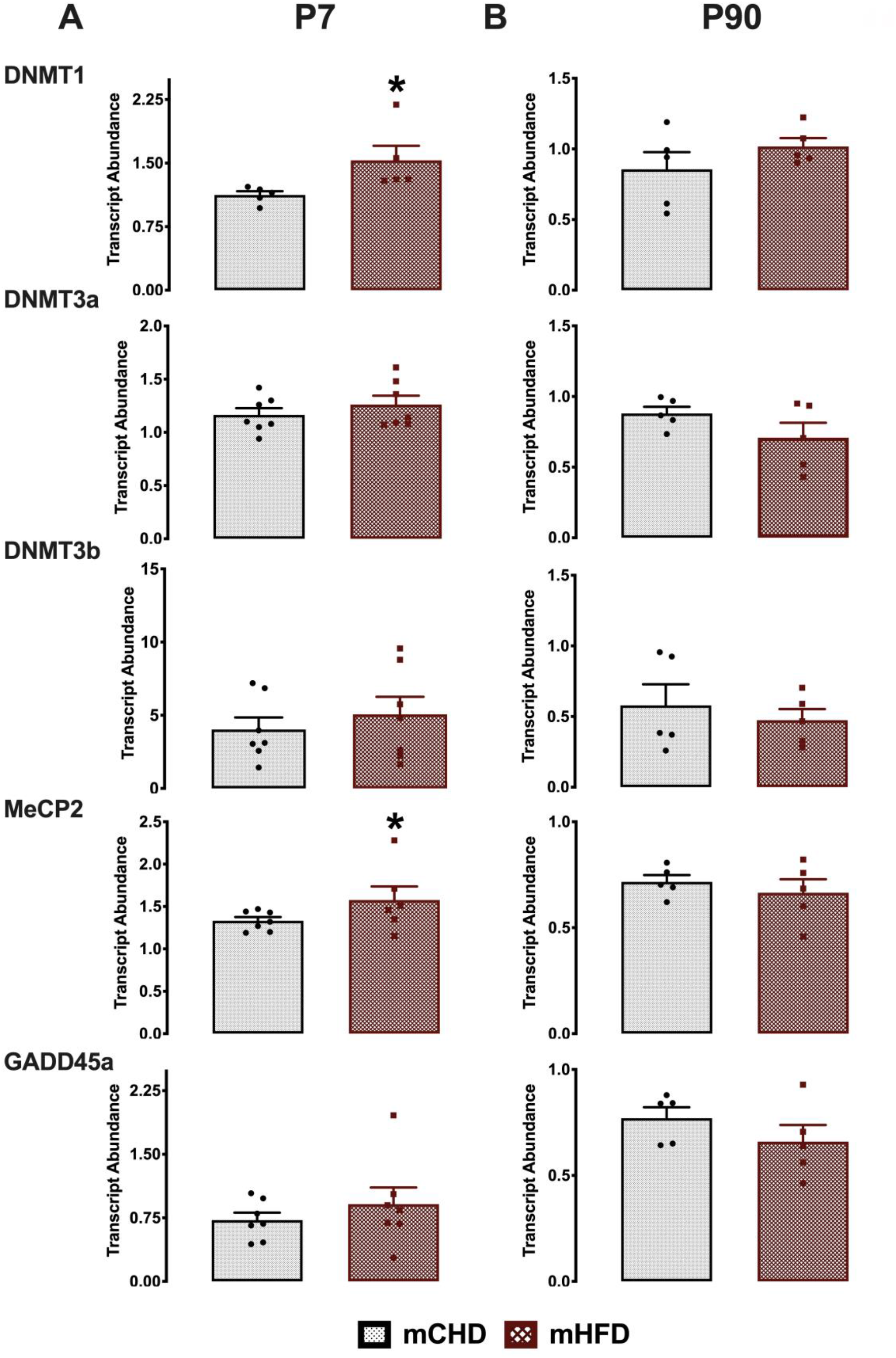
Effect of maternal HFD exposure on relative transcript abundance of DNA methyltransferases and DNA binding proteins in the amygdala of female offspring. (A) P7 and (B) P90. mCHD is control chow diet and mHFD is maternal high-fat diet exposure. Data are mean ± SEM with n = 6 independent biological replicates per experimental group. * Significantly different from mCHD (p < 0.05; two-tailed student t-test).

### DNMT Enzymatic Activity and Global DNA Methylation

We next examined whether changes in transcript abundance of DNMTs was associated with changes in their enzymatic activity and global LINE-1 5-methylcytosine (5mC) levels. In P7 offspring, DNMT enzymatic activity robustly increased in response to mHFD (t(1,5)=−5.95, p=0.02; Fig. 3A). Correspondingly, global LINE-1 5mC (%) increased in response to mHFD when compared to mCHD (t(1,6)=−2.98, p=0.03; Fig. 3B). A strong linear correlation was observed between increased DNMT enzymatic activity (OD/h/mg) and increased global LINE-1 5mC (%) levels at P7 (R_2_ = 0.896, p= 0.001; Fig. 3C).

**Fig 3.**
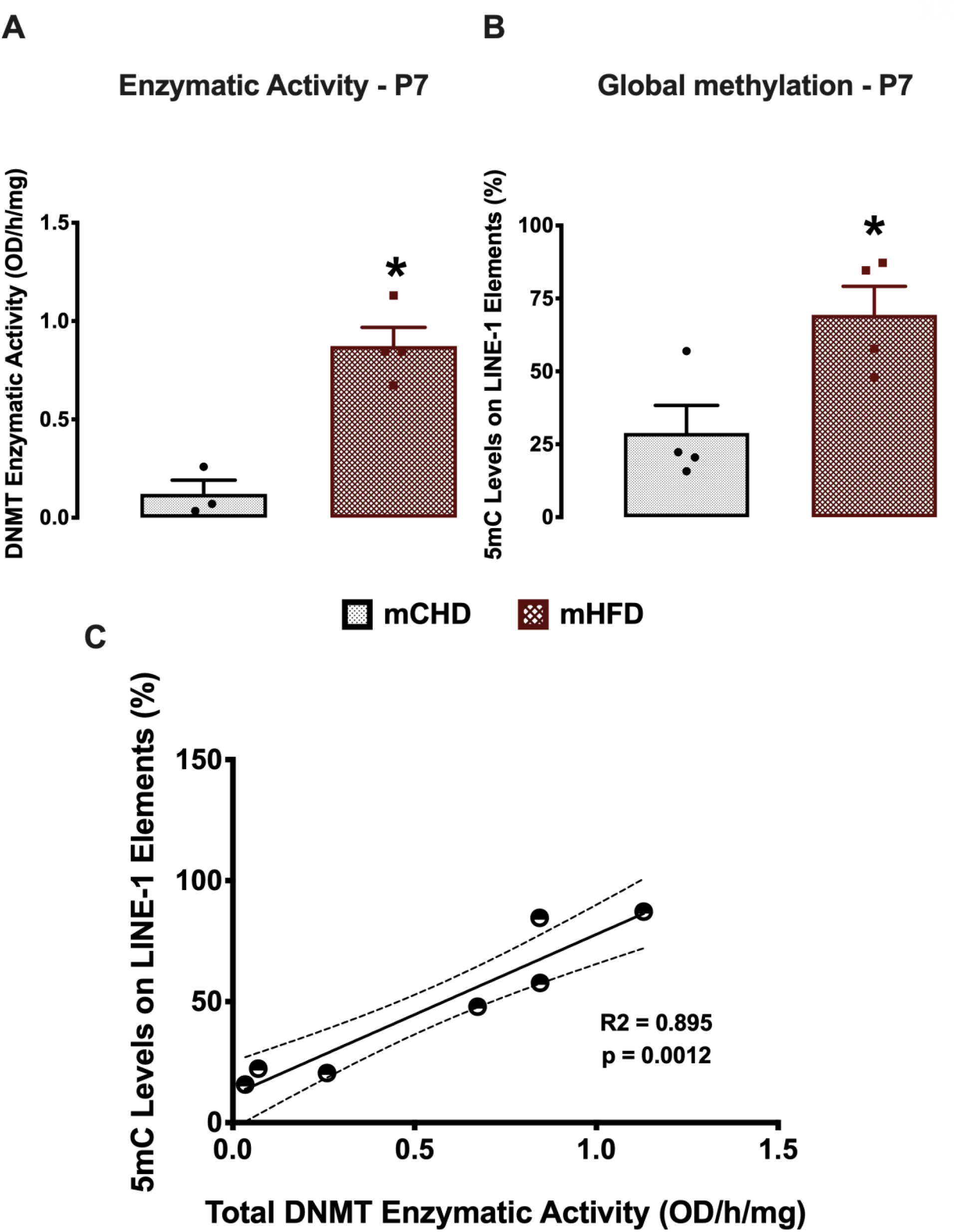
Global DNA methylation in the amygdala of neonatal offspring. A) Total DNA methyltransferase activity (OD/h/mg). B) LINE-1 global 5mC levels (%). C) Pearson correlation between total DNMT enzymatic activity (OD/h/mg) and global 5mC (%) levels (p<0.05). mCHD is control house chow and mHFD is maternal high-fat diet exposure. Data are mean ± SEM with n = 3-4 independent biological replicates per experimental group. * Significantly different from mCHD (p < 0.05; two-tailed student t-test).

At P90, female mHFD offspring had a trend of lower DNMT enzymatic activity compared to mCHD (t(1,6)= 1.794, p = 0.06; Fig. 4A). Global LINE-1 5mC (%) levels significantly decreased in response to mHFD (t(1,6)=3.805, p=0.009; Fig. 4B). However, global LINE-1 5mC (%) was not associated with total DNMT enzymatic activity at P90 (R_2_ = 0.07, p= 0.538; Fig. 4C).

**Fig 4.**
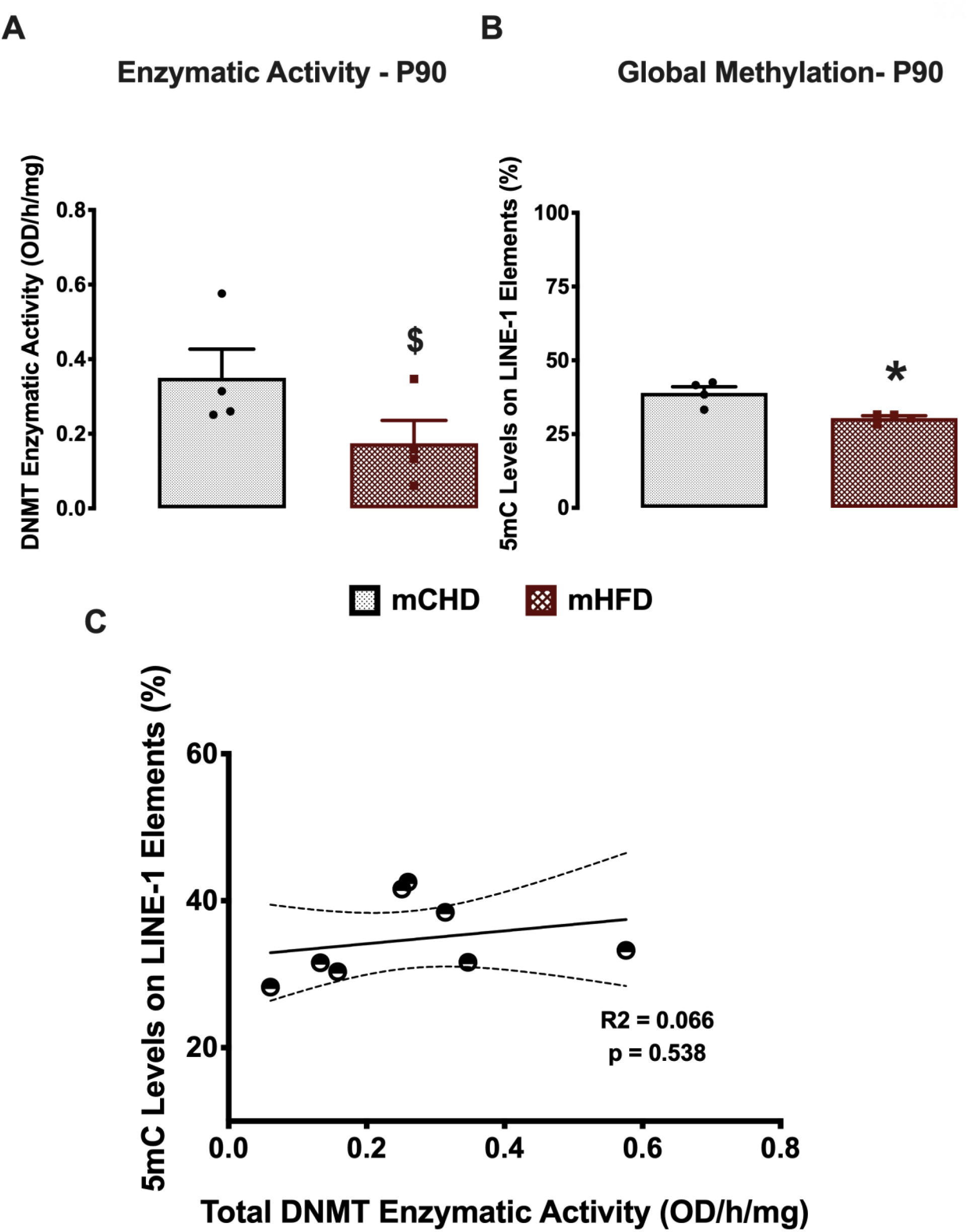
Global DNA methylation in the amygdala of adult offspring. A) Total DNA methyltransferase activity (measured as OD/h/mg). B) LINE-1 global 5mC levels (%). C) Pearson linear correlation between total DNMT activity (OD/h/mg) and global 5mC (%) levels (p<0.05). mCHD is control house chow and mHFD is maternal high-fat diet exposure. Data are mean ± SEM with n = 4 independent biological replicates per experimental group. * Significantly different from mCHD (p < 0.05). $ Trending towards significance (p=0.06).

### Genome-wide DNA Methylation Analysis

Genome-wide DNA methylation at single nucleotide resolution was assessed using reduced representation bisulfite sequencing (RRBS). At P7 and P90, female offspring showed 777 and 1050 significant differentially methylated regions (DMRs), respectively. At P7, 69% of DMRs were hypomethylated and 31% were hypermethylated, and at P90 61% of DMRs were hypomethylated and 39% were hypermethylated (Fig. 5A). At P7, approximately 50% of DMRs were found in intergenic regions, while 21 % and 29 % were found in promoters and gene bodies, respectively. At P90, 56% of DMRs were found in intergenic regions, while only 6% were found in promoters and 38% were found in gene bodies.

**Fig 5.**
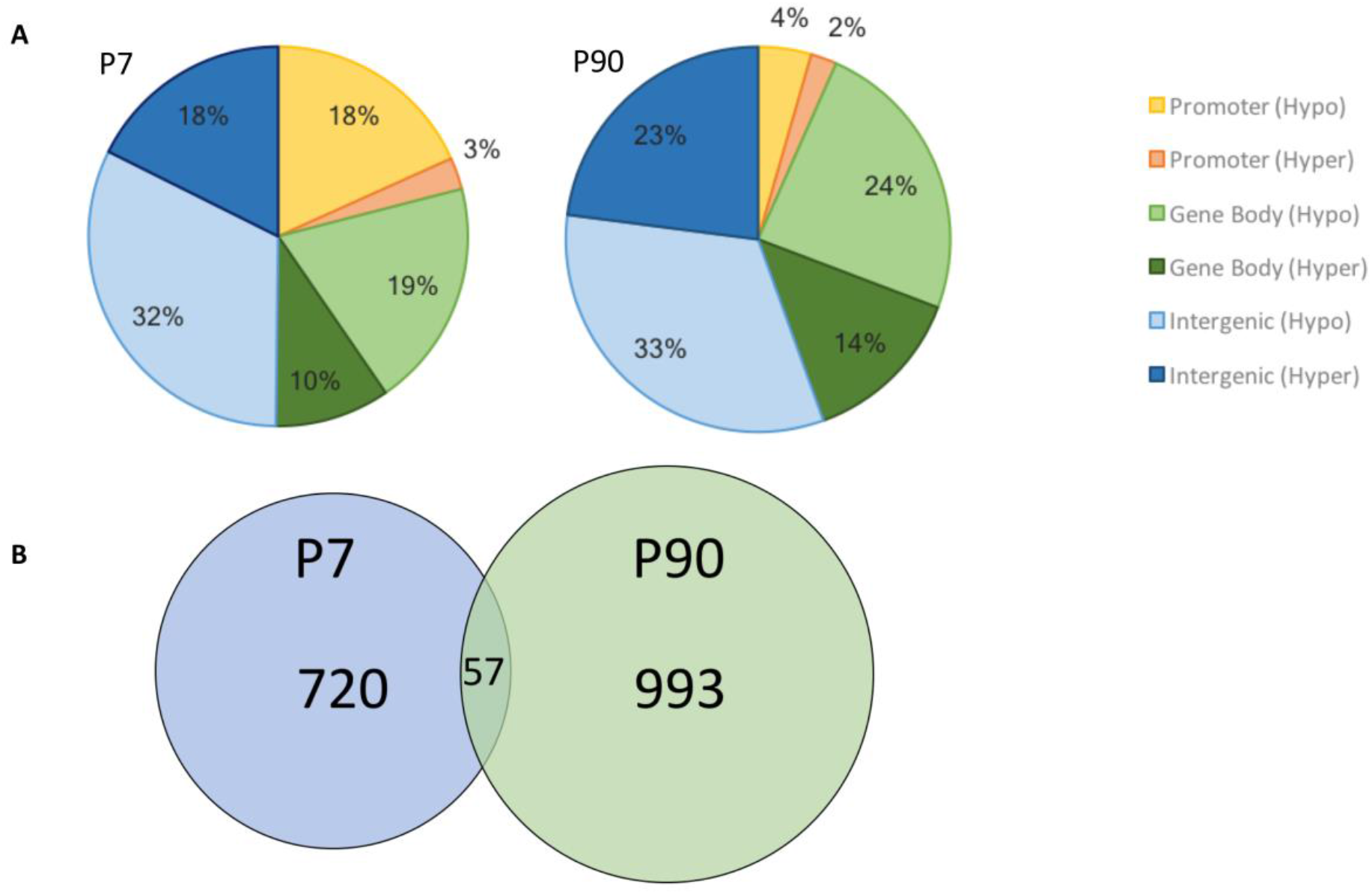
Significant differentially methylated regions (DMRs) in the amygdala of female offspring exposed to maternal HFD at P7 and P90. A) Distribution of hyper- and hypomethylated DMRs in HFD offspring in terms of genic location including promoter regions, gene body, and intergenic regions. B) Venn diagram illustrating the number of DMRs that are unique to female offspring at P7 (blue) and P90 (green) in response to maternal HFD exposure in the amygdala. The area in between represents the numbers of overlapping DMRs across the two ages.

Fifty-seven DMRs were shared across the two age groups (Fig. 5B) corresponding to 26 genes (Table 1). Fifteen of these genes were consistently hypomethylated at both P7 and P90 (LOC310926, Tbc1d10b, Paqr6, Rgs3, Unc79, Icam4, Gtf2e1, Hic2, Tpst2, Rab5a, Nfatc1, Nhs, Lancl3, Mecp2, Ndufb1), while five genes remained hypermethylated at the two ages (LOC499742, Anxa6, Plvap, Rsu1, Zfp423). Seven DMRs that were hypomethylated and 12 DMRs that were hypermethylated in response to mHFD at P7 (33% of the overlapping DMRs) showed opposite methylation differences at P90. The 6 genes associated with these DMRs showed opposite methylation patterns in early life compared to adulthood. Three genes were hypomethylated at P7 and hypermethylated at P90 (Polr3g, Prpf38b, and Htatsf1) and another 3 genes were hypermethylated at P7 and hypomethylated at P90 (Ank1, Dupd1, and Tmprss9). A total of 372 and 444 genes were found to be differentially methylated at P7 and P90, respectively (Supplementary Table 2-3).

**Table 1.** Genes containing significant DMRs sorted by chromosomes, percentage of differential methylation between mCHD and mHFD, and the location of the DMR in the amygdala of female offspring at P7 and P90.

| Chromosome | Gene Symbol | Gene Name | P7 Methylation differences (%) | P90 Methylation differences (%) | Gene Location |
| --- | --- | --- | --- | --- | --- |
| chr1 | LOC310926 | Hypothetical protein LOC310926 | -21.01 | -10.295 | Gene body |
| chr1 | Tbc1d10b | TBC1 domain family, member 10b | -11.427 | -30.145 | Promoter |
| chr2 | Paqr6 | Progesterone and adiponectin receptor family member 6 | -40.93 | -11.022 | Gene body |
| chr2 | Polr3g | RNA polymerase III subunit G | -19.942 | 5.273 | Gene body |
| chr2 | Prpf38b | Pre-mRNA processing factor 38B | -8.727 | 5.153 | Gene body |
| chr3 | LOC499742 | LRRG00137 | 27 | 34.943 | Gene body |
| chr5 | Rgs3 | Regulator of G-protein signaling 3 | -8.432 | -7.377 | Gene body |
| chr6 | Unc79 | Unc-79 homolog | -17.035 | -13.765 | Gene body |
| chr7 | Tmprss9 | Transmembrane protease, serine 9 | 12.28 | -10.145 | Gene body |
| chr8 | Icam4 | Intercellular adhesion molecule 4, Landsteiner-Wiener blood group | -12.475 | -15.715 | Promoter |
| chr10 | Anxa6 | Annexin A6 | 20.15 | 34.822 | Gene body |
| chr11 | Gtf2e1 | General transcription factor IIE subunit 1 | -10.285 | -6.048 | Gene body |
| chr11 | Hic2 | HIC ZBTB transcriptional repressor 2 | -7.472 | -5.325 | Gene body |
| chr12 | Tpst2 | Tyrosylprotein sulfotransferase 2 | -8.54 | -15.268 | Gene body |
| chr14 | Rab5a | Ras-related protein Rab-5A | -7.45 | -7.975 | Promoter |
| chr15 | Dupd1 | Dual specificity phosphatase and pro isomerase domain containing 1 | 11.21 | -8.108 | Gene body |
| chr16 | Ank1 | Ankyrin 1 | 5.695 | -30.753 | Gene body |
| chr16 | Plvap | Plasmalemma vesicle associated protein | 7.403 | 5.117 | Promoter |
| chr17 | Rsu1 | Ras suppressor protein 1 | 5.56 | 7.537 | Gene body |
| chr18 | Nfatc1 | Nuclear factor of activated T-cells 1 | -9 | -17.6 | Gene body |
| chr19 | Zfp423 | Zinc finger protein 423 | 9.005 | 8.777 | Gene body |
| chrX | Nhs | NHS actin remodeling regulator | -5.795 | -18.252 | Gene body |
| chrX | Lancl3 | LanC like 3 | -14.728 | -11.385 | Promoter |
| chrX | Mecp2 | Methyl CpG binding protein 2 | -5.84 | -5.72 | Promoter |
| chrX | Htatsf1 | HIV-1 Tat specific factor 1 | -5.308 | 14.31 | Promoter |
| chrX | Ndufb1<br>1 | NADH:ubiquinone<br>oxidoreductase<br>subunit B11 | -5.608 | -17.19 | Promoter |

### Gene Annotation Enrichment Analysis

gProfiler was used to conduct Gene Ontology (GO) analysis to identify enriched biological processes associated with the differentially methylated genes. Thirty-four significantly enriched overlapping GO terms were shared across early life and adulthood, clustered into 5 functionally related groups (Fig. 6; Supplementary Table 4). P7 and P90 animals shared GO terms associated with cellular morphogenesis and organism development (10 GO terms) as well as neuronal projection and nervous system development (11 terms). In addition, terms associated with protein phosphorylation (5 GO terms) and protein transportation and secretion (8 GO terms) were shared across P7 and P90. The lipid response GO term was uniquely shared across both age groups and did not cluster with other GO terms.

**Fig 6.**
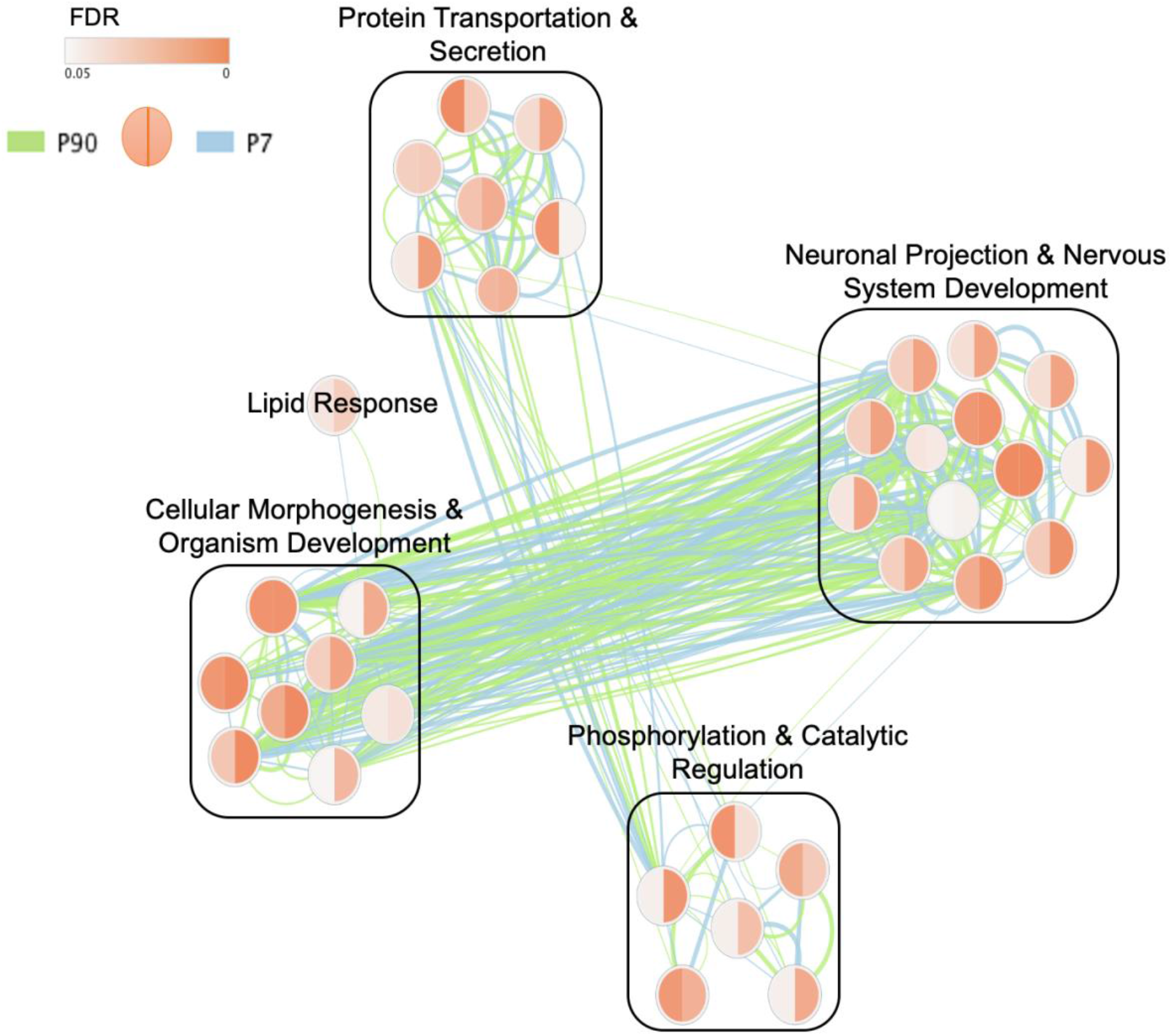
Overlapping clusters of Gene Ontology (GO) Biological Processes enriched in P7 (right half of nodes and blue edges) and P90 (left half of nodes and green edge) differentially methylated gene sets for female offspring exposed to maternal HFD. The size of the nodes represents the number of genes, while the color indicates the FDR p-value. The edges between the nodes indicate shared genes, with edge thickness representing the number of genes in common.

At P7, a total of 372 DMRs were associated with annotated genes. gProfiler identified 163 significantly enriched biological processes that clustered into 10 functionally related groups (Fig. 7; Supplementary Table 5). Three groups were involved in systems development, consisting of processes related to organismal (8 GO terms), cellular (19 GO terms), and neuronal (27 GO terms) development. Two other groups were involved in post-translational modifications, including protein phosphorylation (13 GO terms) and catabolic process (29 GO terms). GO terms involved in cellular responses to stimuli (13 GO terms), including growth factors, stress, and organic compounds, as well as GO terms involving cell secretion and signal transduction (23 GO terms), metabolic regulation (11 GO terms), apoptosis (7 GO terms), and gene expression (13 GO terms) were also enriched in early life (FDRs<0.05).

**Fig 7.**
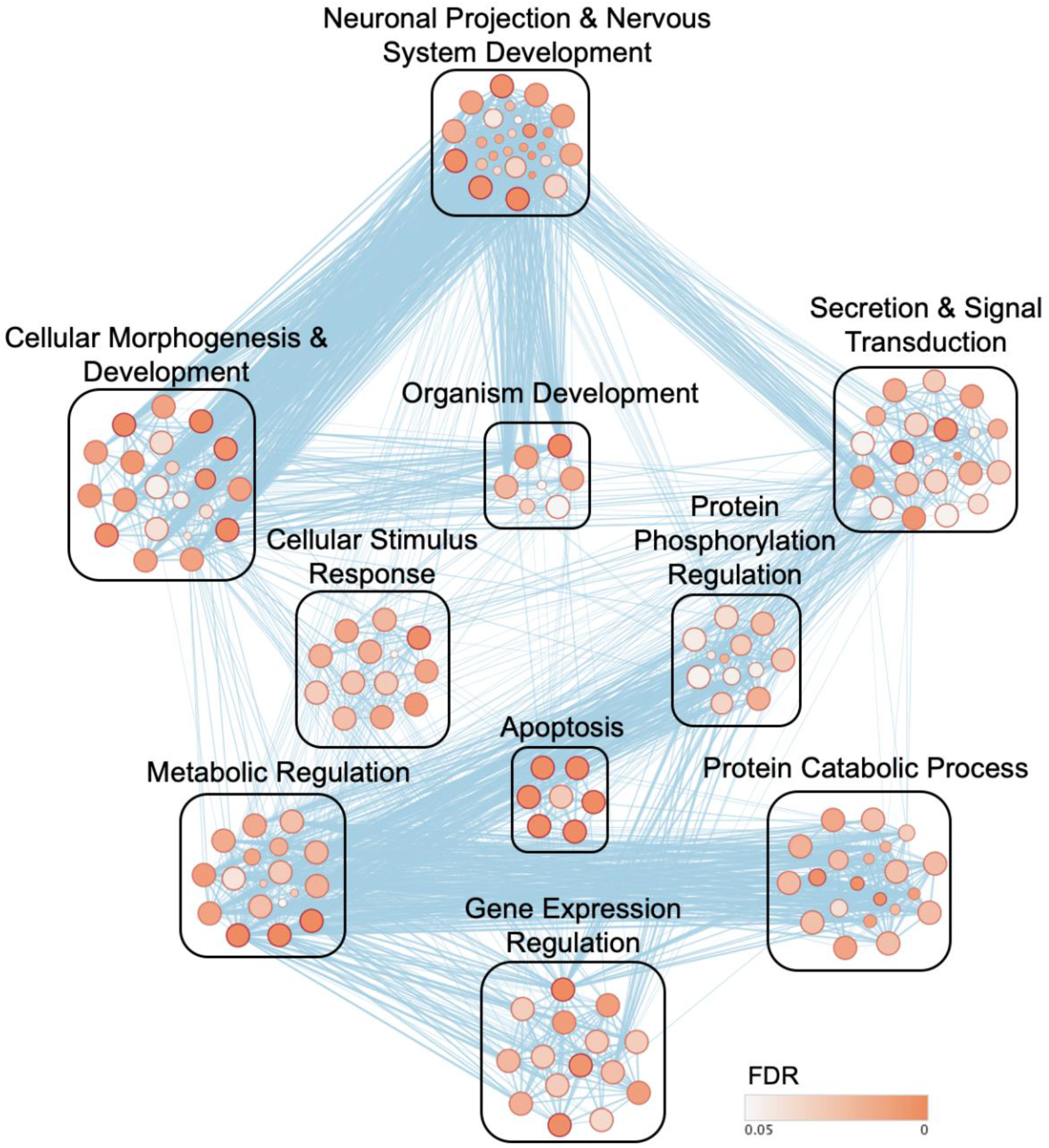
gProfiler clusters of Gene Ontology (GO) Biological Process terms significantly enriched at P7 in female offspring in response to maternal HFD exposure in the amygdala. The size of the nodes represents the number of genes, while the color indicates the FDR p-value. The edges between the nodes indicate shared genes, with edge thickness representing the number of genes in common.

At P90, a total of 444 DMRs were associated with annotated genes, where 105 GO terms were clustered into 7 functionally related groups (Fig. 8; Supplementary Table 6). Three groups were involved in systems development, including organismal (12 GO terms), cellular (14 GO terms), and neuronal (19 GO terms) development. Twenty-four GO terms were clustered into protein phosphorylation and signaling pathways, which included GTPase activity, regulation, and intracellular signal transduction. Twenty-one GO terms were clustered into protein localization and secretion and 10 GO terms were clustered into ion transport. Terms involved in cellular stimulus response (5 GO terms) and the response to lipopolysaccharide and lipids were also enriched in adulthood (FDRs<0.05). DAVID analysis generated a similar list of enriched GO terms that were shared across early life and adulthood as well as GO terms that are unique to each age (Supplementary Fig. 1-3).

**Fig 8.**
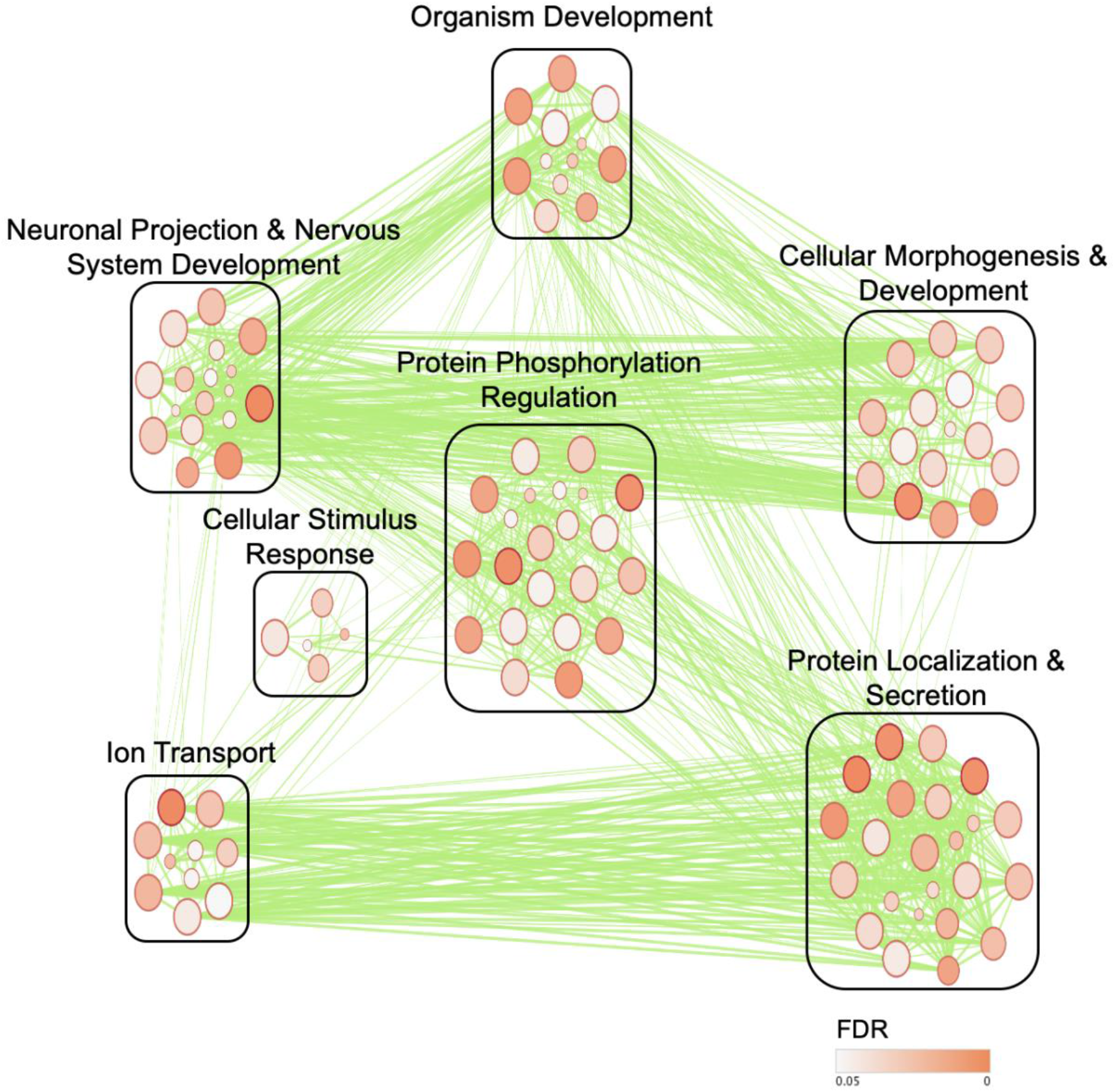
gProfiler clusters of Gene Ontology (GO) Biological Processes terms significantly enriched at P90 in female offspring in response to maternal HFD exposure in the amygdala. The size of the nodes represents the number of genes, while the color indicates the FDR p-value. The edges between the nodes indicate shared genes with edge thickness representing the number of genes in common.

Gene enrichment analysis was conducted using the Kyoto Encyclopedia of Genes and Genomes (KEGG) database of pathways representing both empirical and predicted molecular interactions, to examine networks that include cellular processes, organismal systems, environmental information processing, and metabolism. Eight KEGG pathways were shared across P7 and P90 age groups, including MAPK, cGMP-PKG, cAMP, and calcium signaling pathways (FDR<0.05; Table 2). These signaling pathways are involved in axon guidance, which was also enriched with a large number of differentially methylated genes. In addition, the oxytocin-signaling pathway, tight junction, and circadian entrainment KEGG pathways were significantly enriched from early life to adulthood.

**Table 2.** Common KEGG pathways enriched by the genes identified from DMR analysis at P7 and P90 in response to maternal HFD exposure in female amygdala along with their FDR values and list of enriched genes.

| ID | Description | P7 FDR | P90 FDR | P7 Genes | P90 Genes |
| --- | --- | --- | --- | --- | --- |
| KEGG<br>:04010 | MAPK<br>signaling<br>pathway | 4.98E-03 | 4.47E-02 | MAPK10,RPS6KA<br>6,MAP3K14,RPS6<br>KA3,CACNA1C,C<br>ACNA1F,RASGR<br>F1,NFATC1,<br>MAP3K4,DUSP9,<br>FGF16 | CACNA1C,MAP<br>3K6,MECOM,CA<br>CNA1D,NFATC1<br>,NTRK2,NFKB1,<br>CACNA2D3,<br>FGF13 |
| KEGG<br>:04020 | Calcium<br>signaling<br>pathway | 5.30E-03 | 2.93E-03 | PLCD3,CACNA1C<br>,SLC8A1,CACNA<br>1F,SPHK1,HTR2C<br>,PLCB4,<br>CAMK2B,ATP2B3 | NOS1,P2RX7,M<br>YLK,ITPKB,PH<br>KA1,ADCY8,CA<br>CNA1C,EDNRB,<br>CACNA1D,<br>SLC25A5 |
| KEGG<br>:04022 | cGMP-PKG<br>signaling<br>pathway | 8.64E-03 | 5.38E-03 | CACNA1C,SLC8A<br>1,CACNA1F,NFA<br>TC2,NFATC1,MR<br>VI1,PLCB4,ATP2<br>B3 | ATF6B,MYLK,A<br>DCY8,CACNA1<br>C,EDNRB,CACN<br>A1D,NFATC1,G<br>UCY1A2,<br>SLC25A5 |
| KEGG<br>:04024 | cAMP<br>signaling<br>pathway | 2.73E-03 | 1.21E-02 | MAPK10,CACNA<br>1C,GRIA3,ABCC4<br>,CACNA1F,GRIN3<br>B,NFATC1,TIAM1<br>,CAMK2B, | ADCY8,CACNA<br>1C,GABBR2,CA<br>CNA1D,NFATC1<br>,VAV3,TIAM1,N<br>FKB1, |

|  |  |  |  | ATP2B3 | RAPGEF3 |
| --- | --- | --- | --- | --- | --- |
| KEGG<br>:04360 | Axon<br>guidance | 6.80E-04 | 9.09E-03 | FYN,NTN1,PAK3,<br>EFNB1,NFATC2,R<br>GS3,SLIT1,SEMA<br>6B,CAMK2B,BMP<br>7,PLXNA3 | FYN,EPHB1,DP<br>YSL5,EPHB2,LI<br>MK2,SEMA4A,R<br>GS3,PARD3,BM<br>P7 |
| KEGG<br>:04530 | Tight<br>junction | 2.69E-02 | 1.52E-03 | RUNX1,MAPK10,<br>RAP2C,DLG3,TIA<br>M1,ARHGEF18,E<br>PB41L4B | DLG3,MYH10,S<br>LC9A3R1,CLDN<br>11,CACNA1D,TJ<br>P3,TIAM1,MAGI<br>1,ARHGEF18,PA<br>RD3,CGNL1 |
| KEGG<br>:04713 | Circadian<br>entrainment | 3.82E-02 | 1.41E-02 | CACNA1C,GRIA3<br>,PER3,<br>PLCB4,CAMK2B | NOS1,ADCY8,C<br>ACNA1C,CACN<br>A1D,GNG7,GUC<br>Y1A2 |
| KEGG<br>:04921 | Oxytocin<br>signaling<br>pathway | 6.94E-03 | 2.84E-02 | CDKN1A,CACNA<br>1C,CACNA1F,NF<br>ATC2,EEF2K,NFA<br>TC1,PLCB4,CAM<br>K2B | MYLK,ADCY8,<br>CACNA1C,CAC<br>NA1D,NFATC1,<br>GUCY1A2,CAC<br>NA2D3 |

Ten KEGG pathways were unique to early life (FDR<0.05; Supplementary Table 7), including the Wnt signaling pathway, neurotrophin signaling pathway, MAPK signaling, axonal guidance, and long-term potentiation. In addition, two endocrine system pathways including thyroid hormone and GnRH signaling were enriched with genes differentially methylated at P7. KEGG pathways associated with human diseases affecting the nervous system, including Alzheimer’s disease and amphetamine addiction, and nervous system related signaling pathways involved in retrograde endocannabinoid signaling were also identified.

Twenty KEGG pathways were unique to adulthood (FDR<0.05; Supplementary Table 8). A set of three interrelated signaling pathways were enriched at P90, including the Ras, Rap1, and PI3K-Akt signaling pathways. Three nervous system-specific pathways were enriched including, GABAergic synapse, glutamatergic synapse, and long-term depression. The analysis revealed three endocrine-related pathways, including insulin secretion, relaxin signaling, and aldosterone synthesis and secretion. Two pathways important for cellular integrity, extracellular matrix (ECM)-receptor interaction and focal adhesion, were also identified. A number of cardiovascular pathways relating to cardiomyopathy and vascular smooth muscle contraction as well as pathways involved in cancer development and transcriptional dysregulation were enriched at P90.

## Discussion

In this study, we examined the effects of mHFD exposure on lactation-specific miRNAs that inhibit DNMTs, as well as changes in genome-wide DNA methylation modifications in offspring in early life and adulthood. Members of the lactation-specific miR-148/152 family and miR-21 exhibited decreased abundance in ingested stomach milk and the amygdala of mHFD offspring in early life. Correspondingly, we observed increased transcript abundance of DNMT1 and MeCP2 in response to mHFD in neonates. Total DNMT enzymatic activity and global LINE-1-5mC (%) levels also increased in the amygdala in early life, but not in adulthood. Genes regulating the DNMT machinery as well as neurodevelopment were differentially methylated across the two ages.

Female pups exposed to mHFD showed a reduction in miR-152-5P and miR-21-5P in their stomach milk (Fig. 1A) and a reduction in miR-148a-5P and miR-152-3P levels in the amygdala during early life compared to mCHD offspring (Fig. 1B). These findings support previous research indicating that lactation-specific miRNAs belonging to miR-148/152 family survive the digestive tract (likely encapsulated in stable milk-derived exosomes), are absorbed across offspring’s intestinal barrier, and are subsequently transported to target neural tissues in rodents. Indeed, several previous studies have shown that milk-derived exosomes survive degradation in the GI tract (Benmoussa et al., 2016; Izumi et al., 2015, 2012; Zhou et al., 2012) and can be absorbed across the intestinal barrier (Modepalli et al., 2014), especially during early life when gut permeability is high (Gareau, 2011). Milk-derived exosomes have also been shown to successfully cross the blood brain barrier of the recipient and have unique distribution patterns (Chen et al., 2016; Manca et al., 2018). Interestingly, several studies have failed to detect endogenously produced miR-148/152 family in peripheral organs including the liver and heart and in several regions of the CNS, including cerebellum, thalamus, hippocampus, and spinal cord of adult rats using time dependent expression profiles generated from small RNA sequencing and microarray analyses (Izumi et al., 2014; Minami et al., 2014; Smith et al., 2016). Although we cannot definitively rule out endogenous expression of the miRNAs examined in this study, our findings are consistent with evidence that these miRNAs are likely of maternal origin and are transferred via milk during early life (Alsaweed et al., 2016; Manca et al., 2018).

We found a strong inverse correlation between levels of lactation-specific miRNAs and DNMT expression, where miR-148/152 and miR-21 levels decreased, while DNMT1 and MeCP2 levels increased in the amygdala in early life among mHFD offspring (Fig. 2A). These findings concur with other studies showing post-transcriptional regulation of DNMT1 by the miR-148//152 family. For example, increased expression of miR-152 during lactation in dairy cows has been linked to a marked reduction of DNMT1 mRNA and protein expression in mammary glands (Wang et al., 2014). This reduction is associated with a decrease in global DNA methylation in bovine mammary glands. Likewise, a strong inverse relationship was previously shown between miR-148a and miR-21 and DNMT1 expression in bovine maternal epithelial cell culture (Long et al., 2014; Pan et al., 2010; Xu et al., 2013). In human milk, miR-148a has also been shown to downregulate the expression of DNMT1 *in vitro* (Golan-Gerstl et al., 2017). DNMT1 is known to recruit MeCP2 to methylated loci, inducing transcriptional repression (Chen et al., 2015). Interestingly, our data also show evidence of increased expression of MeCP2 (Fig. 2A) that is positively associated with increased DNMT1 levels and inversely associated with miR-148/152 transcript levels. Further, in silico analysis has predicted MeCP2 to be a direct post-transcriptional target of miR-152-3P, however to our knowledge there is no experimental evidence confirming this prediction to date (Ehrhart et al., 2016; www.targetscan.org).

DNMT expression in offspring is known to be responsive to changes in the maternal nutritional environment. For example, maternal dietary protein restriction was associated with a decrease in DNMT1 expression in the liver of adult offspring (Lillycrop et al., 2007), and mHFD consumption was associated with a decrease in DNMT3a expression in the hippocampus of fetal male rats (Glendining et al., 2018). Similarly, we found that transcript abundance of DNMT1 in the amygdala was sensitive to changes in maternal diet during early life. However, in adulthood transcript abundance of DNMTs and other epigenetic regulators remained unchanged between diet groups (Fig. 2B). Notably, the increase in DNMT1 and MeCP2 expression in early life in the amygdala was associated with a robust increase in DNMT enzymatic activity (Fig. 3A) and global LINE-1 DNA methylation (Fig. 3B). It has been reported that DNMT1 and MeCP2 form a complex on hemimethylated DNA to regulate DNMT activity (Kimura and Shiota, 2003). The strong association between reduced miR-148/152 levels with increased expression of DNMT1 and MeCP2, combined with increased DNMT enzymatic activity and global DNA methylation suggest a mechanism by which mHFD exposure may program the DNA methylome of offspring during early life, and an important regulatory role for lactation-specific miR-148/152 in this process.

We also examined whether changes in DNA methylation patterns we observed during early life were maintained into adulthood, long after the period of mHFD exposure. In adulthood, DNMT enzymatic activity showed a decreasing trend (Fig. 4A), along with a significant reduction in global DNA methylation levels in offspring exposed to mHFD (Fig. 4B). These findings support previous reports showing persistent global DNA hypomethylation in adult offspring exposed to mHFD (Carlin et al., 2013; Vucetic et al., 2010).

Next, we examined genome-wide DNA methylation patterns at single nucleotide resolution using RRBS. Over 60% of mHFD-associated DMRs showed hypomethylation and approximately 50% were found in intergenic regions (Fig. 5A). Notably, the proportion of DMRs found within promoter regions was substantially higher in early life (21%) compared to adulthood (6%). It is possible that this higher proportion of differential DNA methylation in promoters may be linked to the rapid neurodevelopmental processes that are taking place during the first week of postnatal life. Earlier studies have reported unique temporal patterns of DNA methylation modifications in the brain during perinatal life, where a reversal in the direction of genomewide methylation occurs from prenatal to postnatal development (Lister et al., 2013; Numata et al., 2012). In particular, global methylation levels were shown to decrease during prenatal life and increase postnatally (Numata et al., 2012). However, it should be noted that P7 animals in this study were still under the direct influence of the mHFD and its associated metabolic milieu, conditions that were not present in adulthood, as the animals were weaned onto a control diet. Interestingly, we found that exposure to mHFD during perinatal life also lead to increased global LINE-1 methylation in the amygdala (see Fig. 3B). Taken together, our data suggest an increased period of sensitivity of the neural DNA methylome to dietary stress in early life.

We found a total of 57 shared DMRs in both early life and adulthood (Fig. 5B), corresponding to 26 genes annotated to these regions (Table 1). Interestingly, the promoter region of MeCP2 was hypomethylated in response to mHFD across both ages (Table 1); we also observed an increase in the relative transcript abundance of MeCP2 in early life (see Fig. 2A). Indeed, studies have shown that methylation of CpG sites of six previously characterized *cis* regulatory sequences, which are found in MeCP2 promoter and intron 1, is inversely correlated with MeCP2 transcript abundance level (Liyanage et al., 2019; Olson et al., 2014). In contrast to early life, MeCP2 transcript abundance in adulthood remained unchanged between offspring exposed to mHFD and mCHD (Fig. 2B), despite the apparent maintenance of promoter hypomethylation among mHFD offspring. These findings suggest an alternate mode of transcriptional control of MeCP2. Post-transcriptional regulation of MeCP2 by polyadenylation and a number of micro-RNAs during development has been reported (McGowan and Pang, 2015; Samaco et al., 2004). Polyadenylation of MeCP2 transcript results in a longer 3’UTR that has been shown to be highly expressed in the brain at birth, decrease progressively during postnatal development, and increase again in adulthood, regulated by RNA-binding proteins and mirco-RNAs (McGowan and Pang, 2015; Rodrigues et al., 2016; Samaco et al., 2004).

MeCP2 is associated with the regulation of differentially expressed genes in offspring exposed to mHFD. For example, mHFD exposed animals show reduced μ-opioid receptor transcript abundance, promoter hypermethylation, and increased recruitment of MeCP2 to the μ-opioid receptor promoter in reward-related brain regions, including the ventral tegmental area, prefrontal cortex, and nucleus accumbens (Vucetic et al., 2011). In addition, MeCP2 is known to regulate the expression of brain-derived-neurotrophic factor (BDNF) in a dynamic mechanism that could either repress or activate its expression (reviewed in Li and Pozzo-Miller, 2014). Studies have shown a reduction in BDNF transcript levels in several brain regions including the prefrontal cortex, hippocampus, and hypothalamus in developing animals exposed to mHFD (Bae-Gartz et al., 2019; Rincel et al., 2016; Tozuka et al., 2009). It is possible that DNA methylation modifications and changes in transcript abundance in MeCP2 in response to mHFD are involved in the differential gene expression reported in the brains of mHFD offspring.

We identified overlapping GO terms (Fig. 6; Supplementary Table 4) and KEGG pathways (Table 2) that were enriched with differentially methylated genes in early life and adulthood to contextualize differentially methylated genes associated with mHFD exposure. GO terms associated with the development of the nervous system (i.e. neurogenesis, neuron development, and nervous system development) and neuronal projections (i.e. neuron projection morphogenesis, axonogenesis, plasma membrane bounded cell projection organization), as well as axon guidance KEGG pathway were highly enriched with genes annotated to DMRs in early life and adulthood. Several earlier studies have reported altered neuronal morphology in brain limbic regions with exposure to mHFD. Specifically, new hippocampal neurons exhibit impaired dendritic arborization in young mice (P35) exposed to mHFD, which was associated with reduced BDNF and impaired spatial learning (Tozuka et al., 2009). Another study showed that mHFD is associated with reduced dendritic length and complexity in the hippocampus and amygdala of adult rat offspring (Janthakhin et al., 2017). Young and adult mouse offspring exposed to mHFD exhibit reduced stability and loss of dendritic spines (Hatanaka et al., 2015). Taken together, our finding may implicate DNA methylation modifications in these effects, though this remains to be examined more in detail.

The enrichment analysis also identified age-specific GO terms and KEGG pathways (Fig. 7–8; Supplementary Table 7-8). In early life, GO terms and KEGG pathways involving neuronal development, such as axon guidance, Wnt signaling, neurotrophin signaling and thyroid hormone signaling pathways were enriched with DMRs. Interestingly, previous studies have linked changes in thyroid hormone levels and genes involved in thyroid synthesis to mHFD exposure in offspring across several species, including humans, rodents, (Kahr et al., 2016; Tabachnik et al., 2017) as well as primates (Suter et al., 2012). In particular, thyroid hormone levels and the genes that mediate thyroid hormone synthesis decreased in the hypothalamus and thyroid gland in human fetuses that were exposed to an obesogenic maternal environment (Suter et al., 2012). Our findings implicating DNA methylation modifications in thyroid hormone signaling pathways thus warrant further investigation. In adulthood, glutamatergic and GABAergic KEGG pathways were enriched with genes annotated to DMRs (Supplementary Table 8). Anxiety-like behaviour is a well-characterized outcome of dysregulation in glutamatergic and GABAergic systems, and these findings concur with previously published studies showing heightened anxiety-like behaviours in mHFD offspring (Bilbo and Tsang, 2010; Peleg-Raibstein et al., 2012; Sasaki et al., 2013).

## Conclusion

Maternal milk is a primary source of nutrition in early life in mammals, and plays an important role in growth, neurodevelopment, immunity, microbiome composition, and behaviour (Andreas et al., 2015; Bagnell and Bartol, 2019; Ballard and Morrow, 2013; Boquien, 2018; De Leoz et al., 2015; Der et al., 2006; Dettmer et al., 2018; Gareau, 2011; Hamosh, 2001). Collectively, our findings suggest a role for lactation-specific miRNAs in neurodevelopmental programming of the DNA methylome by mHFD. With the significant rise worldwide in the number of women of reproductive age who are overweight or obese, understanding the role of miRNA transfer via maternal milk during early development may help identify mechanisms that lead to adverse health outcomes in children, and ultimately enable preventive and therapeutic interventions to support offspring health.

## Materials and Methods

### Animal Treatment and Handling

Adult female Long Evans rats (7 week) were purchased from Charles River, Canada (St. Constant, QC) and housed with same sex pairs until mating and maintained on a 12:12 - h light– dark cycle (lights on 7:00 am–7:00 pm) with *ad libitum* access to food and water. Females were maintained on either house chow diet (mCHD; 5001; Purina Lab Diets, St. Louis, MO, USA) consisting of 28.5 % protein, 13.5 % fat, and 58 % carbohydrate, or a high fat diet (mHFD; D12492; Research Diets Inc. New Brunswick, NJ, CA), consisting of (by kcal): 20 % protein, 60 % fat, 20 % carbohydrate) 4 weeks prior to mating, during gestation, and lactation (Abuaish et al., 2018; Sasaki et al., 2014, 2013). Mating was conducted over a one-week period and sperm plugs were checked twice a day to determine the onset of pregnancy. Females were then separated from males and singly housed throughout pregnancy. After parturition, dams were moved into clean cages and offspring were weighed and culled to 12 pups / litter (6 males and 6 females) when possible.

At sacrifice, female P7 neonates were individually removed from their litters, rapidly decapitated; brains and curd stomach milk were collected. Female P90 adults were sacrificed by CO_2_ inhalation followed by decapitation to collect brains. Separate litters of animals were used for the various assays: n=4 animals (1/litter/diet group) for LINE-1, DNMT activity, and qPCR analyses at P7; n=4 animals (1/litter/diet group) for RRBS and qPCR analyses at P7; n=4 animals (1/litter/diet group) for milk miRNA analysis at P7; n=4-6 (1/litter/diet group) for LINE-1, DNMT activity and qPCR analyses at P90; and n=4 (1/litter/diet group) for RRBS analysis at P90. Brains obtained from all animals were flash frozen in isopentane and dry ice and stored at −80 °C for later usage. Brains were cryosectioned into 50 μM sections using Research Cryostat Leica CM3050 S (CM3050 S; Leica Biosystems, Concord, ON, CAD) and the amygdala was microdissected using stereotaxic coordinates (P7: bregma: −0.20mm to −1.60 mm (Paxinos et al., 1991); P90 bregma: −1.72mmto −3.00mm (Paxinos and Watson, 2007).

We focused on female offspring in this study for several reasons. First, earlier investigations found greater transcriptional differences in the brains of female offspring compared to male offspring exposed to mHFD in early life (Abuaish et al., 2018; Barrand et al., 2017; Sasaki et al., 2014) and in adulthood (Sasaki et al., 2013). Second, adult female offspring showed stronger behavioral and physiological alterations in response to mHFD, in comparison to male littermates (Sasaki et al., 2013). These alterations were associated with pronounced changes in gene expression in the amygdala (Sasaki et al., 2013).

All experimental protocols were approved by the Local Animal Care Committee at the University of Toronto, Scarborough, and were in accordance with the guidelines of the Canadian Council on Animal Care.

### RNA Extraction from Stomach Milk and Amygdala

Total RNA, >18 nucleotides, was purified from curd stomach milk collected from P7 offspring that were exposed to mCHD (n=4) or mHFD (n=4) using a combination of QIAzol and miRNeasy Mini Kit (217004; Qiagen, Toronto, ON, CA) as described previously (Izumi et al., 2014, 2013). RNA was extracted from the amygdala of female offspring at P7 and P90 (n=6/mCHD, n=6/mHFD) using TRIzol Reagent (15596018; ThermoFisher Scientific, Ottawa, ON, CAD) according to manufacturer’s instructions. RNA concentration and quality were measured using a Nanodrop Spectrophotometer (ND-2000C; ThermoFisher Scientific) and RapidOut DNA Removal Kit (K2981; ThermoFisher Scientific) was used to remove sources of genomic DNA contamination. The samples were stored at −80 °C for future use.

### miRNA Primer Design

miRNA-specific forward primers for miR-148-5P, 148-3P, 152-5P, 152-3P, and miR-21-5P were designed using annotated mature miRNA sequences obtained from miRBase (Kozomara and Griffiths-Jones, 2011) for *Rattus norvegicus,* using a protocol previously described (Biggar et al., 2014). All miRNA targets were quantified in conjunction with a universal reverse primer (Supplementary Table 1) and a miRNA target-specific forward primer (Supplementary Table 1). Four reference genes (U6 snRNA, 5S rRNA, Snord 96a, and Snord 95a) were tested to determine a set of most stable internal controls to be used for data normalization (Supplementary Table 1). Sequences for the internal reference genes, were based on previously published work on rodent-specific miRNA regulation in the brain (Eacker et al., 2011; Minami et al., 2014).

### miRNA Expression Analysis by RT-qPCR

125 ng of total RNA extracted from curd stomach milk and 1 μg of total RNA extracted from the amygdala of the same animals at P7 were processed for miRNA analysis as previously described (Biggar et al., 2014, 2011). miRNA levels in stomach milk and amygdala of P7 offspring were quantified using a StepOne Plus real-time thermocycler with Fast SYBR Green PCR master mix (4385612; Applied Biosystems, Foster City, CA, USA) using a previously established protocol (Biggar et al., 2014). A melt curve analysis was done following each RT-qPCR reaction to ensure that miRNA primers did not yield multiple PCR products.

Five lactation-specific miRNAs were quantified in ingested stomach milk (miR-148-5P, miR-148-3P, miR-152-5P, miR-152-3P, and miR-21-5P) and four of the same miRNAs were quantified in the amygdala at P7 (miR-148-5P, miR-148-3P, miR-152-5P, and miR-152-3P; Supplementary Table 1). miR-21-5P was below the detectable range by RT-qPCR in the amygdala at P7. miRNA quantification was done using ΔΔ**C**_q_ method and all targets were normalized against the GEOmean of two internal reference genes. U6 snRNA and 5S rRNA were determined to be suitable internal controls for both stomach milk and amygdala datasets by NormFinder software (Andersen et al., 2004). Relative miRNA levels were denoted as mean ± SEM representing n=4 (stomach milk) and n=6 (amygdala) biological replicates per experimental condition and three technical replicates per biological replicate. Significant differences across mCHD and mHFD exposures were measured using an independent sample Student’s t-test with a 95 % confidence interval (p<0.05).

### mRNA Expression Analysis by RT-qPCR

Gene expression levels of DNMT1, DNMT3a, DNMT3b, MeCP2, and GADD45 were measured using a StepOne Plus real-time thermocycler with a Fast SYBR Green PCR master mix (4385612; Applied Biosystems) in the amygdala during early life and adulthood. A melt curve analysis was done following each RT-qPCR reaction to ensure that primers did not yield multiple PCR products. Primers (Supplementary Table 1) were designed using nucleotide sequence information available at the National Center for Biotechnology Information (NCBI): www.ncbi.nlm.nih.gov and previously published research (Sasaki et al., 2014, 2013).

mRNA quantification was determined using a standard curve consisting of 11 serial dilutions ranging from 500 to 0.49 ng/μL. Quantity means of each target was normalized against the GEOmean of four reference genes, YWAZ, GAPDH, 18s, and Actin B. These four reference genes were determined to be suitable internal controls in the female amygdala during early life and adulthood by NormFinder Software (Andersen et al., 2004). Relative gene expression levels were denoted as mean ± SEM representing n=6 biological replicates per experimental condition and three technical replicates per biological replicate. Significant differences across mCHD and mHFD exposures were measured using an independent sample Student’s t-test with a 95 % confidence interval (p<0.05).

### Protein Extraction

Total soluble protein was extracted using the organic phase of the TRIzol extraction (n=6/mCHD, n=6/mHFD) from female amygdala. Ethanol (100%; 1:0.3 v/v to TRIzol) was added to the organic phase and centrifuged at 2,000 x g for 5 min at 4 °C to pellet the DNA. Isopropanol (1:1.5 v/v to TRIzol) was added to the supernatant, incubated at RT for 10 min, and later centrifuged at 12,000 x g at 4 °C for 10 min to pellet the proteins. Protein pellet was washed 3x with 0.3 M guanidine hydrochloride in 95 % ethanol (1:2 v/v to TRIzol) followed by a single wash with 2 mL of 100 % ethanol. The pellet was air dried for 15 min and re-suspended in 200 μL of 1 % SDS. The protein concentrations were measured using Pierce ™ BCA Protein Assay Kit (23225; Thermo Scientific) with a Albumin BSA Standards ranging from 2000 μg/mL to 0 μg/mL using a Nanodrop Spectrophotometer cuvette system (ND-2000C; ThermoFisher Scientific).

### DNA Methyltransferase Activity

Total DNMT activity was measured using EpiQuik™ DNA Methyltransferase Activity/Inhibition Assay Kit (P-3001; Epigentek, Farmingdale, NY, USA) according to manufacturer’s instructions. Briefly, a dilution curve ranging from 5 μg to 60 μg was assayed using pooled protein samples from P7 and P90 amygdala to determine the linear portion of the absorption curve along with DNMT positive controls (50 μg/mL; provided by Epigentek) and blanks (containing only assay buffer). According to the dilution curve, 50 μg of total protein was chosen as the optimal range for both P7 and P90 amygdala samples. All samples, positive controls, and blanks were run in duplicates according to manufacturer’s instructions. Absorption was read using a microplate reader (Versamax; Molecular Devices) at 450 nm within 2 min. Total DNMT enzymatic activity was calculated using the following formula:

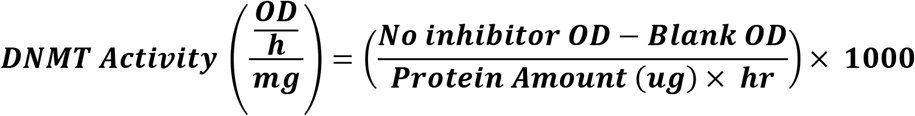

### Genomic DNA Extraction

Genomic DNA (gDNA) was extracted using ZR-Duet™DNA/RNA MiniPrep (D7005; Zymo Research, Irvine, CA, USA) according to manufacturing instructions (n=4/mCHD, n=4/mHFD). The concentration of gDNA was quantified using the Pico Green dsDNA Assay Kit (P11496; ThermoFisher Scientific).

### Global DNA Methylation

Methylation of LINE-1 repeats have been implicated in several complex diseases and used as representative of global levels of DNA methylation, as LINE-1 repeats make up approximately 18 % of the genome with copy number estimated at roughly half a million (Weisenberger et al., 2005). Furthermore, LINE-1 methylation has a strong association with adiposity, dietary weight gain, and body fat mass in humans (Carraro et al., 2016). As such, we used LINE-1 methylation to measure changes in global methylation and the effects of mHFD exposure on 5mC levels. Global % 5mC levels were assayed using a Global DNA Methylation LINE-1 Kit (55017; Active Motif, Carlsbad, CA, USA) according to the manufacturer’s instructions. A standard curve (ranging from 0 ng to 100 ng) was prepared in triplicates using a mixture of methylated and non-methylated DNA standards. A dilution curve (ranging from 0 ng to 500 ng) consisting of pooled P7 and P90 gDNA samples were run in duplicate alongside the standard curve to determine the linear portion of the absorbance readings at 450 nm. Based on the standard curve, 250 ng of gDNA from P7 and 1 μg of gDNA from P90 were chosen for the assay. All absorption readings were taken using a microplate reader (Versamax; Molecular Devices, San Jose, CA, USA) at 450 nm with a reference wavelength of 655 nm within 5 min.

% 5mC levels were calculated by averaging the duplicates for the blanks and sample wells and triplicates for the standards and subtracting the average blank OD at 450 nm from the average standard ODs and sample ODs. % 5mC levels that were associated with each sample were determined based on the total detectable CpG content. Using the standard curve ranging from 0 ng to 100 ng (slope = 0.0297; R_2_-value = 0.950), % 5mC methylation for each sample was extrapolated using MyCurveFit Beta online curve fitting software (https://mycurvefit.com). As the amount of DNA used was 60 ng and 80 ng for P7 and P90, respectively, the extrapolated % 5mC values were further divided by the total amount of gDNA loaded and multiplied by 100.

### Reduced Representation Bisulfite Sequencing (RRBS)

gDNA from P7 and P90 female amygdala used for the global methylation assay was also used for RRBS analysis. Briefly, 100 ng of genomic DNA was used to construct libraries for RRBS using the Ovation RRBS Methyl-Sequencing Kit (0553-32; NuGEN, Redwood City, CA, USA) as per the manufacturer’s instructions. The EpiTect Fast DNA Bisulfite Kit (59824; Qiagen) was used for bisulfite conversion. Libraries were sequenced on a NextSeq 500 (Illumina, San Diego, CA, USA) at the Princess Margaret Genomics Centre (University Health Network, Toronto) using single end sequencing with a 75-base pair read length and multiplexed at 8–10 samples per flowcell. Samples sequenced were as follows; for P7, mCHD: n=4 and mHFD: n=4 and for P90, mCHD: n=6 and mHFD: n=4.

### DNA Methylation Analysis

RRBS fastq files were processed as per our previous study. Files were trimmed to remove low quality reads (q < 30) and adaptors, and later aligned to RGD Rnor_6.0 (Ashbrook et al., 2018). The average reads per sample was 20.8×10_6_ with a mapping efficiency of 50-60 %. Differentially methylated regions (DMRs) were identified using a dynamic sliding window approach and annotated using the methylPipe (Song et al., 2013) and compEpiTools R packages (Kishore et al., 2015). The bisulfite conversion rate was >99 % for all samples. Regions were declared to be differentially methylated if the average methylation of at least 10 consecutive CpG sites within a 1 Kb region was ≥ 5 % between mHFD and mCHD with FDR < 0.05.

### Gene Set Enrichment Analysis

Set of differentially methylated genes identified by the DMR analysis were explored using Gene Ontology Biological Processes (GO BP) (Ashburner et al., 2000; Blake et al., 2015) and KEGG pathway enrichment analysis (Kanehisa et al., 2012) to examine functionally annotated gene pathways and networks. Enrichment analysis was performed using gProfiler (Reimand et al., 2016) with a FDR ≤ 0.05 cutoff for statistical significance.

Networks of GO terms were constructed and visualized using Enrichment Map on Cytoscape 3.6.1 (Merico et al., 2010). Clusters were arranged and labeled using the yFiles algorithm and WordCloud plugin on Cytoscape 3.6.1. GO analysis was also performed using DAVID algorithm to compare to the results generated by gProfiler.

### Statistical analysis

Statistical analysis was carried out using SPSS version 25 (IBM) and figures were constricted using GraphPad Prism7. A Shapiro-Wilk test was used to test for normality for all datasets. All data exhibited a normal distribution and thus parametric analyses were carried out. Maternal and offspring bodyweights were analyzed using repeated measures analysis of variance (ANOVA) Diet X Time. Dam’s caloric intake, offspring relative transcript abundance of miRNA and mRNA targets, DNMT enzymatic activity (OD/h/mg), and the global LINE-1 DNA methylation (%), between mCHD and mHFD were analyzed using a two-tailed student t-test with a 95 % confidence interval. Pearson correlation analysis was used to assess the relationship between DNMT activity and % global DNA methylation levels. All relationships were considered statistically significant at p ≤ 0.05.

## Acknowledgements

This work was supported by a discovery grant from the Natural Sciences and Engineering Research Council (NSERC) of Canada to Dr. Patrick O. McGowan. Dr. Sameera Abuaish was supported by a Graduate fellowship from Princess Nourah bint Abdulrahman University. Dr. Sanoji Wijenayake holds a NSERC postdoctoral research fellowship.

## Author Contribution

S.A, S.W, and P.O.M contributed to the experimental design, conceptualization, and wrote the paper.

S.A contributed to animal testing and sample collections for the P7 cohort, generated RRBS libraries, and performed enrichment and network analysis.

S.W contributed to miRNA measurements and analysis, DNMT enzymatic assays and LINE-1 methylation assays.

W.C.D contributed to RRBS library construction of P90 samples, performed all bioinformatics analysis of the sequencing data and generated the DMRs datasets.

S.A, S.W, W.C.D contributed to gDNA, RNA, and protein extractions from P7 and P90 samples.

S.W and C.W.M.L contributed to P7 and P90 qPCR measurements.

A.S contributed to P90 animal testing and sample collections.

## Data availability

The data sets generated and/or analyzed during the current study are available from the corresponding author on request.

