## Supplementary Figures for "Maternal high fat diet alters lactation-specific miRNA expression and programs the DNA methylome in the amygdala of female offspring"

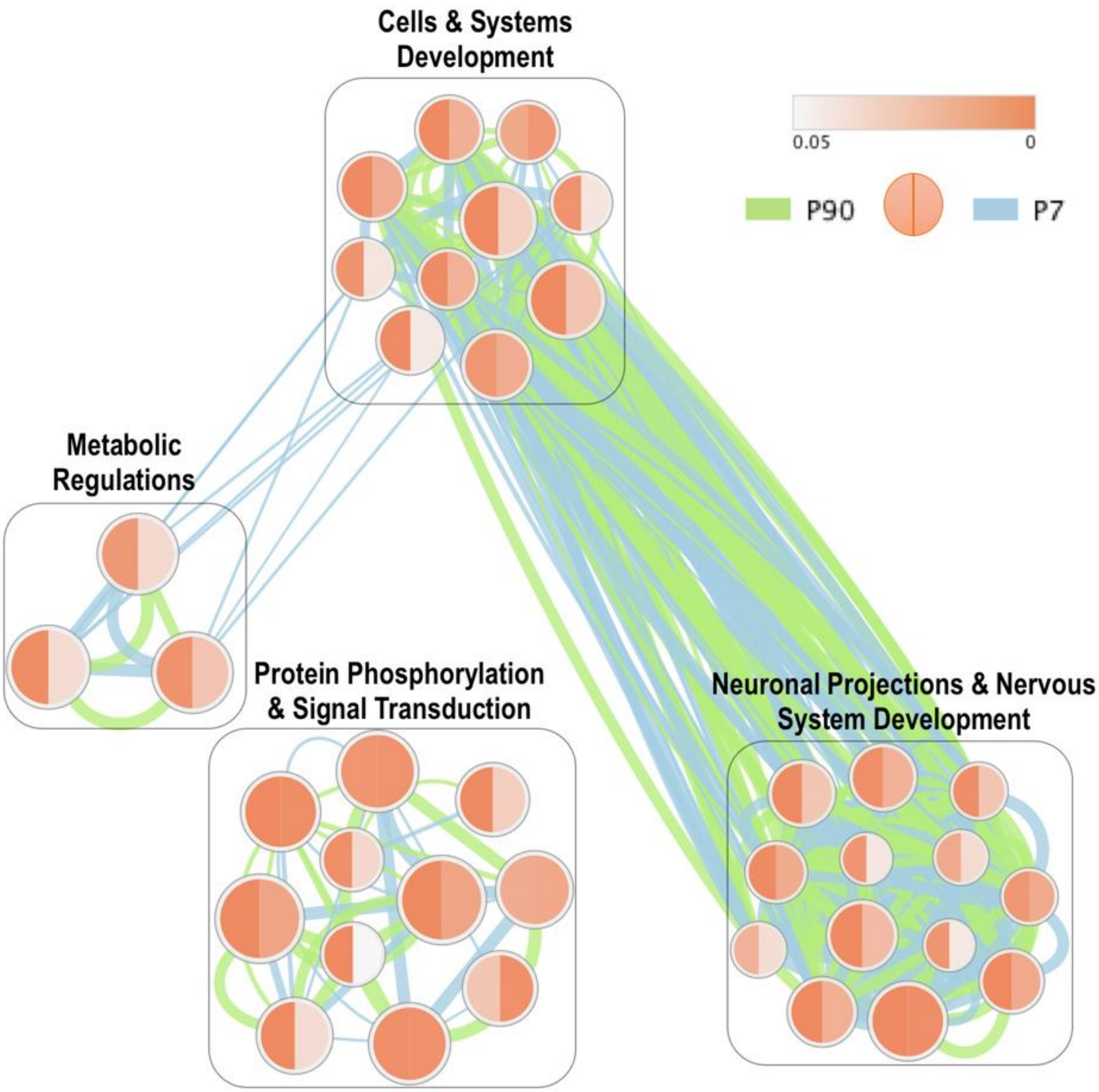


**Supplementary Figure 1.** Overlapping clusters DAVID Gene Ontology Biological Processes enriched in P7 (right half of nodes and blue edges) and P90 (left half of nodes and green edge) differentially methylated gene set for female offspring exposed to mHFD. The size of the nodes represents the number of genes, while the color indicates the FDR p-value. The edges between the nodes indicate shared genes, with edge thickness representing the number of genes in common.

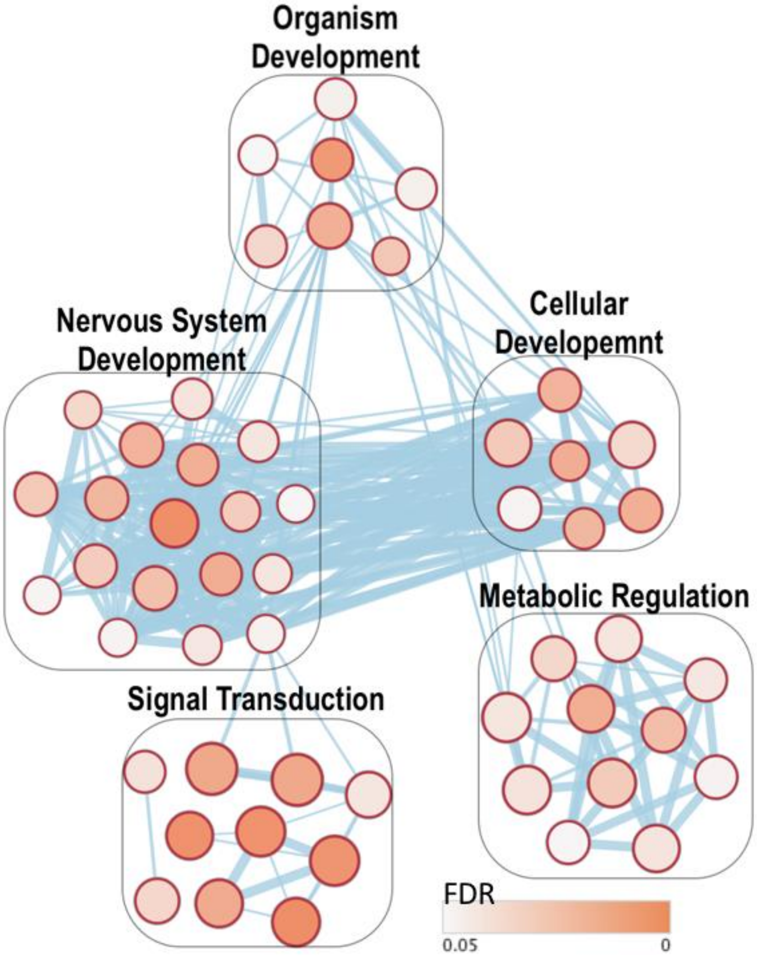


**Supplementary Figure 2.** DAVID clusters of Gene Ontology Biological Processes enriched at P7 in female offspring in response to mHFD exposure in the amygdala. The size of the nodes represents the number of genes, while the color indicates the FDR p-value. The edges between the nodes indicate shared genes, with edge thickness representing the number of genes in common.

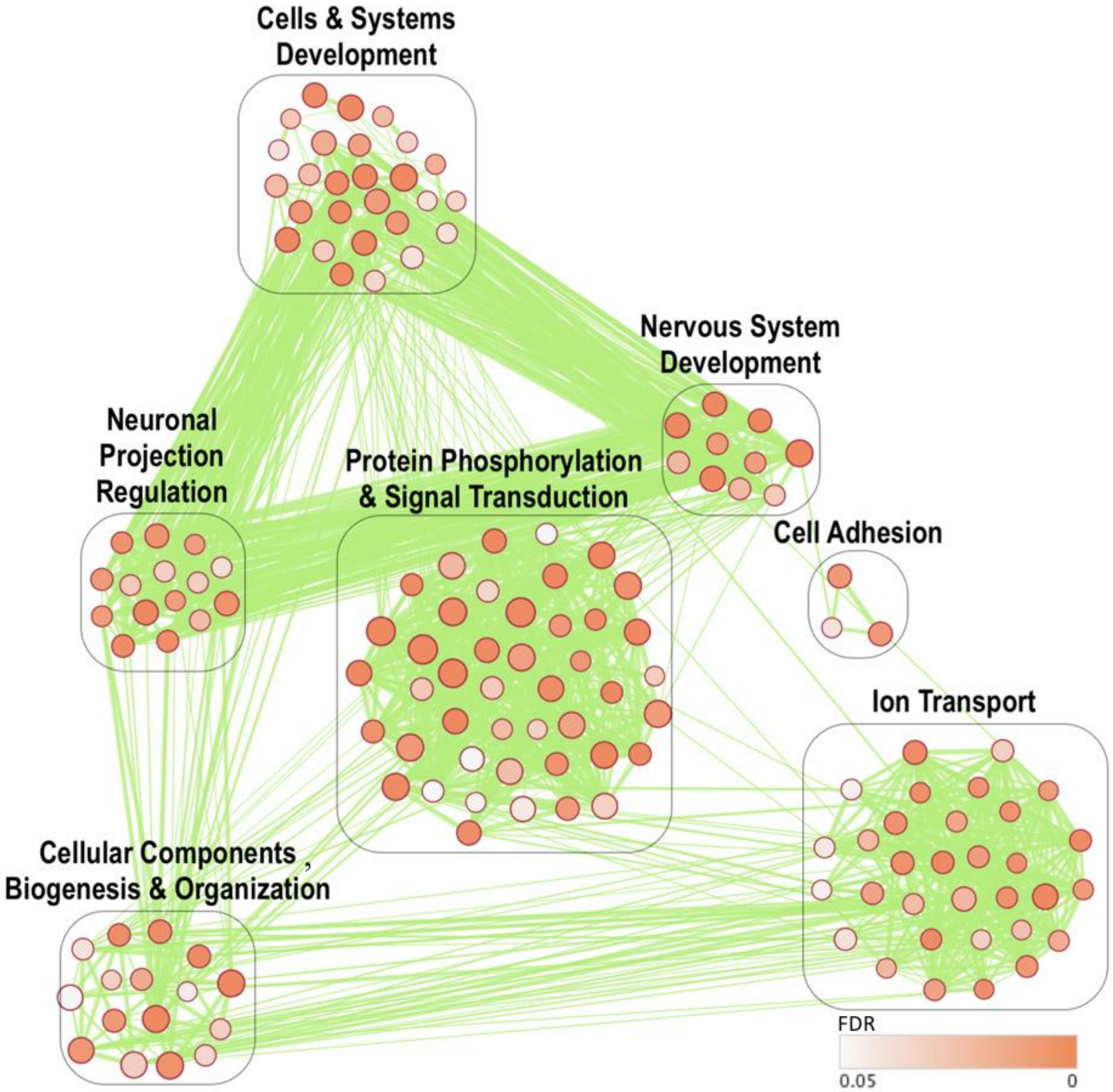


**Supplementary Figure 3.** DAVID clusters of Gene Ontology Biological Processes enriched at P90 in female offspring in response to mHFD exposure in the amygdala. The size of the nodes represents the number of genes, while the color indicates the FDR p-value. The edges between the nodes indicate shared genes, with edge thickness representing the number of genes in common.
