## Supplementary Tables for "Maternal high fat diet alters lactation-specific miRNA expression and programs the DNA methylome in the amygdala of female offspring"

**Supplementary Table 1.** miRNA and mRNA primers.

| Transcript | Forward Primer (5' – 3') | Reverse Primer (5' – 3') |
|---|---|---|
| rno-U6 snRNA | ACA CTC CAG CTG GGG TGC TCG CTT CGG CAG C | - |
| rno-5S rRNA | GGC CTG GTT AGT ACT TGG ATG G | - |
| rno-miR-148-5P | ACA CTC CAG CTG GGA AAG TTC TGA GAC ACT C | - |
| rno-miR-148-3P | ACA CTC CAG CTG GGT CAG TGC ACT ACA GAA C | - |
| rno-miR-152-5P | ACA CTC CAG CTG GGA GGT TCT GTG ATA CAC T | - |
| rno-miR-152-3P | ACA CTC CAG CTG GGT CAG TGC ATG ACA GAA C | - |
| rno-miR-21-5P | ACA CTC CAG CTG GGT AGC TTA TCA GAC TGA T | - |
| Universal Primer | CTC ACA GTA CGT TGG TAT CCT TGT G | - |
| Stem Loop Adaptor Primer | GAT GTT CGA TGC CAT ATT GTA CTG TGA GTT TTT TTT TVN | - |
| YWAZ | TTG AGC AGA AGA CGG AAG GT | GAA GCA TTG GGG ATC AAG AA |
| GAPDH | ACA TCA AAT GGG GTG ATG CT | GTG GTT CAC ACC CAT CAC AA |
| Actin β | TTT GAG ACC TTC AAC ACC CC | ATA GCT CTT CTC CAG GGA GG |
| 18S rRNA | ATG GTA GTC GCC GTG CCT A | CTG CTG CCT TCC TTG GAT G |
| DNMT1 | ACC TAC CAC GCC GAC AT | AGG TCC TCT CCG TAC TCC A |
| DNMT3a | ACG CCA AAG AAG TGT CTG CT | CTT TGC CCT GCT TTA TGG AG |
| DNMT3b | GAT GTG ACA CCT AAG AGC AGC AGT AC | CAA ACT CCT TGT CAT CCT GAT ACT CA |
| MeCP2 | CAA ACA GCG ACG TTC CAT CA | TGT TTA AGC TTT CGC GTC CAA |
| GADD45α | GCT GGC CAT AGA CGA AGA AG | GCC TGA TAC CCT GAC GAT GT |

rno: *Rattus norvegicus*; U6 snRNA: U6 spliceosomal RNA; 5S rRNA: 5S ribosomal RNA; YWAZ: 14-3-3 protein zeta/delta; GAPDH: Glyceraldehyde-3-phosphate dehy

For Sameera: If all the other supplementary tables are not being included in the manuscript, this should also not be included.

Please remove this from the manuscript and upload it as a separate supplementary file during submission.

**Supplementary Table 2: List of annotated genes to**

| Chromosom | Gene Symbol | ation differen | Location |
|---|---|---|---|
| **chr1** | Tbcb | 5.555 | Promoter |
| **chr1** | Clasrp | -23.102 | Genebody |
| **chr1** | Cyp2c7 | -30.46 | Genebody |
| **chr1** | Dhx34 | -41.313 | Genebody |
| **chr1** | Eef2k | 15.238 | Genebody |
| **chr1** | Eml2 | 11.677 | Genebody |
| **chr1** | Gprc5b | 12.41 | Genebody |
| **chr1** | Hbs1l | 13.08 | Genebody |
| **chr1** | Igf2r | -5.923 | Promoter |
| **chr1** | Il20ra | -20.132 | Genebody |
| **chr1** | LOC310926 | -20.203 | Genebody |
| **chr1** | LOC310926 | -21.01 | Genebody |
| **chr1** | Map3k4 | -23.94 | Genebody |
| **chr1** | Mrvi1 | -15.363 | Genebody |
| **chr1** | Nav2 | -10.06 | Genebody |
| **chr1** | Ovol3 | 5.555 | Genebody |
| **chr1** | Pepd | 5.555 | Genebody |
| **chr1** | Plagl1 | 8.992 | Promoter |
| **chr1** | Prx | 5.113 | Genebody |
| **chr1** | Rbm20 | -28.063 | Genebody |
| **chr1** | Relt | 8.665 | Genebody |
| **chr1** | Rn5-8s | 10.092 | Promoter |
| **chr1** | Shank2 | 8.457 | Genebody |
| **chr1** | Slit1 | -6.785 | Genebody |
| **chr1** | Smarca2 | -16.865 | Genebody |
| **chr1** | Smok2a | -25.675 | Genebody |
| **chr1** | Ssc5d | -8.803 | Genebody |
| **chr1** | Synj2 | -5.985 | Genebody |
| **chr1** | Tbc1d10b | -11.427 | Promoter |
| **chr1** | Tenm4 | -6.778 | Genebody |
| **chr10** | Anxa6 | 20.15 | Genebody |
| **chr10** | Arl4d | 17.277 | Promoter |
| **chr10** | Armc7 | -13.075 | Genebody |
| **chr10** | Ccdc40 | -20.665 | Genebody |
| **chr10** | Ccdc40 | -14.155 | Genebody |
| **chr10** | Csnk1d | -41.875 | Genebody |
| **chr10** | Galnt10 | -18.68 | Promoter |
| **chr10** | Jmjd8 | -33.952 | Promoter |
| **chr10** | Khps1a | -5.532 | Promoter |

| | | | |
|---|---|---|---|
| **chr10** | Krt28 | 15.473 | Promoter |
| **chr10** | Map3k14 | -7.5 | Genebody |
| **chr10** | Meox1 | 10.963 | Genebody |
| **chr10** | Nags | 5.933 | Genebody |
| **chr10** | Ntn1 | -13.197 | Genebody |
| **chr10** | Plcd3 | -12.082 | Genebody |
| **chr10** | Rhbdf1 | -27.255 | Promoter |
| **chr10** | Scn4a | -30.967 | Genebody |
| **chr10** | 07-Sep | -43.77 | Genebody |
| **chr10** | Sphk1 | -5.532 | Genebody |
| **chr10** | Srcin1 | 24.65 | Genebody |
| **chr10** | Wdr24 | -33.952 | Genebody |
| **chr10** | Zmynd15 | 12.135 | Genebody |
| **chr11** | App | -7.85 | Genebody |
| **chr11** | Cmss1 | -10.25 | Promoter |
| **chr11** | Golgb1 | -15.58 | Genebody |
| **chr11** | Gtf2e1 | -10.285 | Genebody |
| **chr11** | Hic2 | -7.472 | Genebody |
| **chr11** | Runx1 | -14.58 | Promoter |
| **chr11** | Tiam1 | 22.95 | Genebody |
| **chr12** | Arhgef18 | -23.502 | Genebody |
| **chr12** | Cit | -8.355 | Genebody |
| **chr12** | Cit | -6.173 | Genebody |
| **chr12** | Col26a1 | -20.517 | Genebody |
| **chr12** | Flt1 | 19.235 | Genebody |
| **chr12** | Hip1r | -37.152 | Genebody |
| **chr12** | Ncor2 | -11.325 | Genebody |
| **chr12** | Ncor2 | 10.977 | Genebody |
| **chr12** | Pitpnm2 | -5.77 | Genebody |
| **chr12** | Rimbp2 | -7.048 | Genebody |
| **chr12** | Sez6l | -15.805 | Genebody |
| **chr12** | Tpst2 | -8.54 | Genebody |
| **chr12** | Vps37b | 7.088 | Genebody |
| **chr13** | Cdh20 | -6.75 | Genebody |
| **chr13** | Lrrn2 | -7.603 | Genebody |
| **chr13** | Mark1 | -13.607 | Genebody |
| **chr13** | Rabgap1l | -51.85 | Genebody |
| **chr13** | Rgs8 | 7.38 | Genebody |
| **chr13** | Smyd3 | -6.27 | Genebody |
| **chr14** | Bcl11a | -9.852 | Genebody |
| **chr14** | Bcl11a | -8.957 | Genebody |
| **chr14** | Camk2b | -16.185 | Genebody |

| | | | |
|---|---|---|---|
| **chr14** | Commd1 | 6.315 | Genebody |
| **chr14** | Grb10 | -5.973 | Promoter |
| **chr14** | LOC680039 | -40 | Genebody |
| **chr14** | Mapk10 | -14.935 | Genebody |
| **chr14** | Rab5a | -7.45 | Promoter |
| **chr14** | Rn18s | -25.253 | Promoter |
| **chr14** | Rn45s | -25.253 | Promoter |
| **chr14** | Tns3 | -17.968 | Genebody |
| **chr14** | Zrsr1 | 6.315 | Promoter |
| **chr15** | Abcc4 | 6.332 | Genebody |
| **chr15** | Bmp1 | -30.54 | Genebody |
| **chr15** | Bmp4 | 35.46 | Genebody |
| **chr15** | Carmil3 | -25.709 | Genebody |
| **chr15** | Dmtn | -21.235 | Genebody |
| **chr15** | Dupd1 | 11.21 | Genebody |
| **chr15** | Fbxo34 | -5.152 | Genebody |
| **chr15** | Lrch1 | -38.372 | Genebody |
| **chr15** | RGD1565725 | 7.252 | Genebody |
| **chr15** | Stk24 | 7.808 | Genebody |
| **chr16** | Ank1 | 5.695 | Genebody |
| **chr16** | Gas6 | 7.405 | Promoter |
| **chr16** | Grid1 | 9.09 | Genebody |
| **chr16** | Grid1 | 11.09 | Genebody |
| **chr16** | Mast3 | -7.375 | Genebody |
| **chr16** | Plvap | 7.403 | Promoter |
| **chr16** | Sh2d4b | 10.875 | Genebody |
| **chr16** | Slc20a2 | -32.708 | Genebody |
| **chr17** | Adarb2 | -5.095 | Genebody |
| **chr17** | Atxn1 | 6.075 | Genebody |
| **chr17** | Dip2c | 7.26 | Genebody |
| **chr17** | Gpr50 | -17.002 | Promoter |
| **chr17** | Gpr50 | -5.813 | Promoter |
| **chr17** | Rnf144b | -15.645 | Genebody |
| **chr17** | Rsu1 | 5.56 | Genebody |
| **chr17** | Taf3 | -38.625 | Genebody |
| **chr17** | Trpc7 | -6.243 | Genebody |
| **chr18** | F8 | -23.697 | Genebody |
| **chr18** | F8 | 17.442 | Genebody |
| **chr18** | F8 | 18.015 | Genebody |
| **chr18** | Fem1c | -31.57 | Promoter |
| **chr18** | Il17b | 8.01 | Genebody |
| **chr18** | Nfatc1 | -9 | Genebody |

| | | | |
|---|---|---|---|
| **chr18** | Rnf165 | 6.772 | Genebody |
| **chr18** | St8sia5 | -19.34 | Genebody |
| **chr19** | Cdh13 | 5.932 | Genebody |
| **chr19** | Cmip | -39.082 | Genebody |
| **chr19** | Fto | -7.675 | Genebody |
| **chr19** | Mcm5 | -8.615 | Promoter |
| **chr19** | Mtss1l | 6.23 | Genebody |
| **chr19** | Mtss1l | 6.502 | Genebody |
| **chr19** | Zfp423 | 9.005 | Genebody |
| **chr19** | Zfpm1 | -11.9 | Genebody |
| **chr19** | Zfpm1 | -7.192 | Genebody |
| **chr2** | Adgrl2 | -5.193 | Genebody |
| **chr2** | Paqr6 | -40.93 | Genebody |
| **chr2** | Polr3g | -19.942 | Genebody |
| **chr2** | Prpf38b | -8.727 | Genebody |
| **chr2** | St6galnac3 | 6.19 | Genebody |
| **chr2** | St6galnac5 | 7.855 | Genebody |
| **chr2** | St6galnac5 | -5.193 | Genebody |
| **chr2** | Stpg2 | 5.675 | Promoter |
| **chr2** | Synpo2 | -15.628 | Genebody |
| **chr2** | Wdr70 | -37.635 | Genebody |
| **chr20** | Ank3 | 9.25 | Genebody |
| **chr20** | Anks1a | -6.795 | Genebody |
| **chr20** | C4a | -9.615 | Genebody |
| **chr20** | Cdkn1a | -21.295 | Genebody |
| **chr20** | Cisd1 | -12.375 | Genebody |
| **chr20** | Fyn | -6.875 | Genebody |
| **chr20** | Gnaz | -7.812 | Genebody |
| **chr20** | Grm4 | -9.322 | Genebody |
| **chr20** | Grm4 | 13.03 | Genebody |
| **chr20** | Hsf2bp | -5.328 | Genebody |
| **chr20** | Notch4 | -9.615 | Genebody |
| **chr20** | Pi16 | 13.03 | Genebody |
| **chr20** | Popdc3 | -18.58 | Genebody |
| **chr20** | Ppil1 | -19.31 | Genebody |
| **chr20** | Ppil1 | -9.322 | Genebody |
| **chr20** | Prep | -30.547 | Genebody |
| **chr20** | Prrc2a | -7.073 | Genebody |
| **chr20** | Stk19 | -9.615 | Genebody |
| **chr3** | Bcl2l11 | 6.66 | ıebody; Prom |
| **chr3** | Blcap | 21.09 | Promoter |
| **chr3** | Bmp7 | -26.863 | Genebody |

| | | | |
|---|---|---|---|
| **chr3** | Bpifb4 | -32.242 | Genebody |
| **chr3** | Cir1 | -22.988 | Genebody |
| **chr3** | Col5a1 | 7.585 | Genebody |
| **chr3** | Ebf4 | -6.353 | Genebody |
| **chr3** | Fam110a | 22.335 | Genebody |
| **chr3** | Fnbp1 | 15.312 | Genebody |
| **chr3** | Gnas | 7.277 | Genebody |
| **chr3** | Gnas | 8.758 | Promoter |
| **chr3** | LOC499742 | 27 | Genebody |
| **chr3** | LOC689618 | -9.21 | Genebody |
| **chr3** | Metap1d | 5.955 | Genebody |
| **chr3** | Miga2 | -38.475 | Genebody |
| **chr3** | Mir219-2 | 22.66 | Promoter |
| **chr3** | Mir2964 | 22.66 | Promoter |
| **chr3** | Nfatc2 | -14.34 | Genebody |
| **chr3** | Nnat | 21.09 | Promoter |
| **chr3** | Pax6 | -5.957 | Genebody |
| **chr3** | Plcb4 | -34.733 | Genebody |
| **chr3** | Pmepa1 | -28.773 | Promoter |
| **chr3** | Ppp1r16b | 19.28 | Genebody |
| **chr4** | Atg7 | -19.07 | Genebody |
| **chr4** | Cacna1c | -15.172 | Genebody |
| **chr4** | Foxp1 | -5.217 | Genebody |
| **chr4** | Gata2 | -5.62 | Promoter |
| **chr4** | Itpr2 | -17.982 | Genebody |
| **chr4** | Mest | -24.865 | Promoter |
| **chr4** | P3h3 | -8.125 | Promoter |
| **chr4** | Pik3c2g | 5.845 | Genebody |
| **chr4** | Rassf8 | -45 | Genebody |
| **chr4** | St8sia1 | -6.415 | Genebody |
| **chr5** | Akap2 | -6.85 | Genebody |
| **chr5** | Camta1 | 31.675 | Genebody |
| **chr5** | Car9 | 18.468 | Genebody |
| **chr5** | Ctnnbip1 | -8.142 | Genebody |
| **chr5** | Epb41l4b | -17.38 | Genebody |
| **chr5** | Fam46b | -12.948 | Genebody |
| **chr5** | Kazn | -7.262 | Genebody |
| **chr5** | LOC298116 | -8.125 | Genebody |
| **chr5** | Map7d1 | -13.755 | Genebody |
| **chr5** | Mmel1 | -29.215 | Genebody |
| **chr5** | Padi3 | -18.518 | Genebody |
| **chr5** | Paqr7 | -10.705 | Promoter |

| | | | |
|---|---|---|---|
| **chr5** | Per3 | 8.693 | Genebody |
| **chr5** | Ptpru | -26.237 | Genebody |
| **chr5** | RGD1566134 | -8.125 | Genebody |
| **chr5** | Rgs3 | -8.432 | Genebody |
| **chr5** | Snip1 | 29.058 | Genebody |
| **chr5** | Sytl1 | -9.207 | Genebody |
| **chr5** | Tln1 | 25.835 | Genebody |
| **chr5** | Zfp362 | -29.075 | Genebody |
| **chr5** | Zfp37 | -8.125 | Genebody |
| **chr5** | Zmym4 | -10.95 | Promoter |
| **chr6** | Bcl11b | -7.875 | Genebody |
| **chr6** | Btbd7 | 5.7 | Genebody |
| **chr6** | Dlk1 | 5.307 | Genebody |
| **chr6** | Ptpn21 | -17.69 | Promoter |
| **chr6** | Slc8a1 | 23.538 | Genebody |
| **chr6** | Tnfaip2 | 10.62 | Genebody |
| **chr6** | Twist1 | 6.585 | Promoter |
| **chr6** | Unc79 | -17.035 | Genebody |
| **chr7** | Amigo2 | 15.67 | Genebody |
| **chr7** | Apobec3b | -8.55 | Promoter |
| **chr7** | Arhgap39 | 29.44 | Genebody |
| **chr7** | Arhgap45 | -41.638 | Genebody |
| **chr7** | Brd1 | -18.925 | Promoter |
| **chr7** | Cacna1i | -6.178 | Genebody |
| **chr7** | Ccdc166 | 6.222 | Genebody |
| **chr7** | Dyrk2 | 15.5 | Genebody |
| **chr7** | Fam109b | -20.99 | Genebody |
| **chr7** | Fam19a5 | 12.027 | Genebody |
| **chr7** | Fzd6 | -34.582 | Genebody |
| **chr7** | Grin3b | 11.837 | Genebody |
| **chr7** | Igfbp6 | -12.107 | Promoter |
| **chr7** | Kcnh3 | -32.36 | Genebody |
| **chr7** | Krt5 | -6.637 | Genebody |
| **chr7** | Med16 | -41.638 | Promoter |
| **chr7** | MGC94207 | 29.44 | Promoter |
| **chr7** | Odf3l2 | -21.232 | Genebody |
| **chr7** | Osbpl8 | -7.49 | Promoter |
| **chr7** | Palm | 11.837 | Genebody |
| **chr7** | Polr3b | -44.813 | Genebody |
| **chr7** | Spats2 | -32.36 | Genebody |
| **chr7** | Tmprss9 | 12.28 | Genebody |
| **chr7** | Tob2 | -6.99 | Genebody |

| | | | |
|---|---|---|---|
| **chr8** | Ccrl2 | 14.477 | Promoter |
| **chr8** | Ctdspl | 8.105 | Genebody |
| **chr8** | Icam4 | -12.475 | Promoter |
| **chr8** | Nprl2 | -24.455 | Genebody |
| **chr8** | Rasgrf1 | 19.073 | Genebody |
| **chr8** | Smarcc1 | 10.615 | Promoter |
| **chr8** | Spsb4 | 29.202 | Genebody |
| **chr8** | Tmod3 | -9.308 | Genebody |
| **chr8** | Tmprss13 | -42.615 | Genebody |
| **chr8** | Trak1 | 7.5 | Promoter |
| **chr8** | Trim71 | -5.1 | Promoter |
| **chr8** | Uqcrc1 | 7.83 | Genebody |
| **chr8** | Zmynd10 | -24.455 | Promoter |
| **chr9** | Agap1 | 5.713 | Genebody |
| **chr9** | Ptk7 | 6.57 | Genebody |
| **chr9** | Sema6b | -13.07 | Genebody |
| **chr9** | Slc9a4 | 8.827 | Genebody |
| **chr9** | Sphkap | -38.07 | Genebody |
| **chr9** | Trerf1 | -20.207 | Promoter |
| **chr9** | Ube2f | -20.33 | Genebody |
| **chr9** | Wash1 | 5.69 | Genebody |
| **chrX** | Abcd1 | -14.77 | Promoter |
| **chrX** | Acsl4 | -5.968 | Promoter |
| **chrX** | Aifm1 | -8.35 | Promoter |
| **chrX** | Apln | -6.262 | Promoter |
| **chrX** | Arx | -9.138 | Genebody |
| **chrX** | Arx | -5.05 | Promoter |
| **chrX** | Atp2b3 | -12.972 | Genebody |
| **chrX** | Atp6ap2 | -6.217 | Promoter |
| **chrX** | Atp6ap2 | -5.09 | Promoter |
| **chrX** | Bcap31 | -14.77 | Promoter |
| **chrX** | Bcap31 | -5.005 | Promoter |
| **chrX** | Bcor | -6.812 | Genebody |
| **chrX** | Bcor | -8.11 | Promoter |
| **chrX** | Bex2 | -8.58 | Promoter |
| **chrX** | Bex3 | -7.908 | Promoter |
| **chrX** | Cacna1f | -5.215 | Genebody |
| **chrX** | Cask | -10.768 | Promoter |
| **chrX** | Ccdc22 | -10.825 | Promoter |
| **chrX** | Cdx4 | 8.743 | Promoter |
| **chrX** | Chrdl1 | -6.045 | Promoter |
| **chrX** | Chst7 | -8.943 | Promoter |

| | | | |
|---|---|---|---|
| **chrX** | Chst7 | -6.123 | Promoter |
| **chrX** | Cited1 | -7.333 | Genebody |
| **chrX** | Cited1 | -12.33 | Promoter |
| **chrX** | Ctps2 | -11.612 | Genebody |
| **chrX** | Dlg3 | -9.725 | Genebody |
| **chrX** | Dock11 | -24.54 | Promoter |
| **chrX** | Dusp9 | -7.627 | Promoter |
| **chrX** | Eda | -8.325 | Promoter |
| **chrX** | Eda | 5.605 | Promoter |
| **chrX** | Efnb1 | -7.655 | Genebody |
| **chrX** | Efnb1 | -5.425 | Promoter |
| **chrX** | Efnb1 | -7.665 | Promoter |
| **chrX** | Egfl6 | -5.012 | Promoter |
| **chrX** | Fam199x | -8.428 | Promoter |
| **chrX** | Fgf16 | -14.765 | Promoter |
| **chrX** | Gabre | -22.615 | Genebody |
| **chrX** | Gabre | 7.603 | Promoter |
| **chrX** | Gemin8 | -8.783 | Promoter |
| **chrX** | Gjb1 | -16.56 | Genebody |
| **chrX** | Gk | -9.5 | Promoter |
| **chrX** | Gnl3l | -10.412 | Promoter |
| **chrX** | Gprasp1 | -6.303 | Promoter |
| **chrX** | Gria3 | 6.033 | Promoter |
| **chrX** | Gspt2 | -14.07 | Promoter |
| **chrX** | Hccs | -11.443 | Promoter |
| **chrX** | Hcfc1 | -14.565 | Promoter |
| **chrX** | Hs6st2 | -9.02 | Genebody |
| **chrX** | Htatsf1 | -5.308 | Promoter |
| **chrX** | Htr2c | -6.175 | Promoter |
| **chrX** | Htr2c | -6.098 | Promoter |
| **chrX** | Idh3g;Ssr4 | -5.015 | Promoter |
| **chrX** | Iqsec2 | -7.763 | Promoter |
| **chrX** | Klhl15 | -20.252 | Promoter |
| **chrX** | Lage3 | -5.97 | Promoter |
| **chrX** | Lamp2 | -7.35 | Promoter |
| **chrX** | Lancl3 | -14.728 | Promoter |
| **chrX** | Lonrf3 | -18.26 | Promoter |
| **chrX** | Maged1 | -7.38 | Promoter |
| **chrX** | Map7d3 | -7.755 | Promoter |
| **chrX** | Mbtps2 | -13.175 | Promoter |
| **chrX** | Mbtps2 | -10.063 | Promoter |
| **chrX** | Mecp2 | -5.84 | Promoter |

| | | | |
|---|---|---|---|
| **chrX** | Med12 | -7.345 | Promoter |
| **chrX** | Med14 | -5.82 | Promoter |
| **chrX** | Med14 | 9.02 | Promoter |
| **chrX** | Mid1ip1 | -8.139 | Genebody |
| **chrX** | Mid1ip1 | -8.308 | Promoter |
| **chrX** | Mid1ip1 | -5.92 | Promoter |
| **chrX** | Mid2 | -9.782 | Promoter |
| **chrX** | Mid2 | -5.793 | Promoter |
| **chrX** | Mir3573 | -7.655 | Promoter |
| **chrX** | Mir3573 | -7.665 | Promoter |
| **chrX** | Ndufb11 | -5.608 | Promoter |
| **chrX** | Nhs | -7.022 | Genebody |
| **chrX** | Nhs | -5.795 | Genebody |
| **chrX** | Nhs | -18.183 | Promoter |
| **chrX** | Nhs | -12.49 | Promoter |
| **chrX** | Nhs | -10.023 | Promoter |
| **chrX** | Nono | 10.495 | Promoter |
| **chrX** | Nrk | -7.008 | Promoter |
| **chrX** | Nsdhl | -7.812 | Promoter |
| **chrX** | Nyx | -11.378 | Genebody |
| **chrX** | Ogt | -5.297 | Promoter |
| **chrX** | Ophn1 | 5.128 | Promoter |
| **chrX** | Otud5 | -6.692 | Promoter |
| **chrX** | Pak3 | -6.15 | Promoter |
| **chrX** | Pcdh19 | -7.107 | Promoter |
| **chrX** | Pcdh19 | -5.188 | Promoter |
| **chrX** | Pdzd11 | -6.838 | Promoter |
| **chrX** | Pgk1 | -6.553 | Promoter |
| **chrX** | Pih1d3 | -5.687 | Promoter |
| **chrX** | Pir | -10.24 | Promoter |
| **chrX** | Pja1 | -6.393 | Promoter |
| **chrX** | Plp2 | -13.295 | Promoter |
| **chrX** | Plxna3 | -7.408 | Promoter |
| **chrX** | Pou3f4 | -5.658 | Promoter |
| **chrX** | Ptchd1 | -6.513 | Promoter |
| **chrX** | Rai2 | -14.507 | Promoter |
| **chrX** | Rap2c | -11.463 | Promoter |
| **chrX** | Rbm10 | -5.608 | Promoter |
| **chrX** | Rbm3 | -5.255 | Promoter |
| **chrX** | RGD1562161 | -16.255 | Promoter |
| **chrX** | RGD1562161 | -7.84 | Promoter |
| **chrX** | RGD1562871 | -49.167 | Promoter |

| | | | |
|---|---|---|---|
| **chrX** | RGD1564541 | -8.025 | Promoter |
| **chrX** | Rnf128 | -9.75 | Promoter |
| **chrX** | Rps4x | -9.25 | Promoter |
| **chrX** | Rps6ka3 | -8.055 | Promoter |
| **chrX** | Rps6ka6 | -15.803 | Genebody |
| **chrX** | Sat1 | -7.398 | Promoter |
| **chrX** | 05-Sep | -6.062 | Promoter |
| **chrX** | Sh3bgrl | -7.125 | Promoter |
| **chrX** | Sh3kbp1 | -5.698 | Promoter |
| **chrX** | Shroom2 | -5.472 | Promoter |
| **chrX** | Shroom4 | -13.568 | Promoter |
| **chrX** | Slc16a2 | -6.075 | Genebody |
| **chrX** | Slc16a2 | -6.168 | Promoter |
| **chrX** | Slc7a3 | -9.42 | Promoter |
| **chrX** | Sms | -10.12 | Promoter |
| **chrX** | Sowahd | -8.795 | Promoter |
| **chrX** | Ssr4 | -5.015 | Promoter |
| **chrX** | Stag2 | -13.082 | Promoter |
| **chrX** | Tfe3 | -6.115 | Promoter |
| **chrX** | Thoc2 | -10.548 | Promoter |
| **chrX** | Thoc2 | -8.11 | Promoter |
| **chrX** | Tmem164 | -6.915 | Promoter |
| **chrX** | Tmsb4x | -6.285 | Promoter |
| **chrX** | Tsc22d3 | -19.297 | Promoter |
| **chrX** | Txlng | -16.23 | Promoter |
| **chrX** | Txlng | -5.552 | Promoter |
| **chrX** | Txlng | -5.382 | Promoter |
| **chrX** | Uba1 | -11.959 | Promoter |
| **chrX** | Ube2a | -8.025 | Promoter |
| **chrX** | Ubl4a | -6.505 | Promoter |
| **chrX** | Upf3b | -8.392 | Promoter |
| **chrX** | Usp51 | -8.42 | Promoter |
| **chrX** | Usp9x | -11.412 | Promoter |
| **chrX** | Usp9x | -9.325 | Promoter |
| **chrX** | Usp9x | -6.7 | Promoter |
| **chrX** | Utp14a | -14.135 | Promoter |
| **chrX** | Wdr45 | -5.435 | Promoter |
| **chrX** | Zdhhc15 | -12.722 | Promoter |
| **chrX** | Zdhhc9 | -11.417 | Promoter |
| **chrX** | Zfp275 | -13.72 | Promoter |
| **chrX** | Zic3 | -7.059 | Genebody |
| **chrX** | Abcd1 | -5.005 | Promoter |

**Supplementary Table 3: List of annotated genes to the DMRs at P90 along with their chromosome location, percentage of methylation differences between mHFD and control animals, and the location of gene regulatory region that the DMR spans**

| Chromosome | Gene symbol | tion differen | Location |
|---|---|---|---|
| **chr1** | Abcc8 | -6.595 | Genebody |
| **chr1** | Aig1 | 8.333 | Genebody |
| **chr1** | Apba1 | -14.462 | Genebody |
| **chr1** | Aqp11 | 8.84 | Genebody |
| **chr1** | Ascl3 | -12.288 | Promoter |
| **chr1** | Cdhr5 | -5.55 | Genebody |
| **chr1** | Coro1a | -20.045 | Genebody |
| **chr1** | Cyp2s1 | -6.59 | Genebody |
| **chr1** | Dagla | -5.227 | Genebody |
| **chr1** | Dpy19l3 | -21.363 | Genebody |
| **chr1** | Ebf3 | 15.64 | Genebody |
| **chr1** | Eps8l2 | 11.52 | Genebody |
| **chr1** | Fam120b | -13.665 | Genebody |
| **chr1** | Golga7b | -6.107 | Genebody |
| **chr1** | Gpam | 5.443 | Genebody |
| **chr1** | Gprc5b | -7.578 | Genebody |
| **chr1** | Hps1 | 5.235 | Genebody |
| **chr1** | Igf1r | -17.015 | Genebody |
| **chr1** | Igf2r | 16.578 | Genebody |
| **chr1** | Irf2bp1 | 5.522 | Genebody |
| **chr1** | Irgq | -16.468 | Genebody |
| **chr1** | Katna1 | -7.598 | Genebody |
| **chr1** | Klhl25 | -10.933 | Genebody |
| **chr1** | Lmo1 | -8.703 | Genebody |
| **chr1** | LOC310926 | -26.258 | Genebody |
| **chr1** | LOC310926 | -10.295 | Genebody |
| **chr1** | Ltbp3 | 15.45 | Genebody |
| **chr1** | Ltbp3 | 19.312 | Genebody |
| **chr1** | Ltbp4 | 6.427 | Promoter |
| **chr1** | Mpp1 | 27.897 | Genebody |
| **chr1** | Pank1 | -16.953 | Promoter |
| **chr1** | Park2 | 6.63 | Genebody |
| **chr1** | Pde8a | 24.095 | Genebody |
| **chr1** | Pold1 | -34.733 | Genebody |
| **chr1** | Prkg1 | -5.97 | Genebody |
| **chr1** | Rbm4 | -18.115 | Genebody |
| **chr1** | Serpinh1 | -10.116 | Genebody |
| **chr1** | Slc6a5 | -9.883 | Genebody |
| **chr1** | Slc9a3 | -5.777 | Genebody |
| **chr1** | Sox6 | -25.15 | Genebody |
| **chr1** | Sult2b1 | -10.723 | Genebody |
| **chr1** | Tacc2 | 22.7 | Genebody |
| **chr1** | Tbc1d10b | -30.145 | Promoter |

| | | | |
|---|---|---|---|
| **chr1** | Tenm4 | -5.405 | Genebody |
| **chr1** | Tph1 | 7.385 | Genebody |
| **chr1** | Usf2 | -11.69 | Genebody |
| **chr1** | Ust | -6.825 | Genebody |
| **chr1** | Uvrag | -7.55 | Genebody |
| **chr10** | Adam19 | -27.29 | Genebody |
| **chr10** | Anxa6 | 34.822 | Genebody |
| **chr10** | Axin2 | -6.66 | Genebody |
| **chr10** | C1qtnf2 | -8.015 | Promoter |
| **chr10** | Cbx2 | -8.793 | Genebody |
| **chr10** | Ccdc182 | -14.533 | Promoter |
| **chr10** | Ccdc40 | -12.98 | Genebody |
| **chr10** | Ccdc40 | -6.577 | Genebody |
| **chr10** | Ccdc40 | -8.793 | Genebody |
| **chr10** | Cnp | 13.48 | Genebody |
| **chr10** | Flt4 | 6.617 | Genebody |
| **chr10** | Gucy2e | 6.495 | Genebody |
| **chr10** | Kcnh4 | 10.067 | Genebody |
| **chr10** | Kdm6b | 23.235 | Genebody |
| **chr10** | Ksr1 | -7.88 | Genebody |
| **chr10** | Mgat5b | -10.167 | Genebody |
| **chr10** | Myh10 | -11.898 | Genebody |
| **chr10** | Nat9 | 6.02 | Promoter |
| **chr10** | Ncor1 | -9.12 | Genebody |
| **chr10** | Ogfod3 | -10.615 | Genebody |
| **chr10** | Pdlim4 | 10.647 | Genebody |
| **chr10** | Prss41 | 7.285 | Genebody |
| **chr10** | Rai1 | -6.309 | Genebody |
| **chr10** | Rgs9 | 8.95 | Genebody |
| **chr10** | Rundc3a | -5.417 | Genebody |
| **chr10** | Sdk2 | -16.935 | Genebody |
| **chr10** | Sdk2 | -5.785 | Genebody |
| **chr10** | Sdk2 | 8.055 | Genebody |
| **chr10** | Sept9; | -5.095 | Genebody |
| **chr10** | Slc9a3r1 | -6.983 | Genebody |
| **chr10** | Stc2 | -13.37 | Genebody |
| **chr10** | Stx8 | 6.9 | Genebody |
| **chr10** | Tenm2 | 11.643 | Genebody |
| **chr10** | Tmem104 | 6.02 | Genebody |
| **chr10** | Trim41 | -15.245 | Genebody |
| **chr10** | Wrap53 | -19.07 | Genebody |
| **chr11** | Arhgap31 | -17.18 | Genebody |
| **chr11** | Cd47 | 5.533 | Genebody |
| **chr11** | Dscam | 6.227 | Genebody |
| **chr11** | Gpr156 | 13.023 | Genebody |
| **chr11** | Gtf2e1 | -6.048 | Genebody |
| **chr11** | Hic2 | -5.325 | Genebody |
| **chr11** | Igf2bp2 | -14.723 | Genebody |
| **chr11** | Itgb5 | 6.597 | Genebody |

| | | | |
|---|---|---|---|
| **chr11** | Map3k7cl | -8.613 | Promoter |
| **chr11** | Mylk | 9.172 | Genebody |
| **chr11** | Tiam1 | 11.697 | Genebody |
| **chr11** | Tiam1 | 15.963 | Genebody |
| **chr11** | Tiam1 | -8.893 | Promoter |
| **chr11** | Vwa5b2 | 12.407 | Genebody |
| **chr12** | Arhgef18 | -48.002 | Genebody |
| **chr12** | Asphd2 | -17.92 | Genebody |
| **chr12** | Cdk2ap1 | -14.905 | Genebody |
| **chr12** | Cit | 6.54 | Genebody |
| **chr12** | Cyp3a2 | 6.545 | Genebody |
| **chr12** | Ddx54 | -15.262 | Genebody |
| **chr12** | Fbxo24 | -5.775 | Promoter |
| **chr12** | Flt1 | 6.402 | Genebody |
| **chr12** | Fry | -8.047 | Genebody |
| **chr12** | Gatsl2 | -7.592 | Genebody |
| **chr12** | Mn1 | -11.355 | Genebody |
| **chr12** | Nos1 | 21.438 | Genebody |
| **chr12** | P2rx7 | -6.955 | Promoter |
| **chr12** | Papolb | 6.822 | Promoter |
| **chr12** | Radil | 6.822 | Genebody |
| **chr12** | RGD156231( | -11.29 | Genebody |
| **chr12** | Sgsm1 | -29.753 | Genebody |
| **chr12** | Slc7a1 | -13.85 | Genebody |
| **chr12** | Tex26 | 8.345 | Genebody |
| **chr12** | Tmem132b | 40.185 | Genebody |
| **chr12** | Tpst2 | -15.268 | Genebody |
| **chr12** | Vps37b | -7.645 | Genebody |
| **chr13** | Ccdc93 | -7.52 | Genebody |
| **chr13** | Cfap45 | -19.012 | Genebody |
| **chr13** | Crb1 | -15.995 | Genebody |
| **chr13** | Degs1 | 29.87 | Genebody |
| **chr13** | Dpt | 26.918 | Genebody |
| **chr13** | Esrrg | -20.18 | Genebody |
| **chr13** | Glul | -5.157 | Genebody |
| **chr13** | Hmcn1 | -13.01 | Genebody |
| **chr13** | Igsf8 | 19.797 | Genebody |
| **chr13** | Itpkb | -10.92 | Genebody |
| **chr13** | Itpkb | -7.405 | Genebody |
| **chr13** | Pm20d1 | 8.515 | Genebody |
| **chr13** | Ptpn14 | -31.501 | Genebody |
| **chr13** | Ralb | -8.21 | Genebody |
| **chr13** | RGD1310587 | -9.918 | Promoter |
| **chr13** | Sec16b | 7.582 | Genebody |
| **chr13** | Tdrd5 | -6.33 | Promoter |
| **chr13** | Tor1aip2 | 5.865 | Genebody |
| **chr14** | Abcg3l2 | 11.728 | Genebody |
| **chr14** | Abcg3l4 | 11.728 | Genebody |
| **chr14** | Aff1 | 31.04 | Genebody |

| | | | |
|---|---|---|---|
| **chr14** | Ccni | -8.09 | Promoter |
| **chr14** | Cds1 | 14.87 | Genebody |
| **chr14** | Commd1 | 7.567 | Genebody |
| **chr14** | Ehbp1 | -23.478 | Genebody |
| **chr14** | Hmgb3 | 7.248 | Promoter |
| **chr14** | Kit | -7.708 | Genebody |
| **chr14** | Limk2 | -1.19 | Genebody |
| **chr14** | Mtmr3 | 17.71 | Genebody |
| **chr14** | Rab5a | -7.975 | Promoter |
| **chr14** | Rhbdd3 | 8.327 | Genebody |
| **chr14** | Rn28s | -8.075 | Genebody |
| **chr14** | Rn45s | -8.075 | Genebody |
| **chr14** | Slain2 | 14.425 | Genebody |
| **chr14** | Slc4a4 | -6.42 | Genebody |
| **chr14** | Spred2 | -5.652 | Genebody |
| **chr14** | Spred2 | 8.852 | Genebody |
| **chr14** | Tgfbr3 | -6.445 | Genebody |
| **chr14** | Thegl | -35.277 | Genebody |
| **chr14** | Tmem150c | -11.46 | Genebody |
| **chr14** | Tmem150c | -6.36 | Genebody |
| **chr14** | Zrsr1 | 7.567 | Promoter |
| **chr15** | Cd99l2 | -7.205 | Genebody |
| **chr15** | Cpne6 | 7.05 | Genebody |
| **chr15** | Dupd1 | -8.108 | Genebody |
| **chr15** | Ednrb | -9.507 | Genebody |
| **chr15** | Gpc5 | -11.238 | Genebody |
| **chr15** | Kcnk5 | -7.465 | Genebody |
| **chr15** | Klf5 | -15.1 | Genebody |
| **chr15** | Mycbp2 | -13.535 | Genebody |
| **chr15** | Neil2 | -6.242 | Promoter |
| **chr15** | Pcdh9 | 6.998 | Genebody |
| **chr15** | Peli2 | -13.765 | Genebody |
| **chr15** | Samd4a | -22.715 | Promoter |
| **chr15** | Sgcg | -12.105 | Genebody |
| **chr15** | Spata13 | -7.35 | Genebody |
| **chr15** | Spert | -8.685 | Genebody |
| **chr16** | Ank1 | -30.753 | Genebody |
| **chr16** | Arhgap22 | 10.285 | Genebody |
| **chr16** | Atp7b | -5.655 | Genebody |
| **chr16** | Cacna1d | -21.35 | Genebody |
| **chr16** | Cacna1d | -13.54 | Genebody |
| **chr16** | Cacna2d3 | -14.392 | Genebody |
| **chr16** | Cul4a | 21.477 | Genebody |
| **chr16** | Dlc1 | -6.41 | Genebody |
| **chr16** | Eaf1 | -9.692 | Genebody |
| **chr16** | Nt5dc2 | -6.58 | Promoter |
| **chr16** | Plvap | 5.117 | Promoter |
| **chr16** | Rasa3 | 15.053 | Genebody |
| **chr16** | Sh3bp5 | -25.882 | Genebody |

| | | | |
|---|---|---|---|
| **chr16** | Sh3rf1 | -5.705 | Genebody |
| **chr16** | Sh3rf1 | 27.622 | Genebody |
| **chr16** | Slc27a1 | -31.225 | Genebody |
| **chr16** | Tenm3 | -11.235 | Genebody |
| **chr16** | Tnks | -5.058 | Genebody |
| **chr17** | Actn2 | -13.245 | Genebody |
| **chr17** | Arhgap21 | 10.037 | Genebody |
| **chr17** | Atxn1 | -26.655 | Genebody |
| **chr17** | Fam193b | 10.752 | Genebody |
| **chr17** | Fam50b | 7.26 | Genebody |
| **chr17** | Gfod1 | 5.535 | Genebody |
| **chr17** | Hist1h2aa | 20.515 | Promoter |
| **chr17** | Hist1h2ba | 20.515 | Promoter |
| **chr17** | Hivep1 | 5.268 | Genebody |
| **chr17** | Lman2 | 14.858 | Genebody |
| **chr17** | Msrb2 | 33.997 | Genebody |
| **chr17** | Ntrk2 | -13.502 | Genebody |
| **chr17** | Phactr1 | -6.115 | Genebody |
| **chr17** | Prpf18 | -29.948 | Genebody |
| **chr17** | Rreb1 | 6.052 | Genebody |
| **chr17** | Rsu1 | 7.537 | Genebody |
| **chr17** | Spag6l | -7.282 | Genebody |
| **chr17** | Tgfbi | -23.26 | Genebody |
| **chr17** | Txndc5 | -7.29 | Genebody |
| **chr18** | Cdh2 | -6.202 | Genebody |
| **chr18** | Cep120 | -9.367 | Promoter |
| **chr18** | Impact | 9.805 | Promoter |
| **chr18** | Mapk4 | 6.548 | Genebody |
| **chr18** | Nedd4l | -22.642 | Genebody |
| **chr18** | Nedd4l | 5.452 | Genebody |
| **chr18** | Nfatc1 | -17.6 | Genebody |
| **chr18** | Nr3c1 | -13.148 | Genebody |
| **chr18** | St8sia5 | -11.197 | Genebody |
| **chr18** | Zfp532 | -15.973 | Promoter |
| **chr19** | Arhgap10 | 14.602 | Genebody |
| **chr19** | Fhod1 | 24.125 | Genebody |
| **chr19** | Fto | -12.6 | Genebody |
| **chr19** | Large1 | 6.6 | Genebody |
| **chr19** | Mvd | 6.607 | Genebody |
| **chr19** | Nf1x | -25.203 | Genebody |
| **chr19** | Nf1x | -5.37 | Genebody |
| **chr19** | Pard3 | 11.917 | Genebody |
| **chr19** | Tmem231 | -12.69 | Promoter |
| **chr19** | Tubb3 | 5.418 | Genebody |
| **chr19** | Zfp423 | 8.777 | Genebody |
| **chr2** | Adad1 | 35.98 | Promoter |
| **chr2** | Adgrl4 | -6.243 | Promoter |
| **chr2** | Arhgef28 | 15.132 | Genebody |
| **chr2** | C1qtnf3 | -6.965 | Promoter |

| | | | |
|---|---|---|---|
| **chr2** | Cd101 | -17.558 | Genebody |
| **chr2** | Cldn11 | -9.265 | Genebody |
| **chr2** | Jtb | 5.12 | Genebody |
| **chr2** | Kcnn3 | 5.043 | Promoter |
| **chr2** | Lifr | -6.64 | Genebody |
| **chr2** | Mab21l1 | -6.762 | Promoter |
| **chr2** | Maml3 | 13.457 | Genebody |
| **chr2** | Mecom | -5.727 | Genebody |
| **chr2** | Mgst2 | -8.425 | Promoter |
| **chr2** | Mrps21 | -13.082 | Genebody |
| **chr2** | Nfkb1 | -12.225 | Genebody |
| **chr2** | Noct | -8.377 | Genebody |
| **chr2** | Paqr6 | -11.022 | Genebody |
| **chr2** | Polr3g | 5.273 | Genebody |
| **chr2** | Prpf38b | 5.153 | Genebody |
| **chr2** | Sema4a | -10.852 | Genebody |
| **chr2** | Slc1a3 | 10.643 | Genebody |
| **chr2** | St6galnac3 | -5.93 | Genebody |
| **chr2** | St6galnac5 | -8.552 | Genebody |
| **chr2** | Syde2 | -15.085 | Genebody |
| **chr2** | Synpo2 | -9.087 | Genebody |
| **chr2** | Vav3 | -5.743 | Genebody |
| **chr20** | Ank3 | -9.788 | Genebody |
| **chr20** | Ank3 | 16.072 | Genebody |
| **chr20** | Anks1a | 25.655 | Genebody |
| **chr20** | Atf6b | 24.162 | Genebody |
| **chr20** | Btbd9 | -16.209 | Genebody |
| **chr20** | C4a | 24.162 | Genebody |
| **chr20** | C4a | 16.977 | Promoter |
| **chr20** | Cbs | -6.967 | Genebody |
| **chr20** | Cdh23 | -26.372 | Genebody |
| **chr20** | Col6a2 | -19.052 | Genebody |
| **chr20** | Fkbp5 | 5.76 | Genebody |
| **chr20** | Fyn | -10.847 | Genebody |
| **chr20** | Gnaz | -9.422 | Genebody |
| **chr20** | Grm4 | -8.44 | Genebody |
| **chr20** | Kpna5 | -20.872 | Genebody |
| **chr20** | Lace1 | -10.083 | Genebody |
| **chr20** | Man1a1 | -8.088 | Genebody |
| **chr20** | Neu1 | -6.55 | Promoter |
| **chr20** | Pde9a | -17.427 | Genebody |
| **chr20** | Ppil1 | -8.44 | Genebody |
| **chr20** | Prrc2a | -20.488 | Genebody |
| **chr20** | Sobp | 5.327 | Genebody |
| **chr20** | Srgn | -12.302 | Genebody |
| **chr20** | Stk19 | 24.162 | Genebody |
| **chr20** | Stk19 | 16.977 | Genebody |
| **chr20** | Traf3ip2 | -21.758 | Genebody |
| **chr20** | Trappc10 | -26.56 | Genebody |

| | | | |
|---|---|---|---|
| **chr20** | Ube2d1 | 22.63 | Genebody |
| **chr20** | Zfp523 | 14.935 | Genebody |
| **chr3** | Baz2b | -21.668 | Genebody |
| **chr3** | Bmp7 | 20 | Genebody |
| **chr3** | Cass4 | -16.103 | Promoter |
| **chr3** | Cyp24a1 | -10.057 | Genebody |
| **chr3** | Fam83d | 6.35 | Genebody |
| **chr3** | Gins1 | -6.62 | Genebody |
| **chr3** | Gnas | 9.525 | Genebody |
| **chr3** | Gnas | 7.532 | Genebody |
| **chr3** | Gnas | 6.298 | Promoter |
| **chr3** | Gnas | 19.07 | nebody;promoter |
| **chr3** | Gnas | -7.058 | Promoter |
| **chr3** | Helz2 | -13.595 | Genebody |
| **chr3** | Kcnt1 | -18.467 | Genebody |
| **chr3** | LOC499742 | 34.943 | Genebody |
| **chr3** | Map1a | -11.137 | Genebody |
| **chr3** | Mertk | -7.864 | Genebody |
| **chr3** | Ncs1 | -10.582 | Genebody |
| **chr3** | Nkain4 | 5.662 | Promoter |
| **chr3** | Pck1 | -9.385 | Promoter |
| **chr3** | Ppp1r1c | -5.14 | Promoter |
| **chr3** | Prr5l | 5.06 | Promoter |
| **chr3** | Slc25a25 | -13.762 | Genebody |
| **chr3** | Stom | 6.023 | Genebody |
| **chr3** | Tanc1 | -6.427 | Genebody |
| **chr3** | Tanc1 | -5.5 | Genebody |
| **chr3** | Tanc1 | 5.293 | Genebody |
| **chr3** | Tox2 | -20.05 | Genebody |
| **chr3** | Traf1 | 13.435 | Promoter |
| **chr4** | Add2 | -11.923 | Genebody |
| **chr4** | Asb10 | -6.853 | Promoter |
| **chr4** | Bcat1 | -13.025 | Genebody |
| **chr4** | Cacna1c | -15.688 | Genebody |
| **chr4** | Cacna1c | -7.46 | Genebody |
| **chr4** | Cd36 | -26.05 | Genebody |
| **chr4** | Cdk6 | -7.4 | Genebody |
| **chr4** | Cyp51 | -8.068 | Genebody |
| **chr4** | Foxp1 | -6.91 | Genebody |
| **chr4** | Gucy2c | -11.79 | Genebody |
| **chr4** | Itpr2 | -8.275 | Genebody |
| **chr4** | Magi1 | 34.558 | Genebody |
| **chr4** | Prrt4 | -6.925 | Genebody |
| **chr4** | Ptpro | -5.5 | Genebody |
| **chr4** | RGD1565355 | -26.05 | Genebody |
| **chr4** | Slc6a13 | 11.355 | Genebody |
| **chr4** | Sox5 | -9.75 | Genebody |
| **chr4** | Srpk2 | -25.458 | Genebody |
| **chr4** | Tmem140 | -5.998 | Genebody |

| | | | |
|---|---|---|---|
| **chr4** | Vwf | -5.53 | Genebody |
| **chr4** | Wdr91 | -28.515 | Genebody |
| **chr5** | Adgrb2 | -27.222 | Genebody |
| **chr5** | Arhgef19 | -12.488 | Genebody |
| **chr5** | Astn2 | -9.642 | Genebody |
| **chr5** | Bach2 | -10.555 | Genebody |
| **chr5** | Eif4g3 | -16.717 | Genebody |
| **chr5** | Ephb2 | -9.667 | Genebody |
| **chr5** | Ephb2 | -5.61 | Genebody |
| **chr5** | Fbxo6 | 5.645 | Promoter |
| **chr5** | Gabbr2 | 20.407 | Genebody |
| **chr5** | Kcnab2 | 14.257 | Genebody |
| **chr5** | Kcnb2 | 13.025 | Promoter |
| **chr5** | Map3k6 | 17.75 | Genebody |
| **chr5** | Melk | -14.575 | Genebody |
| **chr5** | Morn1 | 6.143 | Genebody |
| **chr5** | Murc | 12.377 | Promoter |
| **chr5** | Nfia | 17.115 | Genebody |
| **chr5** | Nphp4 | -17.625 | Genebody |
| **chr5** | Otud3 | -17.877 | Genebody |
| **chr5** | Palm2 | 29.563 | Genebody |
| **chr5** | Plekhg5 | 8.245 | Genebody |
| **chr5** | Rgs3 | -7.377 | Genebody |
| **chr5** | Scp2 | 9.677 | Genebody |
| **chr5** | Slc24a2 | 8.332 | Genebody |
| **chr5** | Slc2a5 | -6.237 | Promoter |
| **chr5** | Tmem51 | -11.998 | Genebody |
| **chr5** | Tmem51 | 8.293 | Genebody |
| **chr5** | Tnfrsf25 | 8.245 | Promoter |
| **chr5** | Zbtb40 | -25.34 | Genebody |
| **chr5** | Zfyve9 | -13.85 | Genebody |
| **chr6** | Adgrf3 | -22.505 | Genebody |
| **chr6** | Agmo | -9.08 | Genebody |
| **chr6** | Akap6 | 6.242 | Genebody |
| **chr6** | Crim1 | -14.285 | Genebody |
| **chr6** | Crim1 | -5.02 | Genebody |
| **chr6** | Dpf3 | 5.697 | Genebody |
| **chr6** | Dpysl5 | -8.352 | Genebody |
| **chr6** | Elmsan1 | -28.288 | Genebody |
| **chr6** | Eml1 | 10.013 | Genebody |
| **chr6** | Foxn3 | -25.62 | Genebody |
| **chr6** | Foxn3 | -6.613 | Genebody |
| **chr6** | Frmd6 | -10.86 | Genebody |
| **chr6** | Frmd6 | 5.8 | Genebody |
| **chr6** | Pcnx1 | -24.238 | Genebody |
| **chr6** | Prkce | -11.757 | Genebody |
| **chr6** | Prkce | -5.222 | Genebody |
| **chr6** | Prkch | -6.878 | Genebody |
| **chr6** | RGD1566401 | -17.043 | Genebody |

| | | | |
|---|---|---|---|
| **chr6** | Rgs6 | 11.893 | Genebody |
| **chr6** | Rin3 | -22.602 | Genebody |
| **chr6** | Slc25a29 | 15.123 | Genebody |
| **chr6** | Taf1b | 5.975 | Genebody |
| **chr6** | Traf3 | 12.838 | Genebody |
| **chr6** | Unc79 | -13.765 | Genebody |
| **chr7** | Acvrl1 | -11.875 | Genebody |
| **chr7** | Adcy8 | -7.753 | Promoter |
| **chr7** | Anks1b | -21.85 | Genebody |
| **chr7** | Ano6 | 7.955 | Genebody |
| **chr7** | Chst11 | -5.688 | Genebody |
| **chr7** | Csrnp2 | -23.935 | Genebody |
| **chr7** | Dnajc22 | 12.923 | Promoter |
| **chr7** | Fam19a5 | 7.757 | Genebody |
| **chr7** | Fam19a5 | 9.798 | Genebody |
| **chr7** | Gng7 | -7.64 | Genebody |
| **chr7** | Jsrp1 | -5.29 | Genebody |
| **chr7** | Midn | 21.398 | Genebody |
| **chr7** | Mlc1 | -14.425 | Promoter |
| **chr7** | Panx2 | 25.913 | Genebody |
| **chr7** | Pced1b | -6.995 | Genebody |
| **chr7** | Prr5 | 10.02 | Genebody |
| **chr7** | Rapgef3 | -58.645 | Genebody |
| **chr7** | Rassf3 | -12.738 | Genebody |
| **chr7** | Recql4 | -7.47 | Genebody |
| **chr7** | Slc38a1 | -20.31 | Genebody |
| **chr7** | Syde1 | 11.783 | Genebody |
| **chr7** | Tjp3 | -12.562 | Genebody |
| **chr7** | Tjp3 | -5.085 | Genebody |
| **chr7** | Tmprss9 | -10.725 | Genebody |
| **chr7** | Tmprss9 | -10.145 | Genebody |
| **chr7** | Tmprss9 | 6.857 | Genebody |
| **chr7** | Tob2 | -21.152 | Genebody |
| **chr8** | Acvr2b | 6.705 | Genebody |
| **chr8** | Angptl6 | -12.68 | Promoter |
| **chr8** | Bnip2 | -8.192 | Promoter |
| **chr8** | C1qtnf5 | -12.16 | Promoter |
| **chr8** | C1qtnf5 | -9.848 | Promoter |
| **chr8** | Cadm1 | -11.19 | Genebody |
| **chr8** | Cgnl1 | -13.222 | Genebody |
| **chr8** | Ctdspl | -14.162 | Genebody |
| **chr8** | Ctdspl | -6.155 | Genebody |
| **chr8** | Dscaml1 | -11.413 | Genebody |
| **chr8** | Ephb1 | -16.257 | Genebody |
| **chr8** | Gramd1b | 8.373 | Genebody |
| **chr8** | Gucy1a2 | 15.073 | Genebody |
| **chr8** | Icam4 | -15.715 | Promoter |
| **chr8** | Icam5 | -8.9 | Genebody |
| **chr8** | Kank2 | -9.303 | Genebody |

| | | | |
|---|---|---|---|
| **chr8** | Pth1r | 10.425 | Genebody |
| **chr8** | Tle3 | -10.873 | Genebody |
| **chr8** | Uaca | -5.495 | Genebody |
| **chr8** | Zbtb16 | 9.173 | Genebody |
| **chr9** | Adgrf5 | -6.877 | Genebody |
| **chr9** | Ankrd44 | 15.397 | Genebody |
| **chr9** | Arhgap28 | 17.975 | Genebody |
| **chr9** | Col4a3 | -10.542 | Genebody |
| **chr9** | Efna5 | -7.08 | Genebody |
| **chr9** | Hibch | 15 | Genebody |
| **chr9** | Hspd1 | 5.618 | Promoter |
| **chr9** | Hspe1 | 5.618 | Promoter |
| **chr9** | Klf7 | -15.447 | Promoter |
| **chr9** | Pard3b | 6.217 | Genebody |
| **chr9** | Pdcl3 | 9.117 | Genebody |
| **chr9** | Ppp1r7 | -6.053 | Genebody |
| **chr9** | Ptk7 | -11.27 | Genebody |
| **chr9** | Ptprm | -5.048 | Genebody |
| **chr9** | RGD1305645 | -5.242 | Promoter |
| **chr9** | Runx2 | -20.837 | Genebody |
| **chr9** | Spag16 | -16.91 | Genebody |
| **chr9** | Sult1c2a | -7.733 | Genebody |
| **chr9** | Tbc1d8 | -5.553 | Genebody |
| **chr9** | Tnfrsf21 | -20.322 | Genebody |
| **chrX** | Alg13 | 12.565 | Promoter |
| **chrX** | Ap1s2 | 9.072 | Promoter |
| **chrX** | Arx | 10.207 | Genebody |
| **chrX** | Arx | 14.237 | Genebody |
| **chrX** | Cnksr2 | 6.36 | Promoter |
| **chrX** | Dlg3 | -9.223 | Promoter |
| **chrX** | Dmd | -5.925 | Promoter |
| **chrX** | Dock11 | -6.973 | Promoter |
| **chrX** | Emd | -6.502 | Promoter |
| **chrX** | Fgf13 | -5.74 | Genebody |
| **chrX** | Htatsf1 | 14.31 | Promoter |
| **chrX** | Iqsec2 | 22.513 | Promoter |
| **chrX** | Lancl3 | -11.385 | Promoter |
| **chrX** | OC10255280 | 8.287 | Promoter |
| **chrX** | Mecp2 | -5.72 | Promoter |
| **chrX** | Ndufb11 | -17.19 | Promoter |
| **chrX** | Nhs | -18.252 | Genebody |
| **chrX** | Phka1 | -6.793 | Genebody |
| **chrX** | Plp1 | -18.373 | Genebody |
| **chrX** | Rbbp7 | -12.092 | Promoter |
| **chrX** | Rbm10 | -17.19 | Promoter |
| **chrX** | Slc25a5 | 21.335 | Promoter |
| **chrX** | Wdr45 | 6.607 | Promoter |

**Supplementary Table 4: Common GO terms of biological processes enriched by the genes identified from DMR analysis at P7 and P90 clustered into groups based on network analysis**

| Go ID | Go term | P7 FDR | P7 genes | P90 FDR | P90 genes |
|---|---|---|---|---|---|
| **Neuronal projection & nervous system development** | | | | | |
| **GO:0007399** | Nervous system development | 1.65E-04 | FYN,RUNX1,MARK1,DLG3,PCDH19,PAX6,PAK3,BCL11B,APP,BCL11A,THOC2,SLC8A1,FOXP1,LRRN2,TENM4,SRCIN1,GATA2,PTPRU,NAV2,EEF2K,BCL2L11,ACSL4,TIAM1,NNAT,TRIM71,RNF165,SHANK2,PLXNA3,ATP2B3 | 7.97E-04 | FYN,FRY,KIT,PLP1,DLG3,RGS9,PRKCH,PTPRO,EPHB1,NCS1,DPYSL5,FOXP1,EDNRB,TENM4,TNFRSF21,PTPRM,TENM3,TUBB3,CNP,NTRK2,TACC2,SOX6,ABCC8,TIAM1,ZBTB16,PARD3,KDM6B,PCDH9,FGF13,IMPACT,DMD |
| **GO:0007409** | Axonogenesis | 0.0433 | PAK3,APP,LRRN2,TIAM1,RNF165,PLXNA3 | 0.0409 | PTPRO,EPHB1,PTPRM,TUBB3,NTRK2,TIAM1,FGF13 |
| **GO:0022008** | Neurogenesis | 0.00312 | RUNX1,MARK1,PAX6,PAK3,BCL11B,APP,BCL11A,THOC2,LRRN2,TENM4,SRCIN1,GATA2,PTPRU,EEF2K,ACSL4,TIAM1,NNAT,RNF165,PLXNA3 | 0.00592 | FRY,KIT,PLP1,PRKCH,PTPRO,EPHB1,NCS1,DPYSL5,TENM4,TNFRSF21,PTPRM,TENM3,TUBB3,NTRK2,SOX6,ABCC8,TIAM1,PARD3,FGF13,IMPACT,DMD |
| **GO:0040011** | Locomotion | 0.00309 | FYN,MARK1,USP9X,SH3KBP1,PAX6,PAK3,APP,SLC8A1,SRCIN1,GATA2,GAS6,RHBDF1,TIAM1,SHROOM2,RNF165,PLXNA3 | 0.0292 | FYN,CD47,MYLK,KIT,SLC9A3R1,PTPRO,EPHB1,PTPRM,TUBB3,NTRK2,SEMA4A,ABCC8,TIAM1,PLEKHG5,FGF13 |
| **GO:0048666** | Neuron development | 0.0119 | PAX6,PAK3,APP,BCL11A,THOC2,LRRN2,TENM4,SRCIN1,EEF2K,ACSL4,TIAM1,RNF165,PLXNA3 | 0.0413 | FRY,PTPRO,EPHB1,NCS1,TENM4,PTPRM,TENM3,TUBB3,NTRK2,TIAM1,FGF13,IMPACT,DMD |
| **GO:0048699** | Generation of neurons | 0.00303 | RUNX1,MARK1,PAX6,PAK3,APP,BCL11A,THOC2,LRRN2,TENM4,SRCIN1,GATA2,PTPRU,EEF2K,ACSL4,TIAM1,NNAT,RNF165,PLXNA3 | 0.0158 | FRY,KIT,PRKCH,PTPRO,EPHB1,NCS1,DPYSL5,TENM4,TNFRSF21,PTPRM,TENM3,TUBB3,NTRK2,ABCC8,TIAM1,FGF13,IMPACT,DMD |
| **GO:0048812** | Neuron projection morphogenesis | 0.0111 | PAK3,APP,BCL11A,LRRN2,SRCIN1,EEF2K,TIAM1,RNF165,PLXNA3 | 0.0314 | PTPRO,EPHB1,PTPRM,TUBB3,NTRK2,TIAM1,FGF13,IMPACT,DMD |
| **GO:0048858** | Cell projection morphogenesis | 0.0112 | PAK3,APP,BCL11A,LRRN2,SRCIN1,EEF2K,TIAM1,RNF165,PLXNA3 | 0.0314 | PTPRO,EPHB1,PTPRM,TUBB3,NTRK2,TIAM1,FGF13,IMPACT,DMD |
| **GO:0048870** | Cell motility | 0.0112 | FYN,MARK1,USP9X,SH3KBP1,PAX6,PAK3,SLC8A1,SRCIN1,GATA2,GAS6,RHBDF1,TIAM1,SHROOM2 | 0.0381 | FYN,CD47,MYLK,KIT,SLC9A3R1,PTPRO,EPHB1,PTPRM,NTRK2,ABCC8,TIAM1,PLEKHG5,FGF13 |
| **GO:0120036** | Plasma membrane bounded cell projection organization | 0.0464 | PAX6,PAK3,APP,BCL11A,THOC2,LRRN2,SRCIN1,EEF2K,ACSL4,TIAM1,ZMYND10,RNF165,PLXNA3 | 0.0487 | FRY,P2RX7,KIT,PTPRO,EPHB1,NCS1,PTPRM,TUBB3,CDHR5,NTRK2,TMEM231,TIAM1,FGF13,IMPACT,DMD |
| **GO:0120039** | Plasma membrane bounded cell projection morphogenesis | 0.0112 | PAK3,APP,BCL11A,LRRN2,SRCIN1,EEF2K,TIAM1,RNF165,PLXNA3 | 0.0314 | PTPRO,EPHB1,PTPRM,TUBB3,NTRK2,TIAM1,FGF13,IMPACT,DMD |
| **Cellular morphogenesis & organism development** | | | | | |
| **GO:0000902** | Cell morphogenesis | 6.07E-04 | NOTCH4,SH3KBP1,PAK3,APP,BCL11A,LRRN2,TENM4,SRCIN1,ZMYM4,EEF2K,TIAM1,CSNK1D,RNF165,PLXNA3 | 0.0155 | FRY,PTPRO,EPHB1,TENM4,PTPRM,TENM3,TUBB3,NTRK2,TIAM1,FGF13,IMPACT,DMD |
| **GO:0016477** | Cell migration | 0.00676 | FYN,MARK1,USP9X,SH3KBP1,PAX6,PAK3,SLC8A1,SRCIN1,GATA2,GAS6,RHBDF1,TIAM1,SHROOM2 | 0.0458 | FYN,CD47,MYLK,KIT,PTPRO,EPHB1,PTPRM,NTRK2,ABCC8,TIAM1,PLEKHG5,FGF13 |
| **GO:0032989** | Cellular component morphogenesis | 4.63E-04 | NOTCH4,SH3KBP1,PAK3,APP,BCL11A,LRRN2,TENM4,SRCIN1,ZMYM4,EEF2K,TIAM1,TMOD3,CSNK1D,RNF165,PLXNA3 | 0.0275 | FRY,PTPRO,EPHB1,TENM4,PTPRM,TENM3,TUBB3,NTRK2,TIAM1,FGF13,IMPACT,DMD |
| **GO:0032990** | Cell part morphogenesis | 0.0112 | PAK3,APP,BCL11A,LRRN2,SRCIN1,EEF2K,TIAM1,RNF165,PLXNA3 | 0.0314 | PTPRO,EPHB1,PTPRM,TUBB3,NTRK2,TIAM1,FGF13,IMPACT,DMD |
| **GO:0045595** | Regulation of cell differentiation | 0.0389 | RUNX1,CITED1,PAX6,PAK3,BCL11A,THOC2,FOXP1,SRCIN1,GATA2,EEF2K,GAS6,TIAM1,PLXNA3 | 0.0428 | KIT,PRKCH,EPHB1,NCS1,KLF5,FOXP1,NOCT,TNFRSF21,NTRK2,ABCC8,TIAM1,ZBTB16,FGF13,IMPACT,DMD |

| | | | | | |
|---|---|---|---|---|---|
| **GO:0048468** | Cell development | 0.00365 | NOTCH4,CDKN1A,PAX6,PAK3,APP,BCL11A,THOC2,FOXP1,LRRN2,TENM4,SRCIN1,SMARCA2,GATA2,EEF2K,ACSL4,TIAM1,TMOD3,RNF165,BMP7,PLXNA3 | 0.00402 | FRY,KIT,PRKCH,PTPRO,EPHB1,NCS1,FOXP1,TENM4,TNFRSF21,PTPRM,TENM3,SGCG,HIST1H2BA,TUBB3,NTRK2,LIMK2,ABCC8,TIAM1,PARD3,FGF13,IMPACT,DMD,BMP7 |
| **GO:0048513** | Animal organ development | 0.0149 | CDKN1A,NCOR2,RUNX1,POU3F4,CDX4,CITED1,PAX6,SLC8A1,FOXP1,TWIST1,GATA2,PTPRU,RBM20,IGF2R,CTNNBIP1,BCL2L11,GAS6,SMARCC1,NNAT,NHS,ZFPM1,SHANK2,ATP2B3 | 0.0475 | SRGN,TGFBR3,KIT,PRKCH,PTPRO,EPHB1,KLF5,FOXP1,CCDC182,PTPRM,TGFBI,SGCG,IGF2R,CNP,NTRK2,CORO1A,TACC2,STC2,ZBTB16,NHS,KDM6B,PCDH9,FGF13,DMD |
| **GO:0050793** | Regulation of developmental process | 0.0206 | RUNX1,CITED1,SH3KBP1,PAX6,PAK3,APP,BCL11A,THOC2,FOXP1,SRCIN1,GATA2,ZMYM4,EEF2K,GAS6,TIAM1,CSNK1D,PLXNA3 | 0.0484 | SRGN,KIT,PRKCH,EPHB1,NCS1,KLF5,FOXP1,NOCT,TNFRSF21,PTPRM,NTRK2,ABCC8,TIAM1,ZBTB16,PARD3,FGF13,IMPACT,DMD |
| **GO:0051128** | Regulation of cellular component organization | 3.22E-04 | CIT,RABGAP1L,MID1IP1,APLN,SH3KBP1,PAX6,PAK3,BCL11A,THOC2,ATG7,FOXP1,SRCIN1,GATA2,ZMYM4,SYNPO2,EEF2K,TBC1D10B,GAS6,ANK1,TIAM1,NPRL2,EML2,CSNK1D,PLXNA3 | 0.00597 | SGSM1,CIT,P2RX7,CD47,KIT,PRKCH,RBBP7,EPHB1,NCS1,FOXP1,TBC1D8,SYNPO2,TBC1D10B,NEDD4L,CDHR5,ANK1,NTRK2,ABCC8,TIAM1,SLC25A5,FGF13,IMPACT,DMD |
| **GO:0051674** | Localization of cell | 0.0112 | FYN,MARK1,USP9X,SH3KBP1,PAX6,PAK3,SLC8A1,SRCIN1,GATA2,GAS6,RHBDF1,TIAM1,SHROOM2 | 0.0381 | FYN,CD47,MYLK,KIT,SLC9A3R1,PTPRO,EPHB1,PTPRM,NTRK2,ABCC8,TIAM1,PLEKHG5,FGF13 |
| **Protein transportation & secretion** | | | | | |
| **GO:0002790** | Peptide secretion | 0.0293 | APLN,PAX6,SRCIN1,GAS6,RHBDF1,TIAM1,NNAT | 0.0314 | SRGN,P2RX7,TNFRSF21,HSPD1,MIDN,ABCC8,TIAM1,GLUL |
| **GO:0002791** | Regulation of peptide secretion | 0.0173 | APLN,PAX6,SRCIN1,GAS6,RHBDF1,TIAM1,NNAT | 0.0199 | SRGN,P2RX7,TNFRSF21,HSPD1,MIDN,ABCC8,TIAM1,GLUL |
| **GO:0010646** | Regulation of cell communication | 0.00857 | FYN,RUNX1,RGS8,MAP3K14,APLN,PAX6,PAK3,APP,NYX,IGFBP6,CTNNBIP1,GAS6,DLK1,RHBDF1,TIAM1,NPRL2,NNAT,RNF165,SHANK2,SMOK2A,BMP7,IQSEC2 | 0.0428 | BTBD9,FYN,MAP3K7CL,TGFBR3,KIT,SLC9A3R1,RGS9,PRKCH,PTPRO,EPHB1,SLC24A2,MAP3K6,KSR1,MIDN,NTRK2,ABCC8,TIAM1,JSRP1,SLC25A5,GLUL,BMP7,IQSEC2 |
| **GO:0023051** | Regulation of signaling | 0.00509 | FYN,RUNX1,RGS8,MAP3K14,APLN,PAX6,PAK3,APP,NYX,IGFBP6,CTNNBIP1,GAS6,DLK1,RHBDF1,TIAM1,NPRL2,NNAT,FAM109B,RNF165,SHANK2,SMOK2A,BMP7,IQSEC2 | 0.0458 | BTBD9,FYN,MAP3K7CL,TGFBR3,KIT,SLC9A3R1,RGS9,PRKCH,PTPRO,EPHB1,SLC24A2,MAP3K6,KSR1,MIDN,NTRK2,ABCC8,TIAM1,JSRP1,SLC25A5,GLUL,BMP7,IQSEC2 |
| **GO:0033036** | Macromolecule localization | 0.0472 | RUNX1,RABGAP1L,DLG3,PDZD11,THOC2,COMMD1,TNFAIP2,SRCIN1,IGF2R,TBC1D10B,GAS6,ACSL4,RHBDF1,TIAM1,NNAT,UBL4A,BCAP31 | 0.00402 | SRGN,SGSM1,P2RX7,DLG3,SLC9A3R1,STX8,COMMD1,TNFRSF21,KCNAB2,EPHB2,TBC1D8,HSPD1,IGF2R,GOLGA7B,MIDN,TBC1D10B,PRPF18,ABCC8,TMEM231,TIAM1,ZBTB16,PARD3,MLC1,FGF13,GLUL |
| **GO:0051046** | Regulation of secretion | 0.0157 | RUNX1,APLN,PAX6,SYTL1,SRCIN1,GATA2,GAS6,RHBDF1,TIAM1,NNAT | 0.0261 | SRGN,P2RX7,NCS1,EDNRB,TNFRSF21,HSPD1,MIDN,NTRK2,ABCC8,TIAM1,GLUL |
| **GO:0051049** | Regulation of transport | 0.0304 | RUNX1,RABGAP1L,APLN,PAX6,SLC8A1,SYTL1,COMMD1,SRCIN1,GATA2,EEF2K,TBC1D10B,GAS6,RHBDF1,TIAM1,NNAT | 6.57E-04 | SRGN,SGSM1,P2RX7,CD47,ANO6,NCS1,COMMD1,EDNRB,TNFRSF21,KCNAB2,TBC1D8,HSPD1,MIDN,UVRAG,TBC1D10B,NEDD4L,NTRK2,ABCC8,TIAM1,NKAIN4,MLC1,JSRP1,SLC25A5,GLUL |
| **GO:1903530** | Regulation of secretion by cell | 0.0119 | RUNX1,APLN,PAX6,SYTL1,SRCIN1,GATA2,GAS6,RHBDF1,TIAM1,NNAT | 0.0365 | SRGN,P2RX7,NCS1,TNFRSF21,HSPD1,MIDN,NTRK2,ABCC8,TIAM1,GLUL |
| **Phosphorylation & catalytic regulation** | | | | | |
| **GO:0006468** | Protein phosphorylation | 0.0381 | FYN,MARK1,MAP3K14,PAK3,APP,NRK,NYX,SRCIN1,EEF2K,MAP3K4,GAS6,TIAM1,MAST3,CSNK1D,SMOK2A,BMP7 | 0.00402 | FYN,P2RX7,MAP3K7CL,KIT,SLC9A3R1,PRKCH,EPHB1,MAP3K6,CDK6,EDNRB,KSR1,MELK,JTB,MERTK,NTRK2,LIMK2,TIAM1,PPP1R1C,ACVRL1,PARD3,IMPACT,SH3BP5,BMP7 |
| **GO:0016310** | Phosphorylation | 0.0178 | FYN,MARK1,MAP3K14,APLN,PAK3,APP,NRK,NYX,SRCIN1,EEF2K,MAP3K4,GAS6,TIAM1,MAST3,UQCRC1,CSNK1D,SMOK2A,BMP7,PGK1 | 0.00698 | FYN,P2RX7,MAP3K7CL,KIT,SLC9A3R1,PRKCH,EPHB1,MAP3K6,CDK6,EDNRB,KSR1,MELK,MIDN,JTB,MERTK,NTRK2,LIMK2,TIAM1,PPP1R1C,ACVRL1,PARD3,IMPACT,SH3BP5,BMP7 |
| **GO:0043085** | Positive regulation of catalytic activity | 0.0148 | RGS8,RABGAP1L,MID1IP1,MAP3K14,PAK3,AIFM1,NRK,SRCIN1,TBC1D10B,MAP3K4,GAS6,TIAM1 | 0.0458 | SGSM1,MAP3K7CL,RGS9,ARHGAP21,MAP3K6,WRAP53,TBC1D8,JTB,TBC1D10B,NTRK2,TIAM1,TOR1AIP2 |
| **GO:0044093** | Positive regulation of molecular function | 0.0243 | RGS8,RABGAP1L,MID1IP1,MAP3K14,PAK3,AIFM1,MBTPS2,NRK,SRCIN1,TBC1D10B,MAP3K4,GAS6,TIAM1 | 0.0458 | SGSM1,MAP3K7CL,RGS9,PRKCH,ARHGAP21,MAP3K6,WRAP53,TBC1D8,JTB,TBC1D10B,NTRK2,ABCC8,TIAM1,TOR1AIP2 |
| **GO:0050790** | Regulation of catalytic activity | 0.0294 | RGS8,RABGAP1L,DLG3,MID1IP1,MAP3K14,PAK3,AIFM1,APP,NRK,NYX,SRCIN1,TBC1D10B,MAP3K4,GAS6,TIAM1 | 0.0141 | SGSM1,MAP3K7CL,DLG3,SLC9A3R1,RGS9,SYDE1,ARHGAP21,MAP3K6,WRAP53,TBC1D8,MIDN,JTB,TBC1D10B,NTRK2,TIAM1,TOR1AIP2,PPP1R1C,SYDE2,SH3BP5 |

**Supplementary Table 5: GO terms of biological processes enriched by the genes identified from DMR analysis at P7 clustered into groups based on network analysis**

| GO ID | GO Term | FDR | Genes |
|---|---|---|---|
| **Protein Catabolic Process** | | | |
| **GO:0042176** | Regulation of protein catabolic process | 0.00303 | ATG7,COMMD1,CCDC22,LONRF3,RNF144B,RHBDF1,CSNK1D |
| **GO:1903050** | Regulation of proteolysis involved in cellular protein catabolic process | 0.00303 | COMMD1,CCDC22,LONRF3,RNF144B,RHBDF1,CSNK1D |
| **GO:1903362** | Regulation of cellular protein catabolic process | 0.00309 | COMMD1,CCDC22,LONRF3,RNF144B,RHBDF1,CSNK1D |
| **GO:0061136** | Regulation of proteasomal protein catabolic process | 0.00959 | COMMD1,LONRF3,RNF144B,RHBDF1,CSNK1D |
| **GO:0045732** | Positive regulation of protein catabolic process | 0.011 | ATG7,CCDC22,LONRF3,RNF144B,CSNK1D |
| **GO:0032446** | Protein modification by small protein conjugation | 0.0119 | MED12,KLHL15,COMMD1,CCDC22,LONRF3,RNF144B,UBE2F,TRIM71,UBE2A,RNF165 |
| **GO:0031331** | Positive regulation of cellular catabolic process | 0.0128 | ATG7,CCDC22,LONRF3,RNF144B,NPRL2,CSNK1D |
| **GO:0070647** | Protein modification by small protein conjugation or removal | 0.013 | MED12,KLHL15,COMMD1,CCDC22,LONRF3,RNF144B,UBE2F,USP51,TRIM71,UBE2A,RNF165 |
| **GO:0043085** | Positive regulation of catalytic activity | 0.0148 | RGS8,RABGAP1L,MID1IP1,MAP3K14,PAK3,AIFM1,NRK,SRCIN1,TBC1D10B,MAP3K4,GAS6,TIAM1 |
| **GO:0031329** | Regulation of cellular catabolic process | 0.0155 | ATG7,COMMD1,CCDC22,LONRF3,RNF144B,RHBDF1,NPRL2,CSNK1D |
| **GO:0009896** | Positive regulation of catabolic process | 0.0173 | ATG7,CCDC22,LONRF3,RNF144B,NPRL2,CSNK1D |
| **GO:0016567** | Protein ubiquitination | 0.0183 | MED12,KLHL15,COMMD1,CCDC22,LONRF3,RNF144B,TRIM71,UBE2A,RNF165 |

| | | | |
|---|---|---|---|
| **GO:1903052** | Positive regulation of proteolysis involved in cellular protein catabolic process | 0.0184 | CCDC22,LONRF3,RNF144B,CSNK1D |
| **GO:1903364** | Positive regulation of cellular protein catabolic process | 0.0194 | CCDC22,LONRF3,RNF144B,CSNK1D |
| **GO:0032434** | Regulation of proteasomal ubiquitin-dependent protein catabolic process | 0.022 | COMMD1,LONRF3,RNF144B,CSNK1D |
| **GO:0006511** | Ubiquitin-dependent protein catabolic process | 0.0225 | KLHL15,COMMD1,CCDC22,LONRF3,RNF144B,UBA1,CSNK1D,UBE2A,RNF165 |
| **GO:0030163** | Protein catabolic process | 0.0225 | KLHL15,ATG7,COMMD1,CCDC22,LONRF3,RNF144B,UBA1,RHBDF1,CSNK1D,UBE2A,RNF165 |
| **GO:0019941** | Modification-dependent protein catabolic process | 0.0236 | KLHL15,COMMD1,CCDC22,LONRF3,RNF144B,UBA1,CSNK1D,UBE2A,RNF165 |
| **GO:0051603** | Proteolysis involved in cellular protein catabolic process | 0.0236 | KLHL15,COMMD1,CCDC22,LONRF3,RNF144B,UBA1,RHBDF1,CSNK1D,UBE2A,RNF165 |
| **GO:0009894** | Regulation of catabolic process | 0.0243 | ATG7,COMMD1,CCDC22,LONRF3,RNF144B,RHBDF1,NPRL2,CSNK1D |
| **GO:0010498** | Proteasomal protein catabolic process | 0.0243 | KLHL15,COMMD1,LONRF3,RNF144B,RHBDF1,CSNK1D,UBE2A,RNF165 |
| **GO:0043632** | Modification-dependent macromolecule catabolic process | 0.0246 | KLHL15,COMMD1,CCDC22,LONRF3,RNF144B,UBA1,CSNK1D,UBE2A,RNF165 |
| **GO:0044257** | Cellular protein catabolic process | 0.0246 | KLHL15,COMMD1,CCDC22,LONRF3,RNF144B,UBA1,RHBDF1,CSNK1D,UBE2A,RNF165 |
| **GO:0006508** | Proteolysis | 0.0248 | ATP6AP2,AIFM1,KLHL15,COMMD1,CCDC22,LONRF3,RNF144B,GAS6,UBA1,RHBDF1,USP51,UQCRC1,CSNK1D,UBE2A,RNF165 |
| **GO:0030162** | Regulation of proteolysis | 0.0248 | AIFM1,COMMD1,CCDC22,LONRF3,RNF144B,GAS6,RHBDF1,CSNK1D |
| **GO:0045862** | Positive regulation of proteolysis | 0.0305 | AIFM1,CCDC22,LONRF3,RNF144B,CSNK1D |

| | | | |
|---|---|---|---|
| **GO:0043161** | Proteasome-mediated ubiquitin-dependent protein catabolic process | 0.0391 | KLHL15,COMMD1,LONRF3,RNF144B,CSNK1D,UBE2A,RNF165 |
| **GO:1901565** | Organonitrogen compound catabolic process | 0.0411 | SAT1,KLHL15,ATG7,COMMD1,CCDC22,LONRF3,RNF144B,UBA1,RHBDF1,CSNK1D,UBE2A,RNF165 |
| **GO:0030166** | Proteoglycan biosynthetic process | 0.0497 | CHST7,HS6ST2 |
| **Neuronal projection & nervous system development** | | | |
| **GO:0007399** | Nervous system development | 1.65E-04 | FYN,RUNX1,MARK1,DLG3,PCDH19,PAX6,PAK3,BCL11B,APP,BCL11A,THOC2,SLC8A1,FOXP1,LRRN2,TENM4,SRCIN1,GATA2,PTPRU,NAV2,EEF2K,BCL2L11,ACSL4,TIAM1,NNAT,TRIM71,RNF165,SHANK2,PLXNA3,ATP2B3 |
| **GO:0030182** | Neuron differentiation | 0.00296 | RUNX1,PAX6,PAK3,APP,BCL11A,THOC2,LRRN2,TENM4,SRCIN1,GATA2,PTPRU,EEF2K,ACSL4,TIAM1,NNAT,RNF165,PLXNA3 |
| **GO:0048699** | Generation of neurons | 0.00303 | RUNX1,MARK1,PAX6,PAK3,APP,BCL11A,THOC2,LRRN2,TENM4,SRCIN1,GATA2,PTPRU,EEF2K,ACSL4,TIAM1,NNAT,RNF165,PLXNA3 |
| **GO:0022008** | Neurogenesis | 0.00312 | RUNX1,MARK1,PAX6,PAK3,BCL11B,APP,BCL11A,THOC2,LRRN2,TENM4,SRCIN1,GATA2,PTPRU,EEF2K,ACSL4,TIAM1,NNAT,RNF165,PLXNA3 |
| **GO:0016358** | Dendrite development | 0.00482 | PAK3,APP,BCL11A,SRCIN1,EEF2K,ACSL4,TIAM1 |
| **GO:0061001** | Regulation of dendritic spine morphogenesis | 0.00625 | PAK3,SRCIN1,EEF2K,TIAM1 |
| **GO:0016322** | Neuron remodeling | 0.00662 | APP,BCL11A |
| **GO:0060996** | Dendritic spine development | 0.00857 | PAK3,SRCIN1,EEF2K,ACSL4,TIAM1 |
| **GO:0061003** | Positive regulation of dendritic spine morphogenesis | 0.0092 | PAK3,EEF2K,TIAM1 |
| **GO:0048812** | Neuron projection morphogenesis | 0.0111 | PAK3,APP,BCL11A,LRRN2,SRCIN1,EEF2K,TIAM1,RNF165,PLXNA3 |
| **GO:0060997** | Dendritic spine morphogenesis | 0.0111 | PAK3,SRCIN1,EEF2K,TIAM1 |

| | | | |
|---|---|---|---|
| **GO:0048666** | Neuron development | 0.0119 | PAX6,PAK3,APP,BCL11A,THOC2,LRRN2,TENM4,SRCIN1,EEF2K,ACSL4,TIAM1,RNF165,PLXNA3 |
| **GO:0097061** | Dendritic spine organization | 0.014 | PAK3,SRCIN1,EEF2K,TIAM1 |
| **GO:0106027** | Neuron projection organization | 0.014 | PAK3,SRCIN1,EEF2K,TIAM1 |
| **GO:0048667** | Cell morphogenesis involved in neuron differentiation | 0.0154 | PAK3,APP,LRRN2,SRCIN1,EEF2K,TIAM1,RNF165,PLXNA3 |
| **GO:0031175** | Neuron projection development | 0.0169 | PAX6,PAK3,APP,BCL11A,THOC2,LRRN2,SRCIN1,EEF2K,ACSL4,TIAM1,RNF165,PLXNA3 |
| **GO:0060998** | Regulation of dendritic spine development | 0.0194 | PAK3,SRCIN1,EEF2K,TIAM1 |
| **GO:0048814** | Regulation of dendrite morphogenesis | 0.022 | PAK3,SRCIN1,EEF2K,TIAM1 |
| **GO:0050773** | Regulation of dendrite development | 0.0248 | PAK3,BCL11A,SRCIN1,EEF2K,TIAM1 |
| **GO:0050775** | Positive regulation of dendrite morphogenesis | 0.0338 | PAK3,EEF2K,TIAM1 |
| **GO:0010975** | Regulation of neuron projection development | 0.0341 | PAX6,PAK3,BCL11A,THOC2,SRCIN1,EEF2K,TIAM1,PLXNA3 |
| **GO:0045664** | Regulation of neuron differentiation | 0.0343 | PAX6,PAK3,BCL11A,THOC2,SRCIN1,GATA2,EEF2K,TIAM1,PLXNA3 |
| **GO:0060999** | Positive regulation of dendritic spine development | 0.0364 | PAK3,EEF2K,TIAM1 |
| **GO:0048813** | Dendrite morphogenesis | 0.0366 | PAK3,SRCIN1,EEF2K,TIAM1 |
| **GO:0007409** | Axonogenesis | 0.0433 | PAK3,APP,LRRN2,TIAM1,RNF165,PLXNA3 |

| | | | |
|---|---|---|---|
| **GO:0120036** | Plasma membrane bounded cell projection organization | 0.0464 | PAX6,PAK3,APP,BCL11A,THOC2,LRRN2,SRCIN1,EEF2K,ACSL4,TIAM1,ZMYND10,RNF165,PLXNA3 |
| **GO:0008045** | Motor neuron axon guidance | 0.0497 | RNF165,PLXNA3 |
| **Secretion & signal transduction** | | | |
| **GO:0046903** | Secretion | 0.00241 | NCOR2,RUNX1,PDZD11,APLN,PAX6,SYTL1,TNFAIP2,CACNA1F,SRCIN1,GATA2,GAS6,RHBDF1,TIAM1,NNAT,TSC22D3 |
| **GO:0032940** | Secretion by cell | 0.00482 | RUNX1,PDZD11,APLN,PAX6,SYTL1,TNFAIP2,CACNA1F,SRCIN1,GATA2,GAS6,RHBDF1,TIAM1,NNAT |
| **GO:0035235** | Ionotropic glutamate receptor signaling pathway | 0.00501 | FYN,APP,TIAM1 |
| **GO:0023051** | Regulation of signaling | 0.00509 | FYN,RUNX1,RGS8,MAP3K14,APLN,PAX6,PAK3,APP,NYX,IGFBP6,CTNNBIP1,GAS6,DLK1,RHBDF1,TIAM1,NPRL2,NNAT,FAM109B,RNF165,SHANK2,SMOK2A,BMP7,IQSEC2 |
| **GO:0007010** | Cytoskeleton organization | 0.00591 | CIT,MARK1,MID1IP1,SH3KBP1,PAK3,MAP7D1,ZMYM4,SYNPO2,ZMYND10,MAST3,MAP7D3,EML2,TMOD3,IQSEC2 |
| **GO:0010646** | Regulation of cell communication | 0.00857 | FYN,RUNX1,RGS8,MAP3K14,APLN,PAX6,PAK3,APP,NYX,IGFBP6,CTNNBIP1,GAS6,DLK1,RHBDF1,TIAM1,NPRL2,NNAT,RNF165,SHANK2,SMOK2A,BMP7,IQSEC2 |
| **GO:1903530** | Regulation of secretion by cell | 0.0119 | RUNX1,APLN,PAX6,SYTL1,SRCIN1,GATA2,GAS6,RHBDF1,TIAM1,NNAT |
| **GO:0045807** | Positive regulation of endocytosis | 0.0149 | APLN,GATA2,EEF2K,GAS6 |
| **GO:0051046** | Regulation of secretion | 0.0157 | RUNX1,APLN,PAX6,SYTL1,SRCIN1,GATA2,GAS6,RHBDF1,TIAM1,NNAT |
| **GO:0023061** | Signal release | 0.0172 | RUNX1,PDZD11,APLN,PAX6,SYTL1,CACNA1F,TIAM1,NNAT |
| **GO:0002791** | Regulation of peptide secretion | 0.0173 | APLN,PAX6,SRCIN1,GAS6,RHBDF1,TIAM1,NNAT |
| **GO:0007267** | Cell-cell signaling | 0.0173 | RUNX1,DLG3,PDZD11,PCDH19,APLN,PAX6,FZD6,SYTL1,CACNA1F,CTNNBIP1,TIAM1,NNAT,CSNK1D,SHANK2,IQSEC2 |

| | | | |
|---|---|---|---|
| **GO:0009966** | Regulation of signal transduction | 0.0259 | FYN,RGS8,MAP3K14,APLN,PAK3,APP,NYX,IGFBP6,CTNNBIP1,GAS6,DLK1,RHBDF1,TIAM1,NPRL2,RNF165,SMOK2A,BMP7 |
| **GO:0002790** | Peptide secretion | 0.0293 | APLN,PAX6,SRCIN1,GAS6,RHBDF1,TIAM1,NNAT |
| **GO:0051049** | Regulation of transport | 0.0304 | RUNX1,RABGAP1L,APLN,PAX6,SLC8A1,SYTL1,COMMD1,SRCIN1,GATA2,EEF2K,TBC1D10B,GAS6,RHBDF1,TIAM1,NNAT |
| **GO:0023057** | Negative regulation of signaling | 0.0336 | FYN,APLN,NYX,CTNNBIP1,GAS6,DLK1,NPRL2,SHANK2,SMOK2A,IQSEC2 |
| **GO:0010648** | Negative regulation of cell communication | 0.0338 | FYN,APLN,NYX,CTNNBIP1,GAS6,DLK1,NPRL2,SHANK2,SMOK2A,IQSEC2 |
| **GO:0060627** | Regulation of vesicle-mediated transport | 0.0372 | RABGAP1L,APLN,SYTL1,GATA2,EEF2K,TBC1D10B,GAS6 |
| **GO:0046887** | Positive regulation of hormone secretion | 0.0461 | RUNX1,APLN,PAX6,NNAT |
| **GO:0010647** | Positive regulation of cell communication | 0.0463 | FYN,RUNX1,MAP3K14,APLN,PAX6,GAS6,TIAM1,NNAT,RNF165,SHANK2,BMP7,IQSEC2 |
| **GO:0023056** | Positive regulation of signaling | 0.0472 | FYN,RUNX1,MAP3K14,APLN,PAX6,GAS6,TIAM1,NNAT,RNF165,SHANK2,BMP7,IQSEC2 |
| **GO:0033036** | Macromolecule localization | 0.0472 | RUNX1,RABGAP1L,DLG3,PDZD11,THOC2,COMMD1,TNFAIP2,SRCIN1,IGF2R,TBC1D10B,GAS6,ACSL4,RHBDF1,TIAM1,NNAT,UBL4A,BCAP31 |
| **GO:0007215** | Glutamate receptor signaling pathway | 0.0479 | FYN,APP,TIAM1 |
| **Cellular morphogenesis & development** | | | |
| **GO:0051128** | Regulation of cellular component organization | 3.22E-04 | CIT,RABGAP1L,MID1IP1,APLN,SH3KBP1,PAX6,PAK3,BCL11A,THOC2,ATG7,FOXP1,SRCIN1,GATA2,ZMYM4,SYNPO2,EEF2K,TBC1D10B,GAS6,ANK1,TIAM1,NPRL2,EML2,CSNK1D,PLXNA3 |
| **GO:0032989** | Cellular component morphogenesis | 4.63E-04 | NOTCH4,SH3KBP1,PAK3,APP,BCL11A,LRRN2,TENM4,SRCIN1,ZMYM4,EEF2K,TIAM1,TMOD3,CSNK1D,RNF165,PLXNA3 |
| **GO:0000902** | Cell morphogenesis | 6.07E-04 | NOTCH4,SH3KBP1,PAK3,APP,BCL11A,LRRN2,TENM4,SRCIN1,ZMYM4,EEF2K,TIAM1,CSNK1D,RNF165,PLXNA3 |

| | | | |
|---|---|---|---|
| **GO:0040011** | Locomotion | 0.00309 | FYN,MARK1,USP9X,SH3KBP1,PAX6,PAK3,APP,SLC8A1,SRCIN1,GATA2,GAS6,RHBDF1,TIAM1,SHROOM2,RNF165,PLXNA3 |
| **GO:0048468** | Cell development | 0.00365 | NOTCH4,CDKN1A,PAX6,PAK3,APP,BCL11A,THOC2,FOXP1,LRRN2,TENM4,SRCIN1,SMARCA2,GATA2,EEF2K,ACSL4,TIAM1,TMOD3,RNF165,BMP7,PLXNA3 |
| **GO:0022604** | Regulation of cell morphogenesis | 0.00432 | SH3KBP1,PAK3,BCL11A,SRCIN1,ZMYM4,EEF2K,TIAM1,CSNK1D,PLXNA3 |
| **GO:0000904** | Cell morphogenesis involved in differentiation | 0.00959 | NOTCH4,PAK3,APP,LRRN2,SRCIN1,EEF2K,TIAM1,RNF165,PLXNA3 |
| **GO:0006928** | Movement of cell or subcellular component | 0.0106 | FYN,MARK1,USP9X,SH3KBP1,PAX6,PAK3,APP,SLC8A1,SRCIN1,GATA2,GAS6,RHBDF1,TIAM1,SHROOM2,RNF165,PLXNA3 |
| **GO:0032990** | Cell part morphogenesis | 0.0112 | PAK3,APP,BCL11A,LRRN2,SRCIN1,EEF2K,TIAM1,RNF165,PLXNA3 |
| **GO:0048858** | Cell projection morphogenesis | 0.0112 | PAK3,APP,BCL11A,LRRN2,SRCIN1,EEF2K,TIAM1,RNF165,PLXNA3 |
| **GO:0048870** | Cell motility | 0.0112 | FYN,MARK1,USP9X,SH3KBP1,PAX6,PAK3,SLC8A1,SRCIN1,GATA2,GAS6,RHBDF1,TIAM1,SHROOM2 |
| **GO:0051674** | Localization of cell | 0.0112 | FYN,MARK1,USP9X,SH3KBP1,PAX6,PAK3,SLC8A1,SRCIN1,GATA2,GAS6,RHBDF1,TIAM1,SHROOM2 |
| **GO:0120039** | Plasma membrane bounded cell projection morphogenesis | 0.0112 | PAK3,APP,BCL11A,LRRN2,SRCIN1,EEF2K,TIAM1,RNF165,PLXNA3 |
| **GO:0048588** | Developmental cell growth | 0.0338 | APP,BCL11A,FOXP1,TIAM1,PLXNA3 |
| **GO:0033043** | Regulation of organelle organization | 0.0372 | CIT,RABGAP1L,MID1IP1,PAK3,ATG7,SYNPO2,TBC1D10B,ANK1,NPRL2,EML2 |
| **GO:0045595** | Regulation of cell differentiation | 0.0389 | RUNX1,CITED1,PAX6,PAK3,BCL11A,THOC2,FOXP1,SRCIN1,GATA2,EEF2K,GAS6,TIAM1,PLXNA3 |
| **GO:0001763** | Morphogenesis of a branching structure | 0.0391 | CITED1,BCL11A,CTNNBIP1 |

| | | | |
|---|---|---|---|
| **GO:0010763** | Positive regulation of fibroblast migration | 0.0433 | PAK3,SLC8A1 |
| **GO:1902903** | Regulation of supramolecular fiber organization | 0.0463 | CIT,MID1IP1,PAK3,SYNPO2,EML2 |
| **Gene expression regulation** | | | |
| **GO:0006366** | Transcription from RNA polymerase II promoter | 0.00241 | NCOR2,RUNX1,POU3F4,CDX4,CITED1,MED14,MED12,PAX6,BCL11A,MBTPS2,FOXP1,TFE3,GATA2,NFATC1,CAMTA1,MEOX1,TRERF1,RNF165,HCFC1,BMP7 |
| **GO:0006357** | Regulation of transcription from RNA polymerase II promoter | 0.00482 | NCOR2,RUNX1,CDX4,CITED1,MED14,MED12,BCL11A,MBTPS2,FOXP1,TFE3,GATA2,NFATC1,CAMTA1,MEOX1,TRERF1,RNF165,HCFC1,BMP7 |
| **GO:0006355** | Regulation of transcription, DNA-templated | 0.00857 | NCOR2,RUNX1,CDX4,CITED1,NONO,MED14,MED12,PAX6,BCL11A,MBTPS2,FOXP1,TFE3,MED16,GATA2,ATXN1,NFATC1,GAS6,CAMTA1,MEOX1,TRERF1,RNF165,HCFC1,BMP7,ZFP275 |
| **GO:1903506** | Regulation of nucleic acid-templated transcription | 0.0091 | NCOR2,RUNX1,CDX4,CITED1,NONO,MED14,MED12,PAX6,BCL11A,MBTPS2,FOXP1,TFE3,MED16,GATA2,ATXN1,NFATC1,GAS6,CAMTA1,MEOX1,TRERF1,RNF165,HCFC1,BMP7,ZFP275 |
| **GO:2001141** | Regulation of RNA biosynthetic process | 0.0092 | NCOR2,RUNX1,CDX4,CITED1,NONO,MED14,MED12,PAX6,BCL11A,MBTPS2,FOXP1,TFE3,MED16,GATA2,ATXN1,NFATC1,GAS6,CAMTA1,MEOX1,TRERF1,RNF165,HCFC1,BMP7,ZFP275 |
| **GO:0010628** | Positive regulation of gene expression | 0.017 | RUNX1,CITED1,MED14,PAX6,BCL11A,MBTPS2,TFE3,MED16,GATA2,RBM20,NFATC1,GAS6,CAMTA1,MEOX1 |
| **GO:0051254** | Positive regulation of RNA metabolic process | 0.0209 | RUNX1,CITED1,MED14,PAX6,MBTPS2,TFE3,MED16,GATA2,RBM20,NFATC1,CAMTA1,MEOX1 |
| **GO:0044093** | Positive regulation of molecular function | 0.0243 | RGS8,RABGAP1L,MID1IP1,MAP3K14,PAK3,AIFM1,MBTPS2,NRK,SRCIN1,TBC1D10B,MAP3K4,GAS6,TIAM1 |
| **GO:2000113** | Negative regulation of cellular macromolecule biosynthetic process | 0.0294 | NCOR2,CITED1,NONO,BCL11A,ATG7,FOXP1,ATXN1,GAS6,SNIP1,TRIM71,HCFC1 |
| **GO:0045893** | Positive regulation of transcription, DNA-templated | 0.0305 | RUNX1,CITED1,MED14,PAX6,MBTPS2,TFE3,MED16,GATA2,NFATC1,CAMTA1,MEOX1 |
| **GO:1902680** | Positive regulation of RNA biosynthetic process | 0.0305 | RUNX1,CITED1,MED14,PAX6,MBTPS2,TFE3,MED16,GATA2,NFATC1,CAMTA1,MEOX1 |
| **GO:1903508** | Positive regulation of nucleic acid-templated transcription | 0.0305 | RUNX1,CITED1,MED14,PAX6,MBTPS2,TFE3,MED16,GATA2,NFATC1,CAMTA1,MEOX1 |

| | | | |
|---|---|---|---|
| **GO:0010558** | Negative regulation of macromolecule biosynthetic process | 0.0358 | NCOR2,CITED1,NONO,BCL11A,ATG7,FOXP1,ATXN1,GAS6,SNIP1,TRIM71,HCFC1 |
| **Cellular stimulus response** | | | |
| **GO:0071310** | Cellular response to organic substance | 0.00296 | FYN,MED14,PAX6,AIFM1,KLHL15,APP,BCL11A,ATG7,MBTPS2,SLC8A1,FOXP1,NYX,TWIST1,IL20RA,EEF2K,GAS6,TIAM1,SNIP1,TRIM71,CSNK1D,RNF165,HCFC1,BMP7,FGF16 |
| **GO:0014070** | Response to organic cyclic compound | 0.00676 | CDKN1A,NCOR2,CITED1,MED14,AIFM1,APP,ATG7,SLC8A1,FOXP1,ABCC4,PTPRU,EEF2K,CAR9,TIAM1,PAQR7,SNIP1,PAQR6,HCFC1,BMP7 |
| **GO:0071363** | Cellular response to growth factor stimulus | 0.0129 | FYN,PAX6,APP,TWIST1,EEF2K,TRIM71,CSNK1D,RNF165,BMP7,FGF16 |
| **GO:1901698** | Response to nitrogen compound | 0.0129 | CDKN1A,FYN,NCOR2,RGS8,CITED1,PAX6,AIFM1,APP,BCL11A,ATG7,SLC8A1,ABCC4,EEF2K,TIAM1,SNIP1,UQCRC1,BMP7 |
| **GO:0009719** | Response to endogenous stimulus | 0.013 | CDKN1A,FYN,CITED1,MED14,PAX6,AIFM1,APP,BCL11A,ATG7,SLC8A1,PTPRU,EEF2K,PAQR7,PAQR6,TRIM71,CSNK1D,RNF165,BMP7,FGF16 |
| **GO:0033554** | Cellular response to stress | 0.0172 | CDKN1A,MAP3K14,PAK3,AIFM1,KLHL15,ATG7,MBTPS2,SLC8A1,NRK,FOXP1,FTO,EEF2K,MAP3K4,UBA1,TIAM1,NPRL2,UBE2A,SMOK2A |
| **GO:0070848** | Response to growth factor | 0.0183 | FYN,PAX6,APP,TWIST1,EEF2K,TRIM71,CSNK1D,RNF165,BMP7,FGF16 |
| **GO:1901699** | Cellular response to nitrogen compound | 0.0203 | FYN,RGS8,PAX6,AIFM1,APP,BCL11A,ATG7,SLC8A1,EEF2K,SNIP1 |
| **GO:0071495** | Cellular response to endogenous stimulus | 0.0248 | FYN,MED14,PAX6,AIFM1,APP,BCL11A,ATG7,SLC8A1,EEF2K,TRIM71,CSNK1D,RNF165,BMP7,FGF16 |
| **GO:0010243** | Response to organonitrogen compound | 0.0264 | CDKN1A,FYN,NCOR2,CITED1,PAX6,AIFM1,APP,BCL11A,ATG7,SLC8A1,ABCC4,EEF2K,TIAM1,UQCRC1,BMP7 |
| **GO:0071407** | Cellular response to organic cyclic compound | 0.0295 | MED14,AIFM1,APP,ATG7,SLC8A1,FOXP1,EEF2K,SNIP1,HCFC1 |
| **GO:0033993** | Response to lipid | 0.0305 | CDKN1A,NCOR2,CITED1,MED14,PAX6,AIFM1,FOXP1,GATA2,PTPRU,IGF2R,CAR9,PAQR7,RELT,PAQR6,BMP7 |
| **GO:1902065** | Response to L-glutamate | 0.0497 | AIFM1,BCL11A |

| | | | |
|---|---|---|---|
| **Protein phosphorylation regulation** | | | |
| **GO:0016310** | Phosphorylation | 0.0178 | FYN,MARK1,MAP3K14,APLN,PAK3,APP,NRK,NYX,SRCIN1,EEF2K,MAP3K4,GAS6,TIAM1,MAST3,UQCRC1,CSNK1D,SMOK2A,BMP7,PGK1 |
| **GO:0061097** | Regulation of protein tyrosine kinase activity | 0.0184 | APP,SRCIN1,GAS6 |
| **GO:0045859** | Regulation of protein kinase activity | 0.0244 | MAP3K14,PAK3,APP,NRK,NYX,SRCIN1,MAP3K4,GAS6,TIAM1 |
| **GO:0043549** | Regulation of kinase activity | 0.0288 | MAP3K14,PAK3,APP,NRK,NYX,SRCIN1,MAP3K4,GAS6,TIAM1 |
| **GO:0050790** | Regulation of catalytic activity | 0.0294 | RGS8,RABGAP1L,DLG3,MID1IP1,MAP3K14,PAK3,AIFM1,APP,NRK,NYX,SRCIN1,TBC1D10B,MAP3K4,GAS6,TIAM1 |
| **GO:0045860** | Positive regulation of protein kinase activity | 0.0304 | MAP3K14,PAK3,NRK,SRCIN1,MAP3K4,GAS6,TIAM1 |
| **GO:0033674** | Positive regulation of kinase activity | 0.0343 | MAP3K14,PAK3,NRK,SRCIN1,MAP3K4,GAS6,TIAM1 |
| **GO:0006468** | Protein phosphorylation | 0.0381 | FYN,MARK1,MAP3K14,PAK3,APP,NRK,NYX,SRCIN1,EEF2K,MAP3K4,GAS6,TIAM1,MAST3,CSNK1D,SMOK2A,BMP7 |
| **GO:0061098** | Positive regulation of protein tyrosine kinase activity | 0.0433 | SRCIN1,GAS6 |
| **GO:0051338** | Regulation of transferase activity | 0.0445 | MAP3K14,PAK3,APP,NRK,NYX,SRCIN1,MAP3K4,GAS6,TIAM1 |
| **GO:0032147** | Activation of protein kinase activity | 0.0458 | MAP3K14,PAK3,NRK,MAP3K4,GAS6 |
| **GO:0018212** | Peptidyl-tyrosine modification | 0.0463 | FYN,TPST2,APP,SRCIN1,GAS6 |
| **GO:0051347** | Positive regulation of transferase activity | 0.047 | MAP3K14,PAK3,NRK,SRCIN1,MAP3K4,GAS6,TIAM1 |
| **Metabolic regulation** | | | |

| | | | |
|---|---|---|---|
| **GO:0009893** | Positive regulation of metabolic process | 9.52E-05 | RUNX1,CITED1,MID1IP1,MAP3K14,MED14,APLN,PAX6,PAK3,AIFM1,APP,BCL11A,ATG7,MBTPS2,NRK,COMMD1,TFE3,CCDC22,MED16,SRCIN1,GATA2,LONRF3,RBM20,RNF144B,NFATC1,MAP3K4,GAS6,CAMTA1,MEOX1,TIAM1,NPRL2,CSNK1D,BMP7 |
| **GO:0010604** | Positive regulation of macromolecule metabolic process | 9.52E-05 | RUNX1,CITED1,MAP3K14,MED14,APLN,PAX6,PAK3,AIFM1,APP,BCL11A,ATG7,MBTPS2,NRK,COMMD1,TFE3,CCDC22,MED16,SRCIN1,GATA2,LONRF3,RBM20,RNF144B,NFATC1,MAP3K4,GAS6,CAMTA1,MEOX1,TIAM1,CSNK1D,BMP7 |
| **GO:0031325** | Positive regulation of cellular metabolic process | 1.65E-04 | RUNX1,CITED1,MID1IP1,MAP3K14,MED14,APLN,PAX6,PAK3,AIFM1,ATG7,MBTPS2,NRK,COMMD1,TFE3,CCDC22,MED16,SRCIN1,GATA2,LONRF3,RBM20,RNF144B,NFATC1,MAP3K4,GAS6,CAMTA1,MEOX1,TIAM1,NPRL2,CSNK1D,BMP7 |
| **GO:0051173** | Positive regulation of nitrogen compound metabolic process | 2.84E-04 | RUNX1,CITED1,MAP3K14,MED14,PAX6,PAK3,AIFM1,APP,ATG7,MBTPS2,NRK,COMMD1,TFE3,CCDC22,MED16,SRCIN1,GATA2,LONRF3,RBM20,RNF144B,NFATC1,MAP3K4,GAS6,CAMTA1,MEOX1,TIAM1,CSNK1D,BMP7 |
| **GO:0051247** | Positive regulation of protein metabolic process | 0.00784 | MAP3K14,PAK3,AIFM1,APP,ATG7,NRK,COMMD1,CCDC22,SRCIN1,LONRF3,RNF144B,MAP3K4,GAS6,TIAM1,CSNK1D,BMP7 |
| **GO:0051246** | Regulation of protein metabolic process | 0.0111 | MAP3K14,ATP6AP2,PAK3,AIFM1,APP,ATG7,NRK,COMMD1,NYX,CCDC22,SRCIN1,LONRF3,RNF144B,MAP3K4,GAS6,RHBDF1,TIAM1,SNIP1,TRIM71,CSNK1D,BMP7 |
| **GO:0032268** | Regulation of cellular protein metabolic process | 0.0182 | MAP3K14,PAK3,AIFM1,APP,NRK,COMMD1,NYX,CCDC22,SRCIN1,LONRF3,RNF144B,MAP3K4,GAS6,RHBDF1,TIAM1,SNIP1,TRIM71,CSNK1D,BMP7 |
| **GO:0032270** | Positive regulation of cellular protein metabolic process | 0.0215 | MAP3K14,PAK3,AIFM1,NRK,COMMD1,CCDC22,SRCIN1,LONRF3,RNF144B,MAP3K4,GAS6,TIAM1,CSNK1D,BMP7 |
| **GO:0008215** | Spermine metabolic process | 0.0294 | SAT1,SMS |
| **GO:0055070** | Copper ion homeostasis | 0.0294 | APP,COMMD1 |
| **GO:0045935** | Positive regulation of nucleobase-containing compound metabolic process | 0.0302 | RUNX1,CITED1,MED14,PAX6,PAK3,MBTPS2,TFE3,MED16,GATA2,RBM20,NFATC1,CAMTA1,MEOX1 |
| **Organism development** | | | |
| **GO:0009653** | Anatomical structure morphogenesis | 1.65E-04 | NOTCH4,RUNX1,POU3F4,CDX4,CITED1,SH3KBP1,PAX6,PAK3,APP,BCL11A,FOXP1,LRRN2,TWIST1,TENM4,SRCIN1,ZMYM4,CTNNBIP1,EEF2K,TIAM1,TMOD3,CSNK1D,RNF165,PLXNA3 |
| **GO:0016477** | Cell migration | 0.00676 | FYN,MARK1,USP9X,SH3KBP1,PAX6,PAK3,SLC8A1,SRCIN1,GATA2,GAS6,RHBDF1,TIAM1,SHROOM2 |
| **GO:0022603** | Regulation of anatomical structure morphogenesis | 0.0112 | RUNX1,CITED1,SH3KBP1,PAK3,BCL11A,SRCIN1,ZMYM4,EEF2K,TIAM1,CSNK1D,PLXNA3 |

| | | | |
|---|---|---|---|
| **GO:0048513** | Animal organ development | 0.0149 | CDKN1A,NCOR2,RUNX1,POU3F4,CDX4,CITED1,PAX6,SLC8A1,FOXP1,TWIST1,GATA2,PTPRU,RBM20,IGF2R,CTNNBIP1,BCL2L11,GAS6,SMARCC1,NNAT,NHS,ZFPM1,SHANK2,ATP2B3 |
| **GO:0050793** | Regulation of developmental process | 0.0206 | RUNX1,CITED1,SH3KBP1,PAX6,PAK3,APP,BCL11A,THOC2,FOXP1,SRCIN1,GATA2,ZMYM4,EEF2K,GAS6,TIAM1,CSNK1D,PLXNA3 |
| **GO:0021700** | Developmental maturation | 0.0305 | CDKN1A,ZDHHC15,APP,BCL11A |
| **GO:0060429** | Epithelium development | 0.0488 | NOTCH4,CDKN1A,CITED1,PAX6,CTNNBIP1,MEOX1,TRIM71 |
| **GO:0002066** | Columnar/cuboidal epithelial cell development | 0.0497 | CDKN1A,PAX6 |
| **Apoptosis** | | | |
| **GO:0008219** | Cell death | 8.76E-04 | CDKN1A,FYN,POU3F4,MAP3K14,SH3KBP1,PAX6,PAK3,AIFM1,MAGED1,ATG7,NRK,FOXP1,TWIST1,IGF2R,EEF2K,BCL2L11,MAP3K4,GAS6,TIAM1,RELT,PLAGL1,BMP7,TSC22D3 |
| **GO:0010941** | Regulation of cell death | 9.30E-04 | CDKN1A,FYN,POU3F4,MAP3K14,SH3KBP1,PAX6,PAK3,AIFM1,MAGED1,ATG7,NRK,FOXP1,TWIST1,IGF2R,EEF2K,BCL2L11,MAP3K4,GAS6,RELT,BMP7,TSC22D3 |
| **GO:0006915** | Apoptotic process | 0.0011 | CDKN1A,FYN,POU3F4,MAP3K14,SH3KBP1,PAK3,AIFM1,MAGED1,ATG7,NRK,TWIST1,IGF2R,EEF2K,BCL2L11,MAP3K4,GAS6,TIAM1,RELT,PLAGL1,BMP7,TSC22D3 |
| **GO:0012501** | Programmed cell death | 0.00128 | CDKN1A,FYN,POU3F4,MAP3K14,SH3KBP1,PAK3,AIFM1,MAGED1,ATG7,NRK,TWIST1,IGF2R,EEF2K,BCL2L11,MAP3K4,GAS6,TIAM1,RELT,PLAGL1,BMP7,TSC22D3 |
| **GO:0042981** | Regulation of apoptotic process | 0.0013 | CDKN1A,FYN,POU3F4,MAP3K14,SH3KBP1,PAK3,AIFM1,MAGED1,ATG7,NRK,TWIST1,IGF2R,EEF2K,BCL2L11,MAP3K4,GAS6,RELT,BMP7,TSC22D3 |
| **GO:0043067** | Regulation of programmed cell death | 0.00141 | CDKN1A,FYN,POU3F4,MAP3K14,SH3KBP1,PAK3,AIFM1,MAGED1,ATG7,NRK,TWIST1,IGF2R,EEF2K,BCL2L11,MAP3K4,GAS6,RELT,BMP7,TSC22D3 |
| **GO:0010942** | Positive regulation of cell death | 0.0294 | CDKN1A,PAX6,PAK3,AIFM1,MAGED1,ATG7,FOXP1,IGF2R,BCL2L11 |

**Supplementary Table 6: GO terms of biological processes enriched by the genes identified from DMR analysis at P90 clustered into groups based on network analysis**

| GO ID | GO term | FDR | Genes |
|---|---|---|---|
| **Protein Localization and secretion** | | | |
| **GO:0032879** | Regulation of localization | 1.19E-04 | SRGN,SGSM1,P2RX7,CD47,MYLK,KIT,SLC9A3R1,ANO6,NCS1,COMMD1,EDNRB,TNFRSF21,KCNAB2,PTPRM,EPHB2,TBC1D8,HSPD1,MIDN,UVRAG,TBC1D10B,NEDD4L,NTRK2,ABCC8,TMEM231,TIAM1,NKAIN4,PARD3,MLC1,JSRP1,SLC25A5,GLUL |
| **GO:0008104** | Protein localization | 0.0034 | SRGN,SGSM1,P2RX7,DLG3,STX8,COMMD1,TNFRSF21,KCNAB2,EPHB2,TBC1D8,HSPD1,IGF2R,GOLGA7B,MIDN,TBC1D10B,ABCC8,TMEM231,TIAM1,ZBTB16,PARD3,MLC1,FGF13,GLUL |
| **GO:0033036** | Macromolecule localization | 0.00402 | SRGN,SGSM1,P2RX7,DLG3,SLC9A3R1,STX8,COMMD1,TNFRSF21,KCNAB2,EPHB2,TBC1D8,HSPD1,IGF2R,GOLGA7B,MIDN,TBC1D10B,PRPF18,ABCC8,TMEM231,TIAM1,ZBTB16,PARD3,MLC1,FGF13,GLUL |
| **GO:0032880** | Regulation of protein localization | 0.00643 | SRGN,P2RX7,COMMD1,TNFRSF21,KCNAB2,EPHB2,HSPD1,MIDN,ABCC8,TMEM231,TIAM1,GLUL |
| **GO:0051641** | Cellular localization | 0.011 | SRGN,SGSM1,DLG3,STX8,SLC25A29,ARHGAP21,COMMD1,KCNAB2,EPHB2,TBC1D8,SLC25A25,HSPD1,GOLGA7B,ATXN1,TBC1D10B,CNP,PRPF18,NTRK2,TIAM1,TRAPPC10,ZBTB16,PARD3,MLC1,JSRP1,SLC25A5,FGF13 |
| **GO:0050708** | Regulation of protein secretion | 0.0116 | SRGN,P2RX7,TNFRSF21,HSPD1,MIDN,ABCC8,TIAM1,GLUL |
| **GO:0002791** | Regulation of peptide secretion | 0.0199 | SRGN,P2RX7,TNFRSF21,HSPD1,MIDN,ABCC8,TIAM1,GLUL |
| **GO:0015833** | Peptide transport | 0.0199 | SRGN,SGSM1,P2RX7,SLC9A3R1,STX8,TNFRSF21,TBC1D8,HSPD1,IGF2R,GOLGA7B,MIDN,TBC1D10B,ABCC8,TIAM1,PARD3,MLC1,GLUL |

| | | | |
|---|---|---|---|
| **GO:0009306** | Protein secretion | 0.0238 | SRGN,P2RX7,TNFRSF21,HSPD1,MIDN,ABCC8,TIAM1,GLUL |
| **GO:0050707** | Regulation of cytokine secretion | 0.0238 | SRGN,P2RX7,TNFRSF21,HSPD1 |
| **GO:0051046** | Regulation of secretion | 0.0261 | SRGN,P2RX7,NCS1,EDNRB,TNFRSF21,HSPD1,MIDN,NTRK2,ABCC8,TIAM1,GLUL |
| **GO:0015031** | Protein transport | 0.0286 | SRGN,SGSM1,P2RX7,STX8,TNFRSF21,TBC1D8,HSPD1,IGF2R,GOLGA7B,MIDN,TBC1D10B,ABCC8,TIAM1,PARD3,MLC1,GLUL |
| **GO:0070727** | Cellular macromolecule localization | 0.0288 | SRGN,SGSM1,DLG3,STX8,COMMD1,KCNAB2,EPHB2,TBC1D8,HSPD1,GOLGA7B,TBC1D10B,PRPF18,TIAM1,ZBTB16,PARD3,FGF13 |
| **GO:0045184** | Establishment of protein localization | 0.0313 | SRGN,SGSM1,P2RX7,STX8,TNFRSF21,TBC1D8,HSPD1,IGF2R,GOLGA7B,MIDN,TBC1D10B,ABCC8,TIAM1,PARD3,MLC1,GLUL |
| **GO:0002790** | Peptide secretion | 0.0314 | SRGN,P2RX7,TNFRSF21,HSPD1,MIDN,ABCC8,TIAM1,GLUL |
| **GO:0050663** | Cytokine secretion | 0.0314 | SRGN,P2RX7,TNFRSF21,HSPD1 |
| **GO:0050709** | Negative regulation of protein secretion | 0.0314 | SRGN,TNFRSF21,MIDN,ABCC8 |

| | | | |
|---|---|---|---|
| **GO:0051085** | Chaperone mediated protein folding requiring cofactor | 0.0314 | HSPD1,HSPE1 |
| **GO:0002792** | Negative regulation of peptide secretion | 0.0365 | SRGN,TNFRSF21,MIDN,ABCC8 |
| **GO:1903530** | Regulation of secretion by cell | 0.0365 | SRGN,P2RX7,NCS1,TNFRSF21,HSPD1,MIDN,NTRK2,ABCC8,TIAM1,GLUL |
| **GO:0034613** | Cellular protein localization | 0.0413 | SRGN,SGSM1,DLG3,STX8,COMMD1,KCNAB2,EPHB2,TBC1D8,HSPD1,GOLGA7B,TBC1D10B,TIAM1,ZBTB16,PARD3,FGF13 |
| **Ion and other substances cellular transport** | | | |
| **GO:0051049** | Regulation of transport | 6.57E-04 | SRGN,SGSM1,P2RX7,CD47,ANO6,NCS1,COMMD1,EDNRB,TNFRSF21,KCNAB2,TBC1D8,HSPD1,MIDN,UVRAG,TBC1D10B,NEDD4L,NTRK2,ABCC8,TIAM1,NKAIN4,MLC1,JSRP1,SLC25A5,GLUL |
| **GO:0071705** | Nitrogen compound transport | 0.0199 | SRGN,SGSM1,P2RX7,SLC9A3R1,STX8,SLC25A29,SLC38A1,TNFRSF21,TBC1D8,HSPD1,IGF2R,GOLGA7B,MIDN,TBC1D10B,ABCC8,TIAM1,SLC6A5,PARD3,MLC1,GLUL |
| **GO:1901379** | Regulation of potassium ion transmembrane transport | 0.0223 | ANO6,KCNAB2,NEDD4L,ABCC8 |
| **GO:0042886** | Amide transport | 0.0232 | SRGN,SGSM1,P2RX7,SLC9A3R1,STX8,TNFRSF21,TBC1D8,HSPD1,IGF2R,GOLGA7B,MIDN,TBC1D10B,ABCC8,TIAM1,PARD3,MLC1,GLUL |
| **GO:0015672** | Monovalent inorganic cation transport | 0.0261 | P2RX7,SLC4A4,ANO6,COMMD1,KCNAB2,NEDD4L,ABCC8,NKAIN4,KCNK5 |

| | | | |
|---|---|---|---|
| **GO:0010959** | Regulation of metal ion transport | 0.0314 | P2RX7,ANO6,COMMD1,KCNAB2,NEDD4L,ABCC8,NKAIN4,JSRP1 |
| **GO:0071702** | Organic substance transport | 0.0428 | SRGN,SGSM1,P2RX7,SLC9A3R1,STX8,SLC25A29,SLC38A1,TNFRSF21,TBC1D8,HSPD1,IGF2R,GOLGA7B,MIDN,TBC1D10B,SLC2A5,ABCC8,TIAM1,SLC6A5,PARD3,MLC1,GLUL |
| **GO:0071804** | Cellular potassium ion transport | 0.0471 | ANO6,KCNAB2,NEDD4L,ABCC8,KCNK5 |
| **GO:0071805** | Potassium ion transmembrane transport | 0.0471 | ANO6,KCNAB2,NEDD4L,ABCC8,KCNK5 |
| **GO:0030001** | Metal ion transport | 0.0487 | P2RX7,SLC4A4,ANO6,SLC24A2,COMMD1,KCNAB2,NEDD4L,ABCC8,NKAIN4,JSRP1,KCNK5 |
| **Nervous System development and neuronal projection regulation** | | | |
| **GO:0007399** | Nervous system development | 7.97E-04 | FYN,FRY,KIT,PLP1,DLG3,RGS9,PRKCH,PTPRO,EPHB1,NCS1,DPYSL5,FOXP1,EDNRB,TENM4,TNFRSF21,PTPRM,TENM3,TUBB3,CNP,NTRK2,TACC2,SOX6,ABCC8,TIAM1,ZBTB16,PARD3,KDM6B,PCDH9,FGF13,IMPACT,DMD |
| **GO:0022008** | Neurogenesis | 0.00592 | FRY,KIT,PLP1,PRKCH,PTPRO,EPHB1,NCS1,DPYSL5,TENM4,TNFRSF21,PTPRM,TENM3,TUBB3,NTRK2,SOX6,ABCC8,TIAM1,PARD3,FGF13,IMPACT,DMD |
| **GO:0030900** | Forebrain development | 0.0143 | FOXP1,CNP,NTRK2,TACC2,KDM6B,PCDH9,FGF13,DMD |
| **GO:0048699** | Generation of neurons | 0.0158 | FRY,KIT,PRKCH,PTPRO,EPHB1,NCS1,DPYSL5,TENM4,TNFRSF21,PTPRM,TENM3,TUBB3,NTRK2,ABCC8,TIAM1,FGF13,IMPACT,DMD |

| | | | |
|---|---|---|---|
| **GO:0007417** | Central nervous system development | 0.0261 | FYN,EPHB1,FOXP1,TNFRSF21,CNP,NTRK2,TACC2,SOX6,ZBTB16,KDM6B,PCDH9,FGF13,DMD |
| **GO:0040011** | Locomotion | 0.0292 | FYN,CD47,MYLK,KIT,SLC9A3R1,PTPRO,EPHB1,PTPRM,TUBB3,NTRK2,SEMA4A,ABCC8,TIAM1,PLEKHG5,FGF13 |
| **GO:0042063** | Gliogenesis | 0.0292 | PLP1,PRKCH,TNFRSF21,SOX6,ABCC8,TIAM1,PARD3 |
| **GO:0021537** | Telencephalon development | 0.0314 | FOXP1,NTRK2,TACC2,KDM6B,FGF13,DMD |
| **GO:0048812** | Neuron projection morphogenesis | 0.0314 | PTPRO,EPHB1,PTPRM,TUBB3,NTRK2,TIAM1,FGF13,IMPACT,DMD |
| **GO:0048858** | Cell projection morphogenesis | 0.0314 | PTPRO,EPHB1,PTPRM,TUBB3,NTRK2,TIAM1,FGF13,IMPACT,DMD |
| **GO:0120039** | Plasma membrane bounded cell projection morphogenesis | 0.0314 | PTPRO,EPHB1,PTPRM,TUBB3,NTRK2,TIAM1,FGF13,IMPACT,DMD |
| **GO:0051960** | Regulation of nervous system development | 0.0392 | KIT,PRKCH,EPHB1,NCS1,TNFRSF21,NTRK2,ABCC8,TIAM1,PARD3,FGF13,IMPACT,DMD |
| **GO:0007409** | Axonogenesis | 0.0409 | PTPRO,EPHB1,PTPRM,TUBB3,NTRK2,TIAM1,FGF13 |

| | | | |
|---|---|---|---|
| **GO:0098885** | Modification of postsynaptic actin cytoskeleton | 0.0409 | MYH10,TIAM1 |
| **GO:0099010** | Modification of postsynaptic structure | 0.0409 | MYH10,TIAM1 |
| **GO:0048666** | Neuron development | 0.0413 | FRY,PTPRO,EPHB1,NCS1,TENM4,PTPRM,TENM3,TUBB3,NTRK2,TIAM1,FGF13,IMPACT,DMD |
| **GO:0021987** | Cerebral cortex development | 0.0458 | NTRK2,TACC2,FGF13,DMD |
| **GO:0007612** | Learning | 0.0487 | KIT,NTRK2,ABCC8,FGF13 |
| **GO:0120036** | Plasma membrane bounded cell projection organization | 0.0487 | FRY,P2RX7,KIT,PTPRO,EPHB1,NCS1,PTPRM,TUBB3,CDHR5,NTRK2,TMEM231,TIAM1,FGF13,IMPACT,DMD |
| **Protein Phosphorylation and signal transduction** | | | |
| **GO:0043087** | Regulation of gtpase activity | 0.00279 | SGSM1,RGS9,SYDE1,ARHGAP21,TBC1D8,TBC1D10B,NTRK2,TIAM1,SYDE2 |
| **GO:0006468** | Protein phosphorylation | 0.00402 | FYN,P2RX7,MAP3K7CL,KIT,SLC9A3R1,PRKCH,EPHB1,MAP3K6,CDK6,EDNRB,KSR1,MELK,JTB,MERTK,NTRK2,LIMK2,TIAM1,PPP1R1C,ACVRL1,PARD3,IMPACT,SH3BP5,BMP7 |
| **GO:0065009** | Regulation of molecular function | 0.00597 | SGSM1,MAP3K7CL,TGFBR3,DLG3,SLC9A3R1,RGS9,PRKCH,SYDE1,ARHGAP21,MAP3K6,WRAP53,TBC1D8,MIDN,JTB,TBC1D10B,NEDD4L,NTRK2,ABCC8,TIAM1,TOR1AIP2,PPP1R1C,SYDE2,JSRP1,SH3BP5 |

| | | | |
|---|---|---|---|
| **GO:0016310** | Phosphorylation | 0.00698 | FYN,P2RX7,MAP3K7CL,KIT,SLC9A3R1,PRKCH,EPHB1,MAP3K6,CDK6,EDNRB,KSR1,MELK,MIDN,JTB,MERTK,NTRK2,LIMK2,TIAM1,PPP1R1C,ACVRL1,PARD3,IMPACT,SH3BP5,BMP7 |
| **GO:0019220** | Regulation of phosphate metabolic process | 0.0121 | P2RX7,MAP3K7CL,DLG3,SLC9A3R1,EPHB1,MAP3K6,EDNRB,KSR1,MIDN,JTB,NTRK2,PTH1R,RUNDC3A,TIAM1,PPP1R1C,PARD3,IMPACT,SH3BP5,BMP7 |
| **GO:0051174** | Regulation of phosphorus metabolic process | 0.0121 | P2RX7,MAP3K7CL,DLG3,SLC9A3R1,EPHB1,MAP3K6,EDNRB,KSR1,MIDN,JTB,NTRK2,PTH1R,RUNDC3A,TIAM1,PPP1R1C,PARD3,IMPACT,SH3BP5,BMP7 |
| **GO:0050790** | Regulation of catalytic activity | 0.0141 | SGSM1,MAP3K7CL,DLG3,SLC9A3R1,RGS9,SYDE1,ARHGAP21,MAP3K6,WRAP53,TBC1D8,MIDN,JTB,TBC1D10B,NTRK2,TIAM1,TOR1AIP2,PPP1R1C,SYDE2,SH3BP5 |
| **GO:0042325** | Regulation of phosphorylation | 0.0261 | P2RX7,MAP3K7CL,SLC9A3R1,EPHB1,MAP3K6,EDNRB,KSR1,MIDN,JTB,NTRK2,TIAM1,PPP1R1C,PARD3,IMPACT,SH3BP5,BMP7 |
| **GO:0043547** | Positive regulation of gtpase activity | 0.0303 | SGSM1,RGS9,ARHGAP21,TBC1D8,TBC1D10B,TIAM1 |
| **GO:0001932** | Regulation of protein phosphorylation | 0.0314 | P2RX7,MAP3K7CL,SLC9A3R1,EPHB1,MAP3K6,EDNRB,KSR1,JTB,NTRK2,TIAM1,PPP1R1C,PARD3,IMPACT,SH3BP5,BMP7 |
| **GO:0019934** | Cgmp-mediated signaling | 0.0314 | PDE9A,EDNRB |
| **GO:0051968** | Positive regulation of synaptic transmission, glutamatergic | 0.0314 | NTRK2,GLUL,IQSEC2 |

| | | | |
|---|---|---|---|
| **GO:0090630** | Activation of gtpase activity | 0.0314 | SGSM1,TBC1D8,TBC1D10B,TIAM1 |
| **GO:0051336** | Regulation of hydrolase activity | 0.0365 | SGSM1,DLG3,RGS9,SYDE1,ARHGAP21,TBC1D8,TBC1D10B,NTRK2,TIAM1,TOR1AIP2,SYDE2 |
| **GO:0051345** | Positive regulation of hydrolase activity | 0.0373 | SGSM1,RGS9,ARHGAP21,TBC1D8,TBC1D10B,NTRK2,TIAM1,TOR1AIP2 |
| **GO:0010646** | Regulation of cell communication | 0.0428 | BTBD9,FYN,MAP3K7CL,TGFBR3,KIT,SLC9A3R1,RGS9,PRKCH,PTPRO,EPHB1,SLC24A2,MAP3K6,KSR1,MIDN,NTRK2,ABCC8,TIAM1,JSRP1,SLC25A5,GLUL,BMP7,IQSEC2 |
| **GO:0031399** | Regulation of protein modification process | 0.0428 | P2RX7,MAP3K7CL,SLC9A3R1,EPHB1,MAP3K6,COMMD1,EDNRB,KSR1,JTB,NTRK2,TIAM1,PPP1R1C,PARD3,IMPACT,SH3BP5,BMP7 |
| **GO:1901615** | Organic hydroxy compound metabolic process | 0.0428 | KIT,ITPKB,CYP51,TPH1,CYP24A1,PTH1R,PCK1 |
| **GO:0035556** | Intracellular signal transduction | 0.0444 | FYN,PDE9A,MAP3K7CL,ARHGAP31,SLC9A3R1,PRKCH,EPHB1,NCS1,MAP3K6,SRPK2,EDNRB,KSR1,MELK,NTRK2,LIMK2,PTH1R,TIAM1,PPP1R1C,JSRP1,SH3BP5,BMP7 |
| **GO:0023051** | Regulation of signaling | 0.0458 | BTBD9,FYN,MAP3K7CL,TGFBR3,KIT,SLC9A3R1,RGS9,PRKCH,PTPRO,EPHB1,SLC24A2,MAP3K6,KSR1,MIDN,NTRK2,ABCC8,TIAM1,JSRP1,SLC25A5,GLUL,BMP7,IQSEC2 |
| **GO:0043085** | Positive regulation of catalytic activity | 0.0458 | SGSM1,MAP3K7CL,RGS9,ARHGAP21,MAP3K6,WRAP53,TBC1D8,JTB,TBC1D10B,NTRK2,TIAM1,TOR1AIP2 |

| | | | |
|---|---|---|---|
| **GO:0044093** | Positive regulation of molecular function | 0.0458 | SGSM1,MAP3K7CL,RGS9,PRKCH,ARHGAP21,MAP3K6,WRAP53,TBC1D8,JTB,TBC1D10B,NTRK2,ABCC8,TIAM1,TOR1AIP2 |
| **GO:0048015** | Phosphatidylinositol-mediated signaling | 0.0471 | FYN,SLC9A3R1,NCS1,NTRK2 |
| **GO:0048017** | Inositol lipid-mediated signaling | 0.0471 | FYN,SLC9A3R1,NCS1,NTRK2 |
| **Cellular Differentiation and morphogenesis** | | | |
| **GO:0048468** | Cell development | 0.00402 | FRY,KIT,PRKCH,PTPRO,EPHB1,NCS1,FOXP1,TENM4,TNFRSF21,PTPRM,TENM3,SGCG,HIST1H2BA,TUBB3,NTRK2,LIMK2,ABCC8,TIAM1,PARD3,FGF13,IMPACT,DMD,BMP7 |
| **GO:0051128** | Regulation of cellular component organization | 0.00597 | SGSM1,CIT,P2RX7,CD47,KIT,PRKCH,RBBP7,EPHB1,NCS1,FOXP1,TBC1D8,SYNPO2,TBC1D10B,NEDD4L,CDHR5,ANK1,NTRK2,ABCC8,TIAM1,SLC25A5,FGF13,IMPACT,DMD |
| **GO:0000902** | Cell morphogenesis | 0.0155 | FRY,PTPRO,EPHB1,TENM4,PTPRM,TENM3,TUBB3,NTRK2,TIAM1,FGF13,IMPACT,DMD |
| **GO:0032989** | Cellular component morphogenesis | 0.0275 | FRY,PTPRO,EPHB1,TENM4,PTPRM,TENM3,TUBB3,NTRK2,TIAM1,FGF13,IMPACT,DMD |
| **GO:0008283** | Cell proliferation | 0.0302 | FYN,KIT,PRKCH,EPHB1,KLF5,CDK6,EDNRB,TNFRSF21,PTPRM,POLR3G,NTRK2,TACC2,ABCC8,TNFRSF25,SLC25A5,FGF13,DMD,GLUL |
| **GO:0032990** | Cell part morphogenesis | 0.0314 | PTPRO,EPHB1,PTPRM,TUBB3,NTRK2,TIAM1,FGF13,IMPACT,DMD |

| | | | |
|---|---|---|---|
| **GO:0048103** | Somatic stem cell division | 0.0314 | KIT,FGF13 |
| **GO:0045596** | Negative regulation of cell differentiation | 0.0365 | EPHB1,KLF5,FOXP1,NOCT,ABCC8,ZBTB16,FGF13,DMD |
| **GO:0060341** | Regulation of cellular localization | 0.0365 | COMMD1,KCNAB2,EPHB2,NTRK2,TIAM1,PARD3,MLC1,JSRP1,SLC25A5 |
| **GO:0048870** | Cell motility | 0.0381 | FYN,CD47,MYLK,KIT,SLC9A3R1,PTPRO,EPHB1,PTPRM,NTRK2,ABCC8,TIAM1,PLEKHG5,FGF13 |
| **GO:0051674** | Localization of cell | 0.0381 | FYN,CD47,MYLK,KIT,SLC9A3R1,PTPRO,EPHB1,PTPRM,NTRK2,ABCC8,TIAM1,PLEKHG5,FGF13 |
| **GO:0007043** | Cell-cell junction assembly | 0.0428 | PRKCH,PTPRO,PARD3 |
| **GO:0045595** | Regulation of cell differentiation | 0.0428 | KIT,PRKCH,EPHB1,NCS1,KLF5,FOXP1,NOCT,TNFRSF21,NTRK2,ABCC8,TIAM1,ZBTB16,FGF13,IMPACT,DMD |
| **GO:0016477** | Cell migration | 0.0458 | FYN,CD47,MYLK,KIT,PTPRO,EPHB1,PTPRM,NTRK2,ABCC8,TIAM1,PLEKHG5,FGF13 |
| **Organism and system development** | | | |
| **GO:0051239** | Regulation of multicellular organismal process | 0.0096 | SRGN,PDE9A,P2RX7,KIT,PRKCH,PTPRO,EPHB1,NCS1,KLF5,FOXP1,NOCT,EDNRB,TNFRSF21,TPH1,PTPRM,HSPD1,NTRK2,ABCC8,TIAM1,ZBTB16,PARD3,FGF13,IMPACT,DMD |

| | | | |
|---|---|---|---|
| **GO:0051093** | Negative regulation of developmental process | 0.00988 | SRGN,EPHB1,KLF5,FOXP1,NOCT,TNFRSF21,PTPRM,ABCC8,ZBTB16,FGF13,DMD |
| **GO:0051241** | Negative regulation of multicellular organismal process | 0.00988 | SRGN,PTPRO,EPHB1,KLF5,FOXP1,NOCT,TNFRSF21,TPH1,PTPRM,ABCC8,ZBTB16,FGF13,DMD |
| **GO:0003012** | Muscle system process | 0.0149 | PDE9A,MYLK,KIT,FOXP1,NEDD4L,ABCC8,TIAM1,JSRP1,DMD |
| **GO:2000026** | Regulation of multicellular organismal development | 0.0154 | SRGN,KIT,PRKCH,EPHB1,NCS1,KLF5,FOXP1,NOCT,TNFRSF21,PTPRM,NTRK2,ABCC8,TIAM1,ZBTB16,PARD3,FGF13,IMPACT,DMD |
| **GO:0030279** | Negative regulation of ossification | 0.0286 | SRGN,NOCT,TPH1 |
| **GO:0044057** | Regulation of system process | 0.0372 | PDE9A,KIT,PTPRO,FOXP1,EDNRB,TNFRSF21,ABCC8,PARD3,DMD |
| **GO:0030278** | Regulation of ossification | 0.0381 | SRGN,NOCT,TPH1,ZBTB16 |
| **GO:0021543** | Pallium development | 0.0428 | NTRK2,TACC2,KDM6B,FGF13,DMD |
| **GO:1901343** | Negative regulation of vasculature development | 0.0458 | KLF5,PTPRM,ABCC8 |

| | | | |
|---|---|---|---|
| **GO:0048513** | Animal organ development | 0.0475 | SRGN,TGFBR3,KIT,PRKCH,PTPRO,EPHB1,KLF5,FOXP1,CCDC182,PTPRM,TGFBI,SGCG,IGF2R,CNP,NTRK2,CORO1A,TACC2,STC2,ZBTB16,NHS,KDM6B,PCDH9,FGF13,DMD |
| **GO:0050793** | Regulation of developmental process | 0.0484 | SRGN,KIT,PRKCH,EPHB1,NCS1,KLF5,FOXP1,NOCT,TNFRSF21,PTPRM,NTRK2,ABCC8,TIAM1,ZBTB16,PARD3,FGF13,IMPACT,DMD |
| **Stimulus Response** | | | |
| **GO:0071332** | Cellular response to fructose stimulus | 0.0234 | SLC2A5,PCK1 |
| **GO:0032496** | Response to lipopolysaccharide | 0.0304 | NOCT,EDNRB,TNFRSF21,HSPD1,CNP,ABCC8,TNFRSF25,PCK1,MGST2 |
| **GO:0002237** | Response to molecule of bacterial origin | 0.0314 | NOCT,EDNRB,TNFRSF21,HSPD1,CNP,ABCC8,TNFRSF25,PCK1,MGST2 |
| **GO:0033993** | Response to lipid | 0.0413 | RBBP7,FOXP1,NOCT,EDNRB,TNFRSF21,HSPD1,IGF2R,CNP,NTRK2,STC2,ABCC8,TNFRSF25,PAQR6,PCK1,MLC1,BMP7,MGST2 |
| **GO:0009750** | Response to fructose | 0.0477 | SLC2A5,PCK1 |

**Supplementary Table 7: KEGG Pathways enriched by the genes identified from DMR analysis at P7 along with their FDR values and list of enriched genes.**

| ID | Term | FDR | Genes |
|---|---|---|---|
| **KEGG:00604** | Glycosphingolipid biosynthesis - ganglio series | 9.14E-03 | ST8SIA1, ST8SIA5, ST6GALNAC3 |
| **KEGG:04010** | MAPK signaling pathway | 4.98E-03 | MAPK10, RPS6KA6, MAP3K14, RPS6KA3, CACNA1C, CACNA1F, RASGRF1, NFATC1, MAP3K4, DUSP9, FGF16 |
| **KEGG:04020** | Calcium signaling pathway | 5.30E-03 | PLCD3, CACNA1C, SLC8A1, CACNA1F, SPHK1, HTR2C, PLCB4, CAMK2B, ATP2B3 |
| **KEGG:04022** | cGMP-PKG signaling pathway | 8.64E-03 | CACNA1C, SLC8A1, CACNA1F, NFATC2, NFATC1, MRVI1, PLCB4,ATP2B3 |
| **KEGG:04024** | cAMP signaling pathway | 2.73E-03 | MAPK10, CACNA1C, GRIA3, ABCC4, CACNA1F, GRIN3B, NFATC1, TIAM1, CAMK2B, ATP2B3 |
| **KEGG:04261** | Adrenergic signaling in cardiomyocytes | 4.69E-02 | CACNA1C, SLC8A1, CACNA1F, PLCB4, CAMK2B, ATP2B3 |
| **KEGG:04310** | Wnt signaling pathway | 1.55E-02 | MAPK10, FZD6, NFATC2, CTNNBIP1, NFATC1, PLCB4, CAMK2B |
| **KEGG:04360** | Axon guidance | 6.80E-04 | FYN, NTN1, PAK3, EFNB1, NFATC2, RGS3, SLIT1, SEMA6B,CAMK2B, BMP7, PLXNA3 |
| **KEGG:04530** | Tight junction | 2.69E-02 | RUNX1, MAPK10, RAP2C, DLG3, TIAM1, ARHGEF18, EPB41L4B |
| **KEGG:04713** | Circadian entrainment | 3.82E-02 | CACNA1C, GRIA3, PER3, PLCB4, CAMK2B |
| **KEGG:04720** | Long-term potentiation | 9.14E-03 | RPS6KA6, RPS6KA3, CACNA1C, PLCB4, CAMK2B |
| **KEGG:04722** | Neurotrophin signaling pathway | 2.69E-02 | MAPK10, RPS6KA6, RPS6KA3, MAGED1, BEX3, CAMK2B |
| **KEGG:04723** | Retrograde endocannabinoid signaling | 2.15E-02 | MAPK10, CACNA1C, GRIA3, NDUFB11, CACNA1F, PLCB4, GABRE |
| **KEGG:04912** | GnRH signaling pathway | 8.64E-03 | MAPK10, CACNA1C, CACNA1F, MAP3K4, PLCB4, CAMK2B |
| **KEGG:04919** | Thyroid hormone signaling pathway | 2.73E-03 | NOTCH4, SLC16A2, PLCD3, MED14, MED12, BMP4, MED16, PLCB4 |
| **KEGG:04921** | Oxytocin signaling pathway | 6.94E-03 | CDKN1A, CACNA1C, CACNA1F, NFATC2, EEF2K, NFATC1, PLCB4, CAMK2B |
| **KEGG:05010** | Alzheimer's disease | 3.93E-02 | APP, CACNA1C, NDUFB11, CACNA1F, TAF3, UQCRC1, PLCB4 |
| **KEGG:05031** | Amphetamine addiction | 4.69E-02 | CACNA1C, GRIA3, GRIN3B, CAMK2B |

**Supplementary Table 8. KEGG Pathways enriched by the genes identified from DMR analysis at P90 along with their FDR values and list of enriched genes.**

| ID | Term | FDR | Genes |
|---|---|---|---|
| **KEGG:00230** | Purine metabolism | 2.25E-02 | PDE9A, ADCY8, GUCY2C, GUCY2E, POLR3G, PDE8A, POLD1, GUCY1A2 |
| **KEGG:04010** | MAPK signaling pathway | 4.47E-02 | CACNA1C, MAP3K6, MECOM, CACNA1D, NFATC1, NTRK2, NFKB1, CACNA2D3, FGF13 |
| **KEGG:04014** | Ras signaling pathway | 2.25E-03 | FLT1, KIT, RALB, FLT4, KSR1, IGF1R, RASA3, NTRK2, GNG7, TIAM1, NFKB1, FGF13 |
| **KEGG:04015** | Rap1 signaling pathway | 2.46E-03 | FLT1, KIT, RALB, FLT4, ADCY8, IGF1R, TIAM1, MAGI1, PARD3, FGF13, RAPGEF3 |
| **KEGG:04020** | Calcium signaling pathway | 2.93E-03 | NOS1, P2RX7, MYLK, ITPKB, PHKA1, ADCY8, CACNA1C, EDNRB, CACNA1D, SLC25A5 |
| **KEGG:04022** | cGMP-PKG signaling pathway | 5.38E-03 | ATF6B, MYLK, ADCY8, CACNA1C, EDNRB, CACNA1D, NFATC1, GUCY1A2, SLC25A5 |
| **KEGG:04024** | cAMP signaling pathway | 1.21E-02 | ADCY8, CACNA1C, GABBR2, CACNA1D, NFATC1, VAV3, TIAM1, NFKB1, RAPGEF3 |
| **KEGG:04151** | PI3K-Akt signaling pathway | 2.93E-03 | ATF6B, FLT1, COL6A2, KIT, FLT4, CDK6, IGF1R, COL4A3, NTRK2, VWF, GNG7, NFKB1, PCK1, FGF13 |
| **KEGG:04270** | Vascular smooth muscle contraction | 2.84E-02 | MYLK, PRKCH, ADCY8, CACNA1C, CACNA1D, GUCY1A2 |
| **KEGG:04360** | Axon guidance | 9.09E-03 | FYN, EPHB1, DPYSL5, EPHB2, LIMK2, SEMA4A, RGS3, PARD3, BMP7 |
| **KEGG:04510** | Focal adhesion | 1.41E-02 | FYN, FLT1, COL6A2, MYLK, FLT4, IGF1R, COL4A3, VWF,VAV3 |
| **KEGG:04512** | ECM-receptor interaction | 2.84E-02 | COL6A2, CD47, COL4A3, VWF, CD36 |
| **KEGG:04530** | Tight junction | 1.52E-03 | DLG3, MYH10, SLC9A3R1, CLDN11, CACNA1D, TJP3, TIAM1, MAGI1, ARHGEF18, PARD3, CGNL1 |

| | | | |
|---|---|---|---|
| **KEGG:04713** | Circadian entrainment | 1.41E-02 | NOS1, ADCY8, CACNA1C, CACNA1D, GNG7, GUCY1A2 |
| **KEGG:04724** | Glutamatergic synapse | 9.09E-03 | ADCY8, SLC38A1, CACNA1C, CACNA1D, SLC1A3, GNG7, GLUL |
| **KEGG:04727** | GABAergic synapse | 1.53E-03 | ADCY8, SLC38A1, CACNA1C, GABBR2, SLC6A13, CACNA1D, GNG7, GLUL |
| **KEGG:04730** | Long-term depression | 4.67E-02 | NOS1, GNAZ, IGF1R, GUCY1A2 |
| **KEGG:04911** | Insulin secretion | 1.19E-02 | ATF6B, ADCY8, CACNA1C, CACNA1D, KCNN3, ABCC8 |
| **KEGG:04921** | Oxytocin signaling pathway | 2.84E-02 | MYLK, ADCY8, CACNA1C, CACNA1D, NFATC1, GUCY1A2, CACNA2D3 |
| **KEGG:04925** | Aldosterone synthesis and secretion | 2.84E-02 | ATF6B, ADCY8, CACNA1C, CACNA1D, DAGLA |
| **KEGG:04926** | Relaxin signaling pathway | 2.01E-02 | ATF6B, NOS1, ADCY8, EDNRB, COL4A3, GNG7, NFKB1 |
| **KEGG:04964** | Proximal tubule bicarbonate reclamation | 2.79E-02 | SLC4A4, SLC9A3, PCK1 |
| **KEGG:05165** | Human papillomavirus infection | 2.84E-02 | COL6A2, DLG3, SLC9A3R1, TRAF3, CDK6, COL4A3, VWF, MAGI1, NFKB1, PARD3, AXIN2 |
| **KEGG:05200** | Pathways in cancer | 2.25E-03 | KIT, RALB, ADCY8, TRAF3, CDK6, EDNRB, MECOM, IGF1R, COL4A3, GNG7, PLEKHG5, NFKB1, ZBTB16, FGF13, TRAF1, AXIN2 |
| **KEGG:05202** | Transcriptional misregulation in cancer | 2.74E-02 | FLT1, AFF1, IGF1R, RUNX2, NFKB1, ZBTB16, TRAF1, NCOR1 |
| **KEGG:05410** | Hypertrophic cardiomyopathy (HCM) | 2.95E-02 | CACNA1C, CACNA1D, SGCG, CACNA2D3, EMD |
| **KEGG:05412** | Arrhythmogenic right ventricular cardiomyopathy (ARVC) | 1.53E-03 | CACNA1C, CACNA1D, SGCG, CDH2, ACTN2, CACNA2D3, EMD |
| **KEGG:05414** | Dilated cardiomyopathy (DCM) | 1.19E-02 | ADCY8, CACNA1C, CACNA1D, SGCG, CACNA2D3, EMD |
